# Oxidative Stress-Induced Microglial CD22 Upregulation Impairs Phagocytosis and Exacerbates Huntington’s Disease

**DOI:** 10.64898/2026.03.11.710967

**Authors:** Yan Hua Lee, Jian Jing Siew, Chia-Wei Lee, Hui-Mei Chen, Ya-Ting Lu, Deepa Sridharan, Po-Chun Jimmy Huang, Hua-Chien Chang, Shih-Yun Guu, Pao-Yuan Wang, Yi-Fu Wang, Suh-Yuen Liang, Kay-Hooi Khoo, Shih-Yu Chen, Takashi Angata, Yijuang Chern

## Abstract

**Background:** Huntington’s disease (HD) is a neurodegenerative disorder caused by an abnormal polyglutamine expansion in mutant huntingtin (mHTT) and is characterized by movement dysfunction and neuronal loss. Siglecs, a family of sialic acid-binding proteins, are expressed on brain microglia and implicated in Alzheimer’s disease. Sialic acids are abundant in mammalian brains and cap the termini of the glycocalyx of various brain cells. Alterations in sialoglycans or Siglecs may affect interactions between microglia and other brain cells. However, the roles of Siglecs in HD have not been investigated.

**Methods:** We profiled Siglecs in postmortem caudate nucleus samples from HD subjects and in a mouse model of HD (R6/2) using RT-qPCR and mass cytometry analyses. CD22 functions in microglia were evaluated using a microglial cell line (BV2) and primary microglia. Native ligands for microglial CD22 were assessed via glycomic profiling and flow cytometry. Regulation of CD22 ligands in astrocytes was investigated in an astrocytic cell line (C8-D1A) and primary astrocytes. The role of CD22 in HD was examined by genetic deletion in HD mice, followed by behavioral analyses and pathological evaluation with immunofluorescence staining and MRI.

**Results:** Upregulation of CD22 in microglia, observed in the brains of patients and mice with HD, impairs microglial phagocytosis via ITIM-ITAM signaling crosstalk. This CD22 upregulation was driven by chronic oxidative stress, as antioxidant treatment (N-acetylcysteine) markedly normalized CD22 levels. CD22 ligand, α2,6-sialylated-6-sulfo-LacNAc, primarily expressed by astrocytes, was significantly reduced in HD mice. mHTT, but not wild-type HTT, suppressed ligand synthesis in astrocytes under elevated oxidative stress, allowing more CD22 on the microglial surface to inhibit phagocytosis. Treatment with a neutralizing antibody or ligand-enriched extracellular vesicles depleted surface CD22 and restored the phagocytic function of microglia. Genetic deletion of CD22 in HD mice improved rotarod performance, reduced mHTT inclusion burden, increased Darpp32 expression, and alleviated brain atrophy, supporting the concept that CD22-mediated inhibition of microglial phagocytosis contributes to HD pathogenesis.

**Conclusion:** Our findings suggest that CD22 acts as a checkpoint-like regulator that restrains microglial phagocytosis and contributes to HD progression when astrocyte-microglia communication is impaired, thereby highlighting CD22 as a promising therapeutic target.

## Background

Mammalian cells are coated with a diverse array of glycans, with sialic acids frequently capping the termini of these branches, serving as critical molecular determinants [1]. Siglecs (sialic acid-binding immunoglobulin-type lectins) are a family of immunomodulatory receptors expressed on immune cells that recognize sialic acids through their amino-terminal V-set immunoglobulin domain [2, 3]. The evolutionarily conserved Siglecs, including Siglec-1, CD22 (Siglec-2), MAG (Siglec-4), and Siglec-15, have well-defined orthologs in humans, mice, and other mammalian species [3]. Meanwhile, the CD33-related Siglecs show species-to-species variation, such that Siglec-E, Siglec-F, and Siglec-G in mice are functional counterparts of Siglec-7/9, Siglec-8, and Siglec-10 in humans, respectively [3]. Most Siglecs possess one or more consensus immunoreceptor tyrosine-based inhibitory motifs (ITIMs) in their cytoplasmic domains, whereas some harbor a positively charged amino acid in the transmembrane domain that associates with membrane adaptor proteins containing immunoreceptor tyrosine-based activation motifs (ITAMs) [2, 3]. Numerous studies have demonstrated that interactions between sialic acids and Siglecs may facilitate communication between immune cells and their counterparts, leading to the activation or deactivation of immune cell functions, including activation thresholds, tolerance, and phagocytosis [2–5].

Huntington’s disease (HD) is a progressive neurodegenerative disease caused by a CAG trinucleotide repeat expansion in the HTT gene. This mutation leads to the production of a toxic polyglutamine-expanded form of huntingtin (mHTT), leading to selective neuronal loss and striatal atrophy, which manifests clinically as uncontrollable body movements [6]. Despite extensive investigation of HD pathogenesis over the past two decades, no effective therapeutic agent capable of delaying HD progression is currently available. While neuronal insoluble mHTT aggregates are central to disease pathology, emerging studies indicate that glial cells, including microglia and astrocytes, also contribute to HD pathogenesis [7–12]. Microglia have been extensively studied in Alzheimer’s disease (AD), where genome-wide association studies (GWAS) have identified several risk genes linked to microglia [13–15]. In contrast, GWAS in HD are limited and have yet to identify strong microglia-specific genetic associations. Nevertheless, our previous work demonstrated that suppression of microglia-mediated neuroinflammation ameliorates HD-like symptoms in a mouse model of HD, supporting the importance of microglia in HD pathogenesis [10].

In the present study, we investigated whether sialic acid-Siglec interactions serve as glyco-immune checkpoints regulating microglial functions, motivated by the high abundance of sialic acids in the mammalian brain. Alterations in sialoglycans or Siglecs may cause microglial dysfunction and/or disrupted communication between microglia and other brain cells, potentially contributing to HD pathogenesis. Our findings in both patients and mice with HD reveal significant upregulation of CD22, which suppresses phagocytosis via ITIM-ITAM receptor interactions in microglia. This is particularly intriguing because CD22 has recently been identified as a modifier of age-associated microglia and is highly conserved between mice and humans [16]. We further demonstrate that microglial phagocytic function can be restored through multiple approaches, including depletion of surface CD22 with a neutralizing antibody, astrocyte-derived extracellular vesicles (EVs), or genetic deletion of CD22. Genetic CD22 deletion improved motor performance, reduced mHTT aggregates, restored striatal Darpp32 expression, and enhanced microglial clearance capacity in HD mice. Together, our findings reveal that CD22 modulates microglial function independently of sialic acid binding, while ligand interactions may potentially reshape astrocyte–microglia communication, highlighting the therapeutic relevance of CD22.

## Methods

### Human Brain Sections

Postmortem HD and non-HD human brain tissue specimens were obtained from the NIH NeuroBioBank (USA) (Table S1).

### Animals

R6/2 (B6CBA-Tg(HDexon1)62Gpb/1J) mice were purchased from The Jackson Laboratory (Bar Harbor, ME, USA) [17] and maintained in the animal facility of the Institute of Biomedical Sciences (IBMS), at Academia Sinica (Taipei, Taiwan) under standard conditions. Offspring genotypes were verified by amplifying the human *mHTT* gene from genomic DNA isolated from mouse tails using the following primers: 5’-CCGCTCAGGTTCTGCTTTTA-3’ and 5’-GGCTGAGGAAGCTGAGGAG-3’. Littermate wild-type (WT) mice were used as controls in subsequent experiments.

### Behavioral Analyses

Body weight was measured weekly. Motor coordination was assessed using a rotarod apparatus (UGO BASILE, Comerio, Italy) set at a constant speed of 12 rpm for 120 seconds, with three trials performed per mouse. Limb clasping behavior was evaluated by suspending mice by the tail for 30 seconds. All animal experimental procedures were conducted according to the guidelines established by the Institutional Animal Care and Use Committee (IACUC) of the IBMS at Academia Sinica.

### Plasmid Construction

The pLV-hPGK−mCD22^WT^-T2A-GFP−miR9T, pLV-hPGK−mCD22^R130A^-T2A-GFP−miR9T and pLV- hPGK−Chst2-T2A-GFP−miR9T expression constructs were generated from pCDNA3.1(-)−mCD22^WT^, pCDNA3.1(-)−mCD22^R130A^ and pCDNA3.1(-)−Chst2 and subsequently subcloned into the pLV.PGK.GFP.miR9T vector obtained from Dr. Johan Jakobsson’s lab [18].

### Cell Culture and Transfection

HEK293T cells were cultured in Dulbecco’s modified Eagle’s medium (DMEM; HyClone™, Cytiva, Marlborough, MA, USA) supplemented with 10% heat-inactivated fetal bovine serum (FBS; HyClone™, Cytiva, Marlborough, MA, USA) and penicillin/streptomycin. BV2 and C8-D1A cells were cultured in Dulbecco’s modified Eagle’s medium (DMEM; HyClone™, Cytiva, Marlborough, MA, USA) supplemented with 10% heat-inactivated fetal bovine serum (FBS; HyClone™, Cytiva, Marlborough, MA, USA), penicillin/streptomycin, and GlutaMAX (Thermo Fisher Scientific, Waltham, MA, USA). For C8-D1A cells, 1 mM sodium pyruvate (Thermo Fisher Scientific, Waltham, MA, USA) was additionally added to the medium. Transfection was performed using T-Pro NTR II (Ji-Feng Biotechnology, Taiwan) following the manufacturer’s protocol.

### Lentivirus Preparation and Infection

Lentiviruses were prepared following standard procedures provided by the National RNAi Core Facility, Academia Sinica. Briefly, HEK293T cells were transfected with the empty vector (pLV-hPGK-miR9T-GFP) or vectors containing the target genes (CD22^WT^-T2A-GFP, CD22^R130A^-T2A-GFP, HTT_Ex1_Q_19_-T2A-GFP, HTT_Ex1_Q_89_-T2A-GFP or Chst2-T2A-GFP), along with the packaging vector (pCMV.dR8.74.PpsAX2) and the envelope vector (pMD2.G), using T-Pro NTR II (Ji-Feng Biotechnology, Taiwan). Sixteen hours after transfection, the medium was replaced with high-serum medium (DMEM supplemented with 30% heat-inactivated FBS). The supernatant containing viral particles was harvested at 24, 48, and 72 hours after medium replacement. Virus titers were determined by transducing cells with serially diluted virus and assessing the percentage of GFP-positive cells using flow cytometry. BV2 and C8-D1A cells were transduced at a multiplicity of infection (MOI) of 30. Stable clones of the transduced cells were subsequently sorted based on GFP expression using FACSAria IIIu cell sorter (Becton, Dickinson and Company; East Rutherford, NJ, USA).

### Primary Microglia and Astrocytes

Primary microglia were harvested as described in our previous study [10], and primary astrocytes were prepared according to protocols detailed elsewhere [10, 19]. All cells were maintained in a humidified incubator at 37 °C with 10% CO_2_.

### Phagocytosis

The phagocytosis assay was performed as previously described [16]. BV2 cells and primary microglia were seeded onto coverslips in a 24-well plate and incubated overnight before feeding. A single vial of pHrodo™ Red Zymosan A BioParticles (Invitrogen, Carlsbad, CA, USA) was resuspended in pH-neutral phosphate-buffered saline (PBS, pH 7.4) by adding 2 mL of PBS to create a 0.5 mg/mL suspension. After the indicated treatment, the cells were fed with pHrodo™ Red Zymosan A BioParticles at a ratio of 15 particles per cell, and then incubated at 37 °C for 60 minutes. Following PBS washes, cells were fixed in 4% paraformaldehyde for 15 minutes at room temperature. Nuclei were stained with Hoechst and mounted with EverBrite™ Hardset Mounting Medium (BIOTREND Chemikalien GmbH, Köln, Germany) before imaging with a Zeiss LSM 700 Stage confocal microscope. For each replicate, each data point represents the average of ten images (fields of view) acquired from one coverslip.

### Total RNA Isolation and Quantitative Real-Time PCR (RT-qPCR)

Total RNA was extracted from cells and brain tissues using the GENEzol^TM^ TriRNA Pure Kit (Geneaid Biotech, Taiwan) according to the manufacturer’s protocol. The isolated RNAs were reverse transcribed into cDNA using Superscript III (Invitrogen, Carlsbad, CA, USA). Quantitative real-time PCR (RT-qPCR) was performed using the SYBR® Green PCR Master Mix (Thermo Fisher Scientific, Waltham, MA, USA) and detected using the LightCycler® 480 System (Roche Diagnostics, Basel, Switzerland). Relative gene expression levels were determined by normalization with *Gapdh* and *18S* as the reference genes for mouse and human samples, respectively. All primers used for RT-qPCR are listed in Tables S3 and S4.

### Flow Cytometry

The indicated mouse brain areas were dissected and dissociated with the neural tissue dissociation kit (P) (Miltenyi Biotec; 130-092-628) following the manufacturer’s protocol as previously described [20]. Briefly, tissue was minced in Buffer X, transferred to preheated Buffer X (37 °C) with Enzyme Mixture 1 (Buffer X with Enzyme P), and triturated. After a 10 minutes incubation at 37 °C, the mixture was triturated again and Enzyme Mixture 2 (Buffer Y with Enzyme A) was added. Following another 10 minutes incubation at 37 °C, the mixture was triturated and 10 mL of modified Dulbecco’s Phosphate-Buffered Saline (DPBS) buffer (Gibco; 14287080) was added. The mixture was allowed to settle for 2 minutes, then filtered through a 70-µm cell strainer (Falcon; 352350) and pelleted (600 xg, 10 minutes, 4 °C). Myelin was removed by resuspending cell pellets in 0.9 M sucrose in DPBS with calcium and magnesium (Gibco; 14040) and centrifuging (850 xg, 15 minutes, 4 °C). The cell pellet was then resuspended in FACS buffer (5% FBS in 1x PBS) and passed through a 40-µm cell strainer (Falcon; 352340) before final centrifugation (600 xg, 5 minutes, 4 °C). The single cell suspension was first blocked with 1:65 Fc block (BD Pharmingen; 553142) for 10 minutes on ice before being stained with 1:10 CD90.2/Thy-1.2–APC/Fire™ (BioLegend; 105348), 1:50 ACSA2–PE-Vio^®^770 (Miltenyi Biotec; 130-116-246), 1:200 CD11b–BV421 (BioLegend; 101236), 1:10 anti-Galactocerebroside–FITC (Merck Milli-Mark^®^; FCMAB312F), 1:50 CD31–APC (Miltenyi Biotec; 130-123-813), and 1:1000 Fixable Viability Dye eFluor™506 (Invitrogen eBioscience™; 65-0866-14). Cells were then fixed with 2% paraformaldehyde for 10 minutes on ice. After fixation, cells were stained with 1:100 KN343 (Tokyo Chemical Industry Co. Ltd.; A3428) for 20 minutes on ice, followed by staining with goat anti-mouse IgM–PE (R&D Systems; F0116).

For adult primary microglia, cortex and striatum were dissected and dissociated in cold Medium A (0.5% glucose and 15 mM HEPES in Hank’s Balanced Salt Solution (HBSS, without phenol red) using a Dounce homogenizer as previously described [20]. The resulting mixture was filtered through a 70-µm cell strainer (Falcon; 352350) and pelleted (600 xg, 10 minutes, 4 °C). The cell pellet was resuspended in 1 mL MACS buffer (0.5% bovine serum albumin (BSA) and 2 mM EDTA in PBS) with 1:10 myelin removal beads (Miltenyi Biotec; 130-096-433) and incubated for 10 minutes on ice. Myelin was depleted by passing the cell suspension through an LD column (Miltenyi Biotec; 130-042-901) placed on a QuadroMACS™ Separator (Miltenyi Biotec, Bergisch Gladbach, North Rhine-Westphalia, Germany). The cell suspension was filtered through a 40-µm cell strainer (Falcon; 352340) and centrifuged again (600 xg, 5 minutes, 4 °C). The microglia-enriched pellet was resuspended in FACS buffer (5% FBS in 1x PBS) and blocked with 1:65 Fc block (BD Pharmingen; 553142) for 10 minutes on ice before being stained with 1:100 CD11b–BV421 (BioLegend; 101236), 1:100 P2RY12–Alexa Fluor^®^ 488 (BioLegend; 848016), 1:200 CX3CR1– Alexa Fluor^®^ 647 (BioLegend; 149004), 1:50 AXL–Alexa Fluor^®^ 700 and, 1:1000 Fixable Viability Dye eFluor™ 506 (Invitrogen eBioscience™; 65-0866-14). Cells were then fixed with 2% paraformaldehyde and permeabilized with 0.3% saponin for 10 minutes on ice, followed by staining with 1:100 MerTK–PE/Cyanine7 (BioLegend; 151522) and 1:100 Galectin-3–PE (Invitrogen eBioscience™; 12-5301-82) in FACS buffer containing 0.3% saponin. All flow cytometry analyses were performed using an Attune CytPix Flow Cytometer (Thermo Fisher Scientific, Waltham, MA, USA).

### Mass Cytometry Analysis

The dissociated mouse brain cells were fixed with 1.6% paraformaldehyde (Electron Microscopy Sciences, Hatfield, PA, USA) in serum-free RPMI 1640 (Gibco) at room temperature for 10 minutes and stained with immune cell markers and Siglecs. Each independent replicate was generated from pooled samples of 3–4 mice for each genotype. For surface marker staining, cells were incubated with a cell-surface antibody cocktail prepared in cell staining medium (CSM), containing 1x PBS, 0.5% protease-free bovine serum albumin (BSA), and 0.02% NaN_3_, in a final volume of 100 μL for 1 hour at room temperature. After washing once with CSM, cells were permeabilized with 100% ice-cold methanol for 10 minutes. For intracellular marker staining, cells were washed twice with CSM and stained with an intracellular antibody cocktail prepared in CSM in a final volume of 100 μL for 1 hour at room temperature. After staining, cells were washed twice with CSM, and then stained with Cell-ID Intercalator-Ir (Fluidigm Corporation; 191Ir and 193Ir) at a final concentration of 125 nM in 1000 μL of 1.5% fresh paraformaldehyde (diluted in 1x PBS) overnight at 4 °C for DNA staining. Finally, cells were resuspended in Milli-Q water containing EQ™ Four Element Calibration Beads (Fluidigm Corporation, San Francisco, CA, USA) and acquired on a CyTOF2 mass cytometer (Fluidigm Corporation, San Francisco, CA, USA).

The resulting raw flow cytometry standard files were normalized to the bead signals using the Premessa R package (https://github.com/ParkerICI/premessa). The data were uploaded to Cytobank, and marker intensities were arcsinh-transformed with a cofactor of 5 prior to analysis. Singlets, live cells, and CD90.2^−^ populations were gated. An initial Uniform Manifold Approximation and Projection (UMAP) of CD90.2^−^ population was conducted using 27 immune markers: AXL, CCR2, CD11b, CD11c, CD19, CD27, CD3, CD39, CD4, CD40, CD44, CD45, CD49d, CD49e, CD80, CXCR3, F4/80, Ly6C, Ly6G, MerTK, MHCII, P2RY12, PDL1, TIM-3, TIM-4, TMEM119, and TREM2. The UMAP parameters included all events, neighborhood size = 15, and minimum distance = 0.01. Next, the cluster exhibiting higher CD39 and P2RY12 expression was selected for a secondary UMAP analysis focusing on Siglecs (Siglec-1, Siglec-15, Siglec-2, Siglec-3, Siglec-E, Siglec-F, Siglec-G, Siglec-H). The parameters were equal event numbers, neighborhood size = 20, and minimum distance = 0.02. FlowSOM clustering was then applied to the secondary UMAP-1 and UMAP-2 data, resulting in the identification of nine metaclusters. Antibodies used for mass cytometry are listed in Table S5.

### Single-nuclei RNA Sequencing

Cortical and striatal tissues were dissociated according to the procedures by 10x Genomics in Nuclei Isolation from Embryonic Mouse Brain for Single Cell Multiome ATAC + Gene Expression Sequencing (CG000366, 10x Genomics). Nuclei were emulsified with gel beads using the 10x Genomic Chromium Next GEM Single Cell Multiome ATAC + Gene Expression kit (CG000338, 10x Genomics) according to the manufacturer’s instructions. Libraries were prepared and sequenced on NovaSeq 6000 platform (Illumina, San Diego, CA, USA).

Raw sequencing data were processed with Cell Ranger software (3.1.0, 10x Genomics). Reads were aligned to the *Mus musculus* reference genome (mm10) and Ensembl gene annotation (release 102). Genes with the following biotypes were exclude, IG gene, ncRNA, TEC, and TR gene. Downstream gene profile analyses were performed using the Seurat package in R Seurat 4.0.4 [21]. Data preprocessing included filtering cells based on the following criteria: (1) cells with unique molecular identifier (UMI) counts below 200 or above 6,000 per cell were removed; (2) cells with fewer than 200 genes per cell were removed; and (3) cells with more than 10% mitochondrial gene expression were excluded. After filtering, a total of 7,431 and 6,307 cells were retained from the R6/2 and WT mice, respectively.

The Seurat objects for each condition were processed using “SCTransform” for normalization and variance stabilization, regressing out mitochondrial gene percentages [22]. Integration of the datasets was performed using the SCTransform-based workflow. Specifically, 3,000 integration features were selected using the “SelectIntegrationFeatures” function, and integration anchors were identified with the “FindIntegrationAnchors” function. The datasets were integrated using the “IntegrateData” function, producing a single combined dataset for downstream analysis.

Principal component analysis (PCA) was performed on the integrated dataset using the top 100 principal components. Uniform Manifold Approximation and Projection (UMAP) was computed using the first 65 PCs to generate a two-dimensional representation of the data. Clustering was performed using the “FindNeighbors” and “FindClusters” functions, with a resolution of 0.7, to identify distinct cell clusters. Cell clusters were visualized using UMAP with labels applied for individual clusters. The cluster markers and differentially expressed genes of each cluster were identified with “FindAllMarkers” and “FindMarkers”, respectively, and were subjected to GO analysis to identify the major biological processes.

Cell annotation was performed by combining results from “SingleR”, “PanglaoDB”, and the “ScType” [23–25]. SingleR was used to infer cell identities based on reference datasets, while PanglaoDB provided additional insights into cell types based on known marker genes. The sctype package further validated annotations by leveraging a curated database of cell type markers. The following markers were applied to distinguish striatal (*Ppp1r1b*, *Rgs9*, *Kit*, *Sst*, *Gad1*) and cortical (*Slc17a7*, *Slc30a3*, *Rgs6*, *Car10*) neurons. This integrative approach ensured robust and accurate cell type identification across all datasets.

Data processing and analysis steps were consistent across all samples, and all computational analyses were performed in R. Cluster-specific and condition-specific visualizations were generated to explore cell type distributions and potential condition-dependent changes in cellular populations.

### Glycomic Analysis

N-glycans were released by PNGase F from digested glycoproteins extracted from the striatum of mouse brains exactly as described in previous work [26]. The released N-glycans were then reduced by 250 mM NaBH_4_ in 2M NH_3_ at 37 °C for 3 hours and neutralized by 30% acetic acid and desalted using Dowex 50W-X8 resin (Merck, Darmstadt, Germany) and C_18_ Sep-Pak cartridge (Waters Corporation, Milford, MA, USA). Reduced N-glycans were dimethylamidated by reacting with 250 mM 1-ethyl-3-(3-(dimethylamino)propyl)-carbodiimide (EDC), 500 mM 1-hydroxybenzotriazole (HOBt), and 250 mM dimethylamine in DMSO at 60 °C for 1.5 hours [27]. After drying at 60 °C in Speed-Vac, excess reagents were removed by passing through a Sep-Pak C_18_ cartridge (Waters Corporation, Milford, MA, USA) in 5% acetic acid. Derivatized N-glycans were permethylated by DMSO/NaOH/iodomethane at 4 °C for 3 hours as described before [28]. Permethylated N-glycans were fractionated by Waters® OASIS MAX cartridge (Waters Corporation, Milford, MA, USA) with the following eluent: 95% acetonitrile (for neutral glycans), 1 mM ammonium acetate in 80% acetonitrile (for singly negatively charged glycans), and 100 mM ammonium acetate in 60% acetonitrile and 20% methanol (for doubly and multiply negatively charged glycans). Prior to MS analysis, the permethylated glycans were further cleaned using ZipTip C18 in 10% acetonitrile and eluted in 75% acetonitrile/0.1% formic acid. Glycan samples were dissolved in 100% acetonitrile and mixed 1:1 with matrix (2,5- dihydroxybenzoic acid or 3,4 -diaminobenzophenone) and subjected to MALDI-MS analysis in reflectron positive and negative ion modes on an AB SCIEX MALDI TOF/TOF 5800 system (SCIEX, Toronto, Canada). Permethylated N-glycans were subjected to nanoLC-HCD MS^2^ analysis on an Orbitrap Fusion™ Tribrid™ Mass Spectrometer (Thermo Fisher Scientific, Waltham, MA, USA), using a ReproSil-Pur 120 C18-AQ column (120 Å, 1.9 µm, 75 µm x 200 mm; packed by ESI Source Solution LLC, Woburn, MA, USA). The solvent system consisted of buffer A (100% H_2_O with 0.1% formic acid) and buffer B (100% acetonitrile with 0.1% formic acid), with a 60 minutes linear gradient of 30–80% B at a constant flow rate of 300 nL/min. Other LC-MS/MS settings were as described previously [28, 29].

### Immunofluorescence Staining and Image Analyses

Immunofluorescence staining was performed as described [30]. Briefly, brain sections (20 μm) were permeabilized with 0.2% Triton X-100 and blocked with 4% BSA for 1 hour at room temperature. The brain sections were incubated with primary antibodies) for two overnights at 4 °C in a humidified chamber. The primary antibodies included NeuN (Merck Millipore; ABN78), mEM48 (Merck Millipore; MAB5374), Darpp32 (Santa Cruz; sc-271111), Iba1 (FUJIFILM Wako Pure Chemical Corporation; 019-19741), and CD68 (Bio-Rad; MCA1957). After extensive washing, the brain sections were incubated with the corresponding secondary antibody for 90 minutes at room temperature. Nuclei were stained with Hoechst. Images were captured using a Zeiss LSM 700 stage confocal microscope with Zen 2012 software (Carl Zeiss, Oberkochen, Baden-Wiirttemberg, Germany), and analyzed by MetaMorph Microscopy Automation & Image Analysis software (Molecular Devices, San Jose, CA, USA). For each replicate, a data point represents the average of three images (fields of view) acquired from a single coverslip.

### Proximity Ligation Assay (PLA)

Cells were fixed with 4% paraformaldehyde and 4% sucrose in PBS for 15 minutes, followed by permeabilization with 0.05% NP-40 in PBS for 5 minutes. After blocking with 4% BSA for 1 hour at room temperature, samples were incubated with anti-CD22 (R&D Systems; AF2296) and anti-V5 (Invitrogen; SV5-Pk1) overnight at 4 °C in a humidified chamber. PLA was performed according to the manufacturer’s protocol and as described previously [30]. Coverslips were mounted with EverBrite™ Hardset Mounting Medium with DAPI (BIOTREND Chemikalien GmbH, Koln, Germany) before imaging with Zeiss LSM 700 stage confocal microscope. For each replicate, a data point represents the average of five images (fields of view) acquired from a single coverslip.

### Extracellular Vesicle Isolations

C8-D1A^Vector^ or C8-D1A^CHST2^ cells were cultured in exosome-depleted fetal bovine serum (Invitrogen, Carlsbad, CA, USA) for extracellular vesicle (EV) collection. Conditioned medium harvested from C8-D1A^Vector^ or C8-D1A^CHST2^ cells was centrifuged at 2,000 xg for 30 minutes and concentrated with Amicon Ultra-15 Centrifugal Filter Units (3 kDa cutoff; Merck, Darmstadt, Germany) at 4,000 xg for 55 minutes. The concentrated conditioned medium was then mixed with 0.5 volumes of Total Exosome Isolation reagent (Invitrogen, Carlsbad, CA, USA). After overnight incubation at 4°C, samples were centrifuged at 10,000 xg for 60 minutes at 4 °C. EV pellets were resuspended in PBS for subsequent analysis. For Western blot analysis, KN343 levels were compared using equal 30 μL volumes of EV lysates and normalized to Alix. Proteinase K protection assays were conducted as detailed elsewhere [31]. Briefly, EVs were treated with 1 mg/mL proteinase K for 30 minutes at 37 °C, followed by enzyme inactivation with 2 mM phenylmethanesulfonylfluoride (PMSF).

### Co-Immunoprecipitation and Western Blot Analysis

Western blotting was performed as described previously [30]. Transferred blots were incubated overnight at 4 °C with the following primary antibodies: anti-CD22 (R&D Systems; AF2296), anti-V5 (Invitrogen; SV5-Pk1), and anti-GAPDH (GeneTex; GTX100118). Immunoreactive bands were detected by Clarity™ enhanced chemiluminescence (ECL) substrate (Bio-Rad, Hercules, CA, USA) and recorded using Azure 300 Imaging System (Dublin, CA, USA).

### Magnetic Resonance Imaging (MRI)

T2-weighted imaging (T2WI) of the mice brain was performed using a Bruker BioSpec 70/20 MRI Scanner (Bruker Corporation, Billerica, MA, USA) at the Animal Imaging Facility, Taiwan Mouse Clinic, Academia Sinica. T2WI images were acquired with a repetition time (TR) of 6500 ms, an echo time (TE) of 42 ms, a field of view (FOV) of 18 x18 mm, 64 coronal slices at a slice thickness of 0.25 mm, and an acquisition matrix of 256 x 256. Data analysis was conducted using Amira and Avizo software (Thermo Fisher Scientific, Waltham, MA, USA). Regions of interest (ROIs), including the whole brain, cortex, striatum, corpus callosum, hippocampus, and cerebellum, were delineated based on the Allen Mouse Brain Atlas. The volume of each region was calculated by summing the voxel volumes across the relevant slices.

### Statistical analysis

Data are presented as the mean ± SEM. Statistical analyses were performed using GraphPad Prism 9 (GraphPad Software, San Diego, CA, USA) software. Statistical significance was determined by paired or unpaired *t*-test, one-way ANOVA, or two-way ANOVA as indicated, with *p* values < 0.05 considered statistically significant.

## Results

### CD22 transcript is upregulated in the striatum of HD patients and HD mice

Given the high abundance of sialic acids in mammalian brains, we hypothesized that Siglecs, each with distinct ligand specificities, may play crucial roles in regulating the activation and deactivation of microglial functions. To analyze the disease-associated changes in the expression of Siglecs in the striatum, the brain region most sensitive to mHTT, we performed RT-qPCR using cDNAs prepared from post-mortem caudate nuclei of non-HD and HD subjects. As shown in Figure 1a, transcripts for CD22 (Siglec-2), CD33 (Siglec-3), MAG (Siglec-4), Siglec-5, Siglec-6, Siglec-8, Siglec-9, Siglec-11, Siglec-12, Siglec-14, Siglec-15, and Siglec-16 were detected in the post-mortem caudate nucleus. In contrast, transcripts of Siglec-1, Siglec-7, and Siglec-10 were not detected. Notably, CD22 and Siglec-16 levels were significantly upregulated in the caudate nuclei of HD subjects. We chose to focus on CD22 because it is conserved between mice and humans, whereas Siglec-16 lacks a direct ortholog in mice and is non-functional in a substantial proportion of the human population [32].

**Figure 1.**
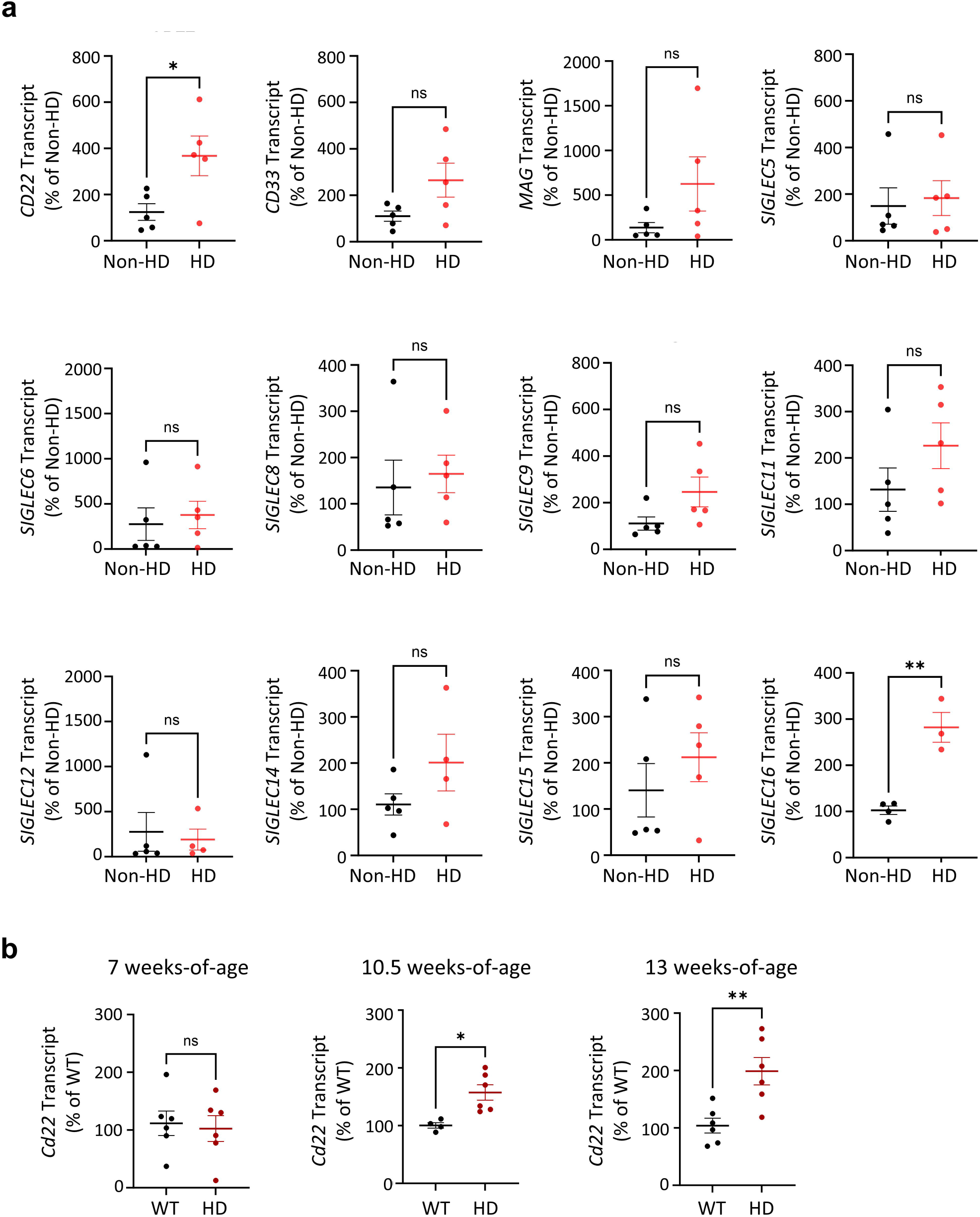
CD22 is upregulated in the striatum of HD patients and HD mice. (**a**) qPCR analysis using cDNA prepared from post-mortem caudate nucleus of non-HD and HD subjects detected transcripts for CD22 (Siglec-2), CD33 (Siglec-3), MAG (Siglec-4), Siglec-5, Siglec-6, Siglec-8, Siglec-9, Siglec-11, Siglec-12, Siglec-14, Siglec-15, and Siglec-16. Note that, due to polymorphic frameshift insertion mutation or null polymorphism in some individuals, Siglec-12, -14, and -16 are not expressed in all subjects [1–3]. Importantly, CD22 and Siglec-16 were significantly upregulated in HD samples. Data are presented as mean ± SEM (n = 5 per group). (**b**) qPCR analysis using cDNA prepared from striatum of HD mice at 7 weeks (pre-symptomatic stage), 10.5 weeks (symptomatic stage), and 13 weeks (late-stage). Mouse CD22 is significantly upregulated in the striatum of HD mice at disease onset and further increases at late-stage. Data are presented as mean ± SEM (n = 4-6 per group). \**p* < 0.05 and ** *p* < 0.01 by unpaired *t*-test.

In mouse brains, CD22 is expressed exclusively in microglia, whereas in human brains, it is predominantly expressed in oligodendrocytes (Figure S1) [33, 34]. Intriguingly, analysis of a publicly available single-nucleus RNA sequencing (snRNA-seq) dataset from the post-mortem caudate and putamen of HD patients revealed that CD22 is significantly upregulated in microglia but not in oligodendrocytes, suggesting a potential role in microglial function in HD (Table S2) [35]. We further analyzed the mRNA transcript levels of moulse *Cd22* in the striatum of HD mice (R6/2 [17], a transgenic mouse model of HD). RT-qPCR results showed that *Cd22* transcripts were significantly upregulated in the striatum of HD mice at 10 weeks of age (symptomatic stage) and further increased at 13 weeks (late stage). No increase in *Cd22* expression was observed at 7 weeks of age (pre-symptomatic stage) (Figure 1b).

### Mass cytometry analysis reveals heterogeneous Siglec expression and expansion of CD22-positive microglia in HD mice

Despite comprising only about 10% of total brain cells, microglia exhibit remarkable heterogeneity and dynamic functionality. Distinct microglial subsets can undergo further differentiation into stimulus-dependent states during development, aging, and disease [36–38]. To analyze which Siglecs are expressed on mouse microglia and whether their expression levels change in the brain of HD model mice, we developed a Siglec antibody panel and characterized the expression patterns of mouse Siglecs on microglia using mass cytometry (Figure 2a) [39]. Mass cytometry analysis revealed a differential distribution pattern of Siglecs in microglia, with Siglec-H (a molecular signature for microglia) being widely expressed, followed by CD33 and Siglec-E. In contrast, CD22, Siglec-F, Siglec-G, and Siglec-15 showed more restricted expression patterns. These findings support the notion that microglia are highly heterogeneous, with individual subsets expressing distinct receptors and molecules that likely reflect specialized functional roles. Importantly, CD22 is enriched in a cluster of microglia (Metacluster 8, Figure 2e) that also expresses a high level of disease-associated microglia (DAM) genes (i.e., AXL), and low levels of homeostasis gene (i.e., P2RY12) (Figure 2f). In addition, we found that the CD22-positive subset was less prominent in wild-type microglia but significantly expanded in HD microglia (Metacluster 8, Figure 2b-d). This observation was further validated by flow cytometry (Figure S2), confirming that CD22 is upregulated at both the transcript and protein levels.

**Figure 2.**
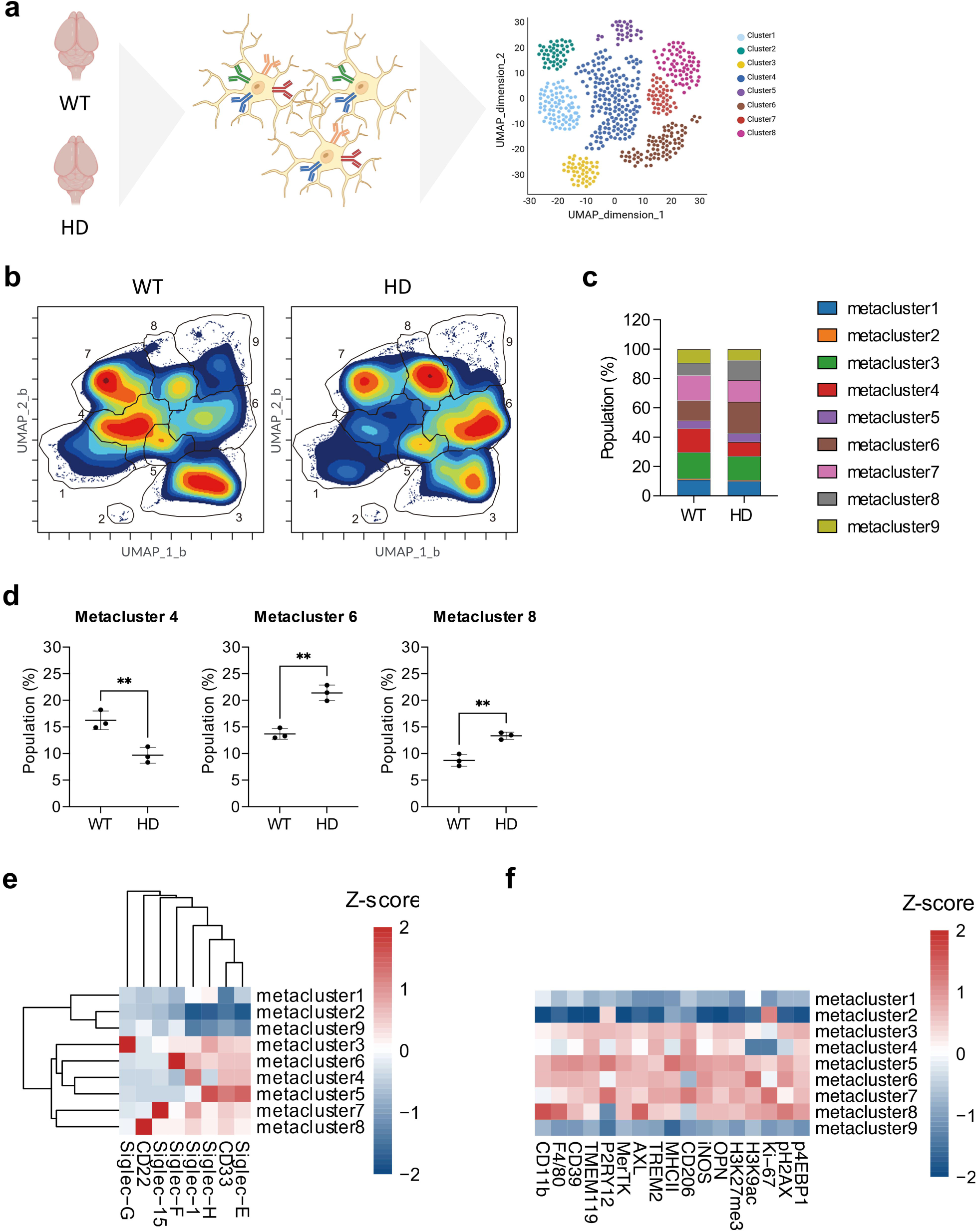
Mass cytometry revealed heterogeneous Siglec expression and expansion of CD22⁺ microglia in HD mice. (**a**) Schematic overview of mass cytometry analysis of Siglec expression in adult primary microglia isolated from WT and HD brains at 13 weeks of age. (**b**) UMAP dimension reduction and FlowSOM clustering of mass cytometry data from WT and HD mice. Mass cytometry analysis demonstrated a differential distribution of Siglecs in microglia. (**c**) Population percentages and statistical analysis were conducted for each metacluster identified by FlowSOM clustering. (**d**) Significant changes were observed in the HD population compared to WT in metaclusters 4, 6, and 8. (**e**) Detailed expression profiles of Siglecs and immune features across metaclusters. Data are presented as mean ± SD (n = 3 independent replicates). \*\**p* < 0.01 by unpaired *t*-test.

### CD22 suppresses the phagocytic function of microglia independently of its sialic acid-binding domain

CD22 was recently identified as a negative regulator of microglial phagocytosis during aging [16]. However, It remains unclear whether CD22 also affects other aspects of microglial function. To test if the upregulation of CD22 affects microglial functions, we established microglial cell lines (BV2) stably expressing CD22 (designated BV2-CD22) or a control vector (BV2-miR9T) (Figure S7a) and subjected them to functional tests.

We found that over-expression of CD22 impairs the phagocytic uptake of pHrodo Red-conjugated Zymosan beads (Figure 3a-b). Similar results were observed in primary microglia isolated from wild-type (CD22⁺/⁺) and CD22⁻/⁻ mice (Figure 3c-d). On the other hand, CD22 over-expression did not affect the expression of pro-inflammatory cytokines (e.g., IL-1β (*Il1b*) and IL-6 (*Il6*)) or anti-inflammatory cytokines (e.g., IL-10 (*Il10*)) upon LPS stimulation (Figure S7b-c). Likewise, CD22 over-expression did not alter the expression of oxidative stress markers (e.g., HIF1α (*Hif1a*) and NRF2 (*Nfe2l2*)) or oxidative stress-induced inflammation markers (e.g., COX-2 (*Ptgs2*)) following elevated oxidative stress (Figure S7d-e). Taken together, these results suggest that aberrant upregulation of CD22 primarily affects the phagocytic capacity of microglia.

**Figure 3.**
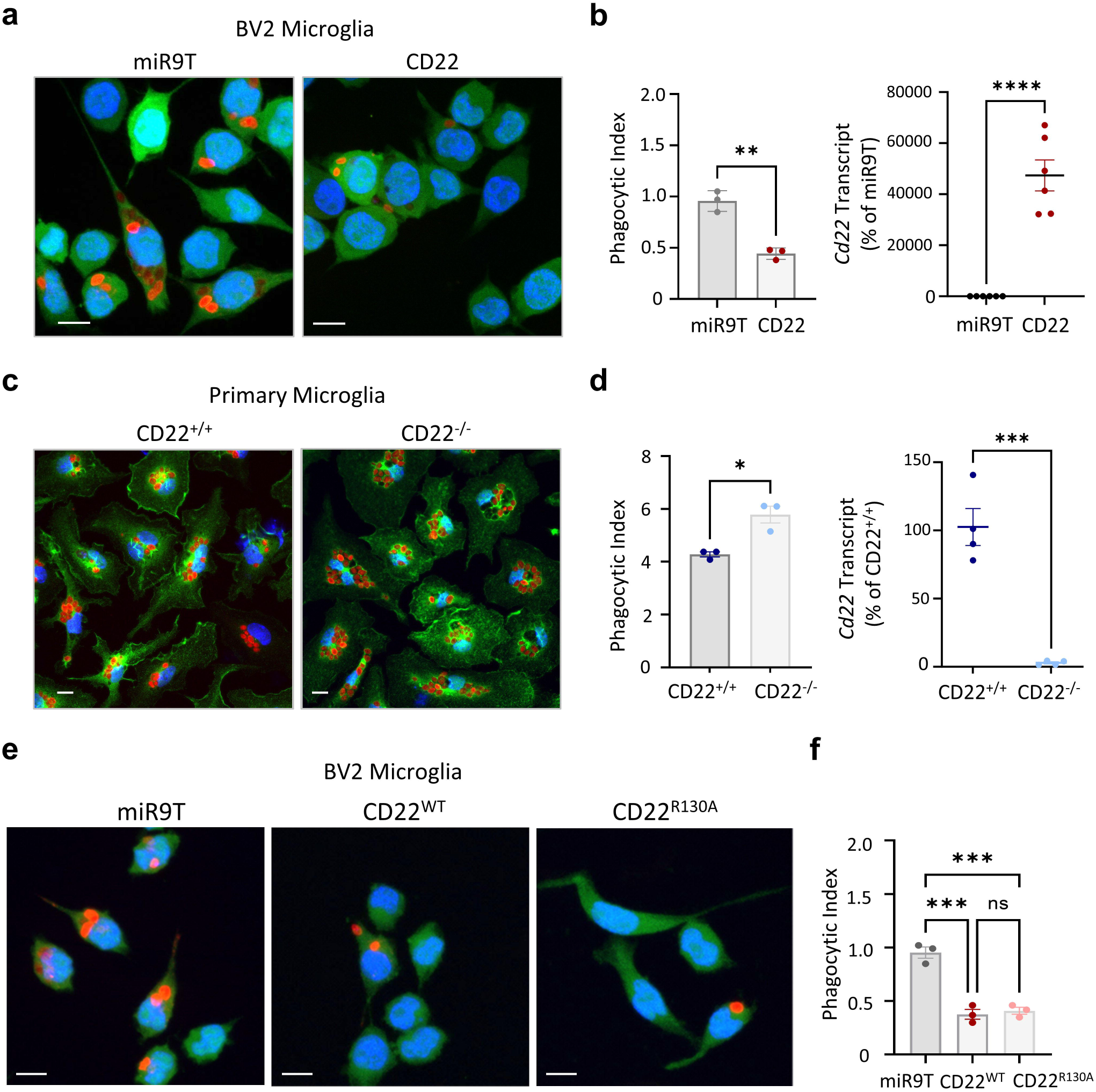
CD22 suppresses phagocytic function of microglia independently of its sialic acid binding domain. (**a-b**) Phagocytic function of BV2 microglia with stable expression of vector control (miR9T) and CD22 was analyzed using pHRodo red Zymosan bioparticles. Overexpression of CD22 reduced the phagocytic function of BV2 microglia. (**c-d**) Phagocytic function was similarly assessed in neonatal primary microglia derived from wild-type (CD22^WT^) and CD22-knockout (CD22^KO^) mice. Knockout of CD22 enhanced the phagocytic function of primary microglia. (**e**) Phagocytic function of BV2 microglia with stable expression of vector control (miR9T), wild-type CD22 (CD22^WT^), or sialic acid binding mutant CD22 (CD22^R130A^) was analyzed using pHRodo red Zymosan bioparticles. Expression of either CD22^W^ or CD22^R130A^ reduced the phagocytic function of BV2 microglia. Phagocytic index was quantified by counting the number of internalized particles per cell in the field. Data are presented as mean ± SEM (n = 3 independent replicates). \**p* < 0.05, \*\**p* < 0.01, \*\*\**p* < 0.001 and \*\*\*\**p* < 0.0001 by unpaired *t*-test or one-way ANOVA. Scale bar = 10 µm.

CD22 does not function as a direct phagocytic receptor but likely serves as a co-receptor that modulates the activity of phagocytic receptors on the same cell. To determine whether CD22-mediated suppression of phagocytosis depends on its sialic acid-binding domain, we generated a BV2 microglial cell line expressing a sialic acid-binding-deficient mutant (CD22^R130A^). Our results show that both BV2-CD22^WT^ and BV2-CD22^R130A^ cells exhibited reduced phagocytic activity compared to control BV2-miR9T cells (Figure 3e-f). This finding indicates that CD22-mediated suppression of phagocytosis is independent of its sialic acid-binding domain.

### CD22 proximity suppresses phagocytosis via canonical ITIM–ITAM signaling crosstalk

Given that CD22-mediated suppression of *cis*-phagocytic receptors is independent of its sialic acid-binding domain, we hypothesized that this suppression may occur through the canonical ITIM–ITAM signaling crosstalk. It is well documented that the association of CD22 with *cis*-receptors, such as the B cell receptor (BCR), typically induces an inhibitory signaling cascade via the recruitment of SHP-1, which dephosphorylates ITAM-bearing receptors [3, 40, 41]. Since CD22 contains an ITIM motif, we analyzed whether it inhibits Dectin-1, a phagocytic receptor that contains an ITAM motif [3, 40, 42], by assessing the uptake of pHrodo Red-conjugated Zymosan [40, 43]. To this end, Dectin-1 (*Clec7a*–V5) and CD22^WT^, CD22^R130A^ (the sialic acid binding-deficient mutant), or CD22^Y783/843/863F^ (the ITIM motif mutant) were co-expressed in HEK293T cells. Immunofluorescence staining confirmed the co-localization of CD22 and Dectin-1 (*Clec7a*–V5; Figure S8), and a proximity ligation assay (PLA) further verified their close molecular proximity (Figure 4a).

**Figure 4.**
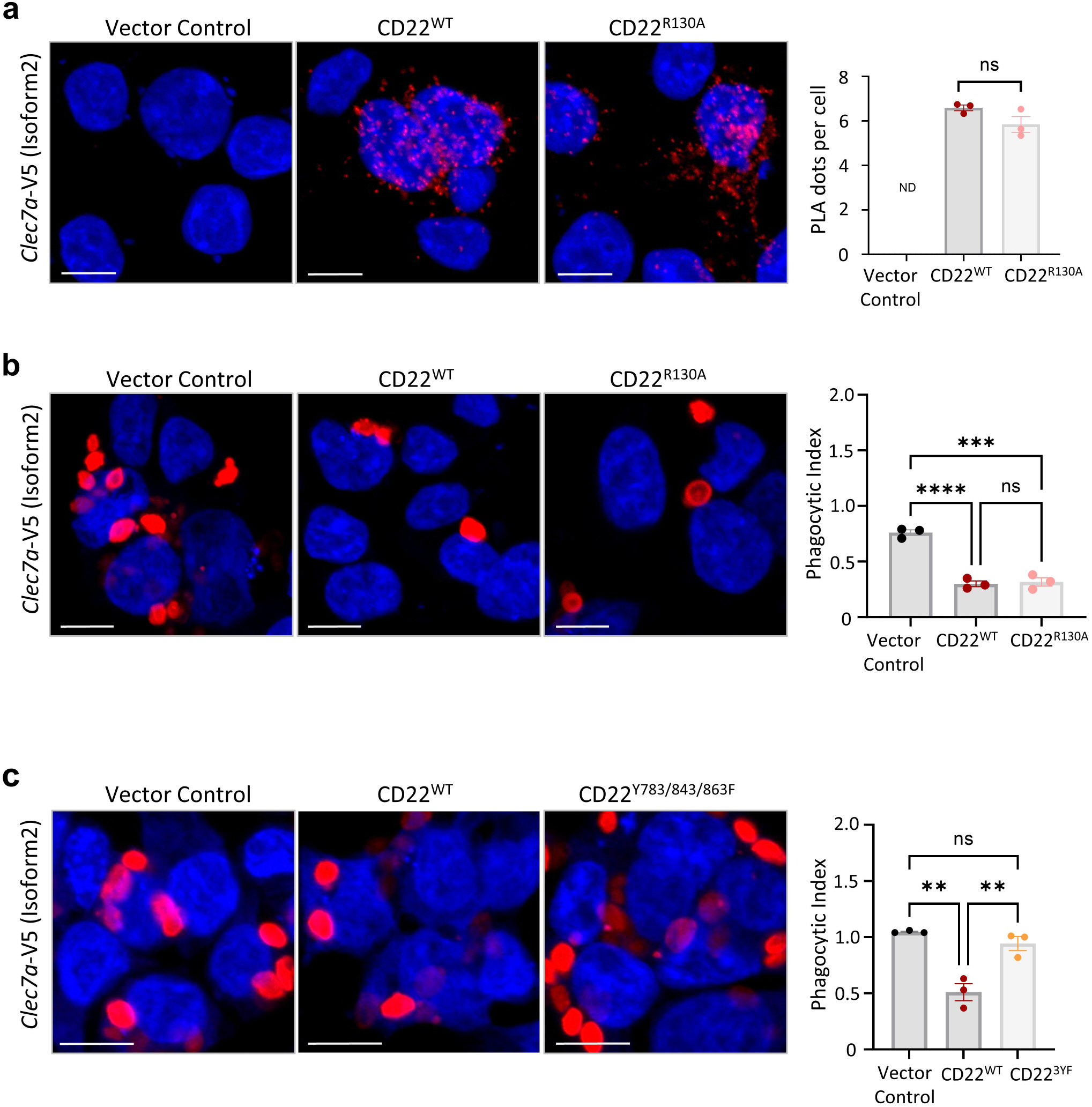
CD22 proximity suppresses phagocytosis via canonical ITIM-ITAM signaling crosstalk. The ITAM-bearing Dectin-1 (*Clec7a*) was cloned from C57BL/6 mice and tagged with V5. *Clec7a*-V5 was overexpressed in HEK293T cells along with either vector control, CD22^WT^, or CD22^R130A^. (**a**) Proximity ligation assay (PLA) using anti-CD22 and anti-V5 antibodies revealed molecular proximity between *Clec7a*-V5 and CD22^WT^ or CD22^R130A^. Data are presented as mean ± SEM (n = 3 independent replicates). Overexpression of *Clec7a*-V5 in HEK293T cells enabled the uptake of pHRodo red Zymosan bioparticles, and co-expression of CD22^WT^ or CD22^R130A^ (**b**) significantly suppressed this uptake, while co-expression of CD22^Y783/843/863F^ (**c**) did not. Data are presented as mean ± SEM (n = 3 independent replicates). \*\**p* < 0.01, \*\*\**p* < 0.001 and \*\*\*\**p* < 0.0001 by one-way ANOVA. Scale bar = 10 µm.

As expected, over-expression of Dectin-1 (*Clec7a*–V5) enabled HEK293T cells to uptake Zymosan, while co-expression of CD22^WT^ or CD22^R130A^ significantly inhibited Zymosan uptake (Figure 4b). In contrast, co-expression of CD22^Y783/843/863F^ did not inhibit Zymosan uptake (Figure 4c). These findings support the hypothesis that CD22 inhibits ITAM-bearing receptor (Dectin-1)-mediated phagocytosis through canonical ITIM–ITAM signaling crosstalk.

### An anti-CD22 neutralizing antibody restores phagocytosis in CD22-positive microglia

Previous studies in B cell lymphoma have shown that drugs conjugated to anti-CD22 antibodies can enhance the uptake of therapeutic agents [44–46], suggesting that an anti-CD22 antibody could trigger CD22 internalization. Here, we employed an anti-CD22 neutralizing antibody (Cy34.1), which targets the ligand-binding pocket of CD22, to investigate its impact on CD22 distribution and phagocytic function in a microglial cell line over-expressing CD22 (BV2-CD22). As shown in Figure 5a, treatment with Cy34.1 resulted in reduced CD22 expression on the cell surface due to enhanced internalization, as evidenced by the colocalization of CD22 (red) and Cy34.1 (grey) puncta within cells. The CD22 expression construct contains both CD22 cDNA and a GFP reporter linked via a T2A linker, which allows ribosomal skipping and independent translation of CD22 and GFP. GFP expression thus serves as a proxy for total CD22 expression in BV2-CD22 cells treated with either an IgG isotype control or Cy34.1 (Figure 5b). Treatment with Cy34.1 significantly decreased the surface level of CD22, as assessed by both immunocytochemistry and flow cytometry (Figure 5a-c). Importantly, the depletion of CD22 from the cell surface by Cy34.1 restored phagocytosis (Figure 5d). To evaluate whether this effect is functionally meaningful, we assessed the uptake of myelin debris derived from both wild-type and HD brains. The myelin debris were labeled with pHrodo Red succinimidyl ester, which fluoresces upon endocytosis and lysosomal acidification. Flow cytometry revealed that Cy34.1 treatment significantly increased the uptake of both WT and HD myelin debris by BV2-CD22 cells (Figure 5e). Consistent results were observed in primary microglia isolated from wild-type (CD22⁺/⁺) and CD22⁻/⁻ mice (Figure 5f). Together, these findings demonstrate that depleting CD22 from the cell surface by inducing its internalization relieves its inhibitory effect on *cis*-phagocytic receptors and restores phagocytosis in microglia.

**Figure 5.**
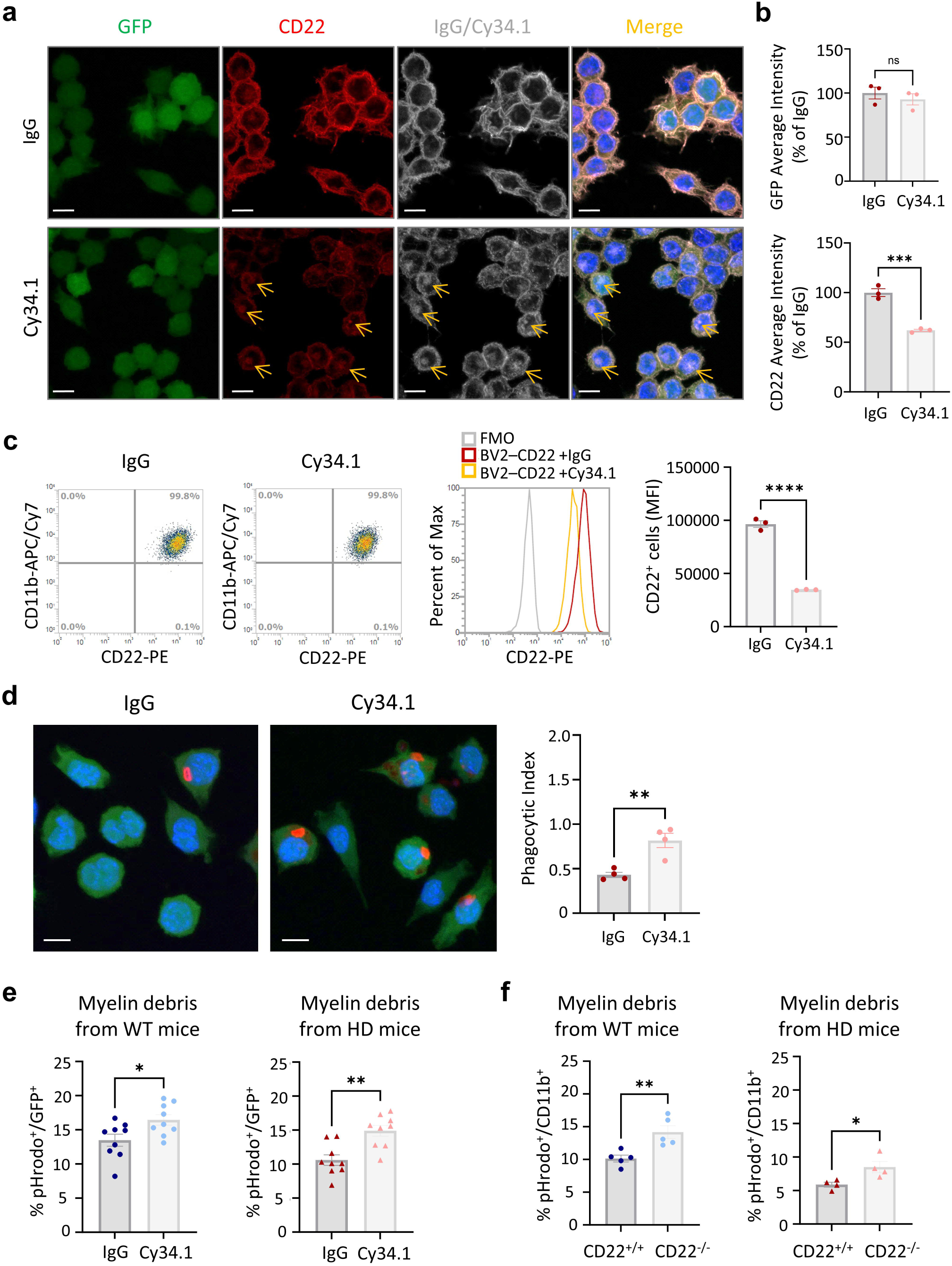
An anti-CD22 neutralizing antibody restores phagocytic function of CD22+ microglia. BV2-CD22 microglia were treated with either IgG isotype control or anti-CD22 antibody (Cy34.1) for 24 hours. (**a**) Immunofluorescence staining showed colocalization of CD22 (red) and Cy34.1 (grey) puncta (yellow arrows) within the cells. (**b**) Quantification of GFP showed that cells treated with IgG or Cy34.1 had comparable levels of CD22-GFP, while treatment with Cy34.1 reduced the overall amount of CD22. (**c**) Flow cytometry analysis confirmed similar CD22⁺ percentages (∼99.8%) across groups, but treatment with Cy34.1 reduced the amount of CD22 on the cell surface, as reflected by MFI. Data are presented as mean ± SEM (n = 3). (**d**) Phagocytosis of pHRodo red Zymosan particles was restored in BV2-CD22 cells treated with Cy34.1. Data are presented as mean ± SEM (n = 4 independent experiments). (**e-f**) Phagocytosis of pHRodo-labeled myelin debris was assessed in BV2-CD22 cells treated with Cy34.1 and neonatal primary microglia derived from wild-type (CD22^WT^) or CD22-knockout (CD22^KO^) mice. Phagocytic index was quantified as the percentage of pHRodo⁺ among CD11b⁺ microglia. Data are presented as mean ± SEM (n = 4-5 biological replicates). \**p* < 0.05, \*\**p* < 0.01, \*\*\**p* < 0.001 and \*\*\*\**p* < 0.0001 by unpaired *t*-test. Scale bar = 10 µm.

### Genetic removal of CD22 ameliorates HD pathology

To investigate the biological impact of CD22 in HD, we crossed CD22^-/-^ mice with HD mice (R6/2, Figure 6a), and assessed the effects of CD22 deletion on disease phenotypes. The R6/2 mouse model is widely used to study HD pathogenesis, as it recapitulates many features of HD, including body weight loss, motor impairment, and the accumulation of insoluble mHTT [47]. CD22^-/-^ mice appeared normal in size and exhibited no significant physical or behavioral abnormalities, suggesting that genetic deletion of CD22 does not notably affect their phenotype [48]. The absence of CD22 significantly improved rotarod performance in HD mice (Figure 6b). However, no differences were observed in body weight loss, limb clasping responses, or lifespan between HDWT (CD22^+/+^) and HDKO (CD22^-/-^) mice (Figure S9). We further analyzed insoluble mHTT aggregate accumulation in the striatum of HD mice at 13 weeks of age (late-stage). Immunofluorescence staining revealed widespread large mHTT aggregates in the striatum of HDWT mice. Notably, the area and intensity of mHTT aggregates were significantly reduced in the striatum of HDKO mice (Figure 6c, d). Additionally, the reduction of Darpp32, a marker for medium spiny neurons in the striatum, was mitigated by the removal of CD22 (Figure 6e, f). These findings suggest that CD22 upregulation during HD progression may play a detrimental role.

**Figure 6.**
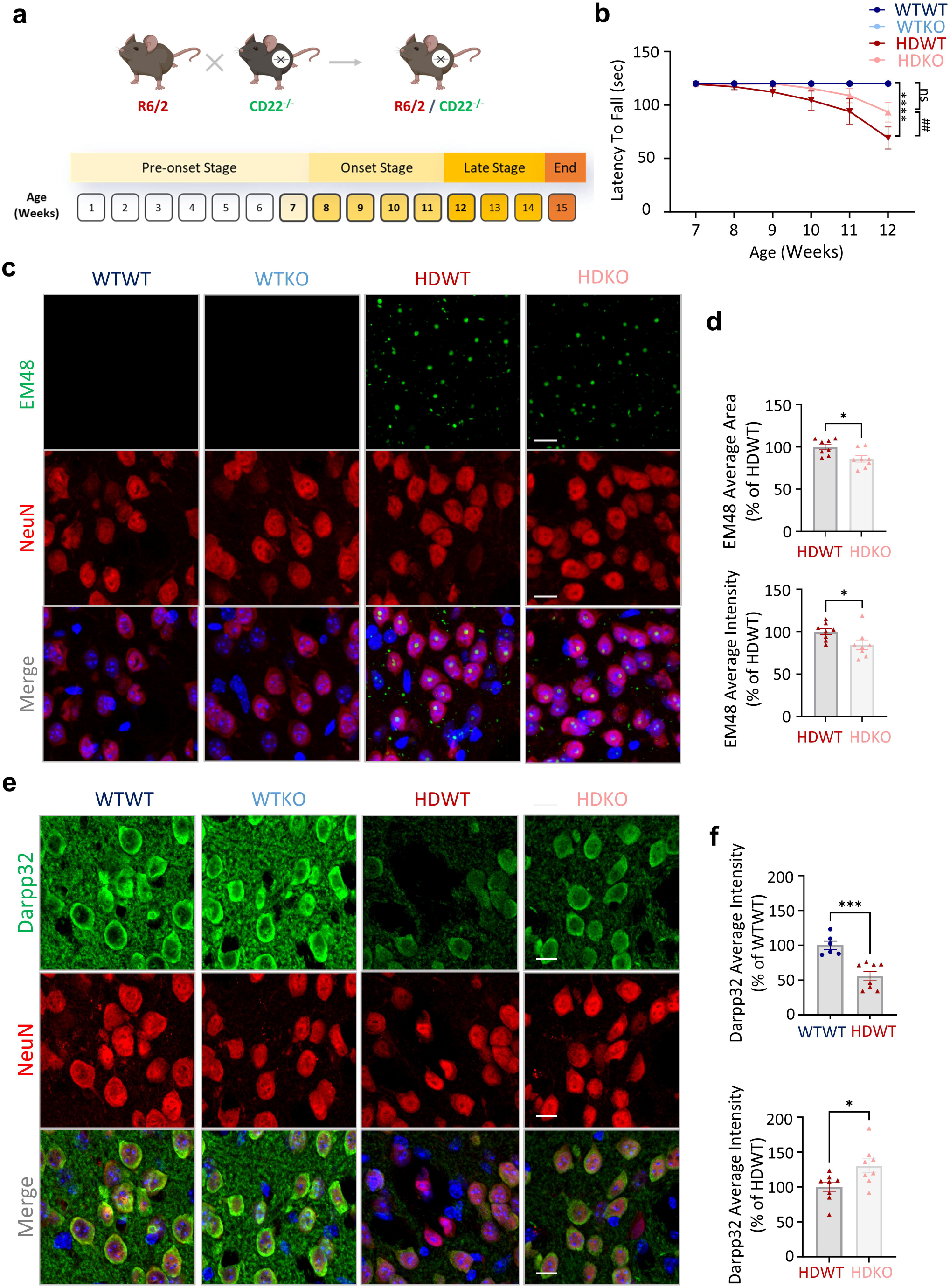
CD22 knockout ameliorates motor dysfunction and pathological outcomes in HD mice. **(a**) HD mouse model (R6/2) was crossed with CD22-knockout (KO) mice to evaluate the biological impact of CD22 *in vivo.* (**b**) Rotarod performance was assessed from 7 to 12 weeks of age. Knockout of CD22 significantly improved the rotarod performance of the HD mice. Data are presented as mean ± SEM (n = 9-12 animals per group). *Specific comparison between WT and HD mice in each condition; #Specific comparison between CD22-WT and CD22-KO in each condition by two-way ANOVA. Immunofluorescence staining of brain sections was performed to detect mHTT aggregates (green, EM48) (**c**) and the expression of the medium spiny neuron marker (green, Darpp32) (**e**) in the neurons (red, NeuN) of HD mice. (**d**) Quantification of (c). (**f**) Quantification of (e). Knockout of CD22 reduces the area and intensity of mHTT aggregates and the loss of Darpp32 in HD mice. Data are presented as mean ± SEM (n = 6-8 animals per group). \**p* < 0.05, ** ^or^ ^##^*p* < 0.01, \*\*\**p* < 0.001 and \*\*\*\**p* < 0.0001 by unpaired *t*-test. Scale bar = 10 µm.

Since microglia from patients and mice with late-stage HD exhibit impaired phagocytosis [49], we hypothesized that upregulation of CD22 in HD microglia might contribute to this deficiency. The phagocytic capacity of microglia in the striatum was assessed using CD68, a lysosomal marker specific to microglia/macrophages. Ample evidence suggests that CD68 closely correlates with phagocytic activity and thus serves as a reliable indicator of microglial phagocytosis. Immunofluorescence staining revealed strong CD68 intensity in microglia of WT brains, reflecting robust phagocytic capacity [49–51]. In contrast, both protein and transcript levels of CD68 were significantly reduced in microglia from HD brains, indicating diminished phagocytic activity (Figure 7a-b, S10). Importantly, CD68 fluorescence intensity was significantly higher in HDKO compared to HDWT brains, suggesting that CD22 deletion restores phagocytic function in HD microglia (Figure 9c).

**Figure 7.**
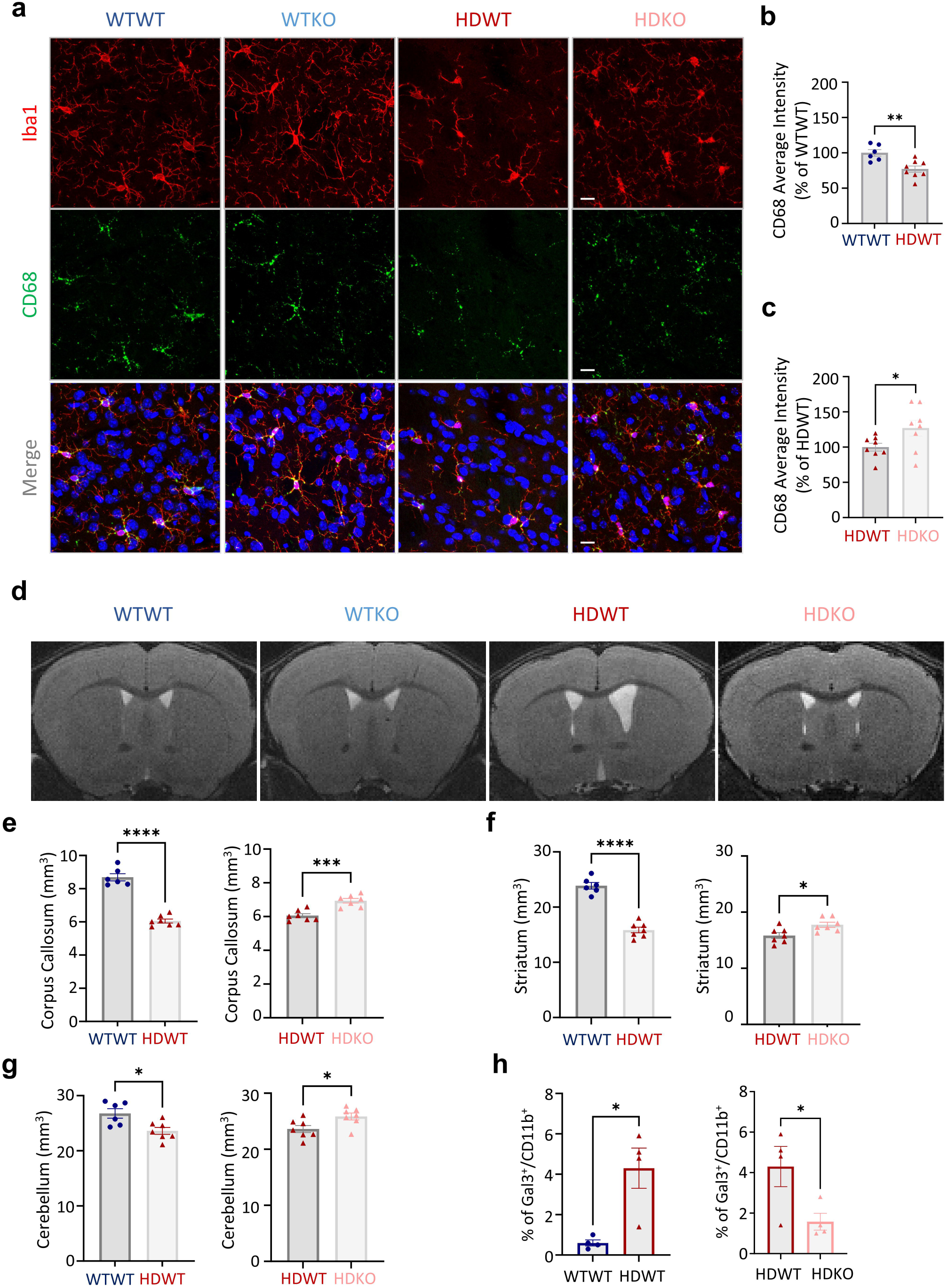
CD22 knockout restores phagocytic clearance and mitigates myelin degeneration in HD mice. (**a**) Immunofluorescence staining of brain sections was performed to detect phagocytic marker CD68 (green) in the microglia (red, Iba1). CD68 is a lysosomal marker associated with phagocytic activity and can therefore reflect the phagocytic function of the microglia *in vivo*. (**b**) Quantification showed reduced average intensity of CD68 in WT/CD22^WT^ compared to HD/CD22^WT^ mice. (**c**) Quantification revealed increased average intensity of CD68 in HD/CD22^KO^ compared to HD/CD22^WT^ mice, indicating restored phagocytic activity with CD22 deletion. Data are presented as mean ± SEM (n = 6-8 animals per group). \**p* < 0.05 by unpaired t-test. Scale bar = 10 µm. (**d**) Representative T2-weighted images of the indicated mice at 12 weeks of age. Quantification of the corpus callosum (**e**), striatal (**f**), and cerebellar (**g**) volumes showed that HD mice had reduced corpus callosum, striatal, and cerebellar volumes compared to WT mice and were partially restored by CD22 knockout. The data are presented as the means ± SEM (n = 5-8 animals per group). \**p* < 0.05, \*\*\**p* < 0.001 and \*\*\*\**p* < 0.0001 by unpaired *t*-test. (**h**) Flow cytometry analysis revealed a significant upregulation of Gal3⁺ (neuroinflammation-associated) microglia in HD mice. CD22 knockout prevented the expansion of Gal3⁺ microglia. Data are shown as mean ± SEM (n = 4 independent experiments). \**p* < 0.05, \*\**p* < 0.01, \*\*\**p* < 0.001 and \*\*\*\**p* < 0.0001 by unpaired *t*-test.

**Figure 8.**
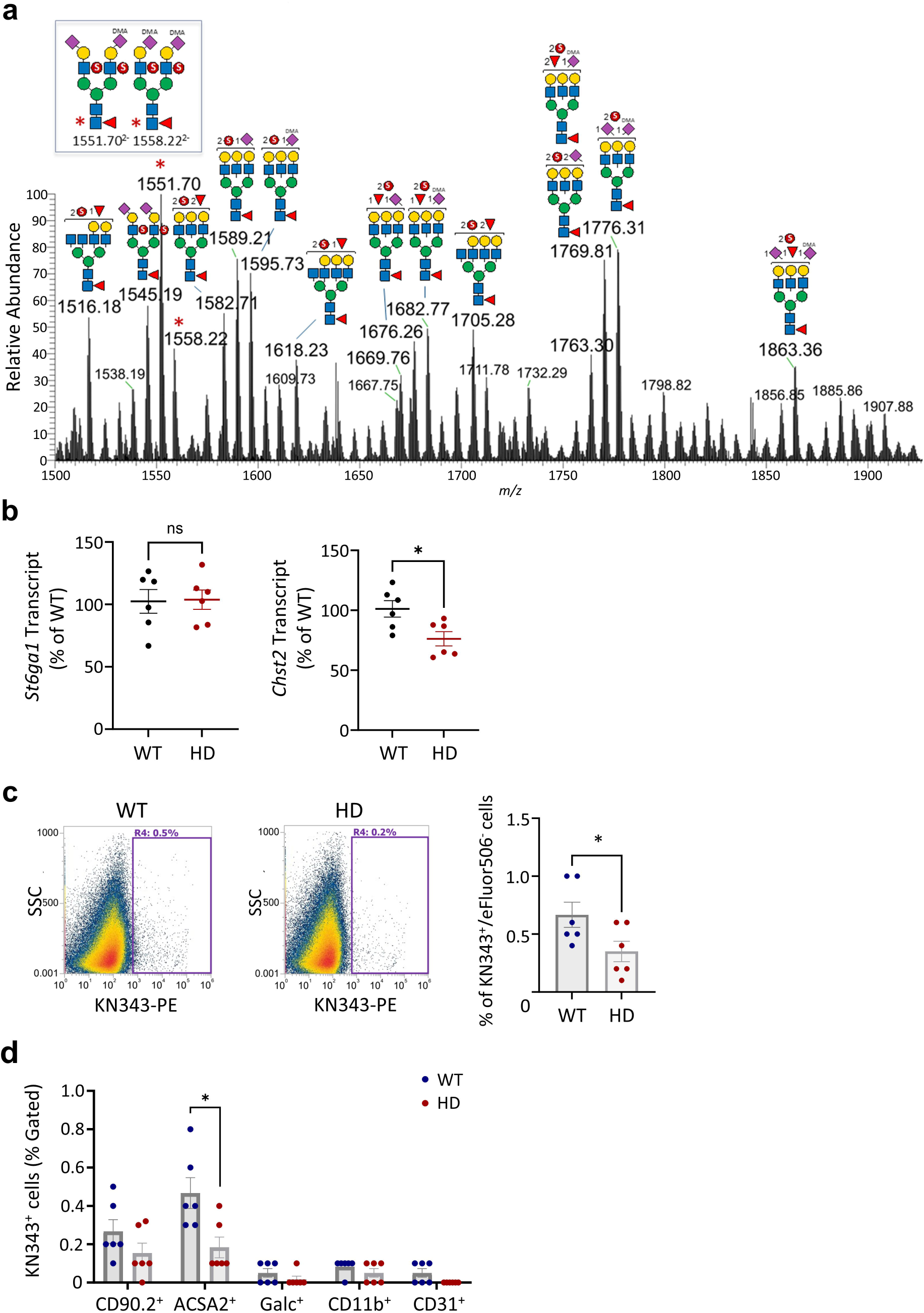
Natural ligands of CD22 in the brain are primarily expressed on astrocytes. (**a**) LC-MS/MS analysis of the permethylated sulfated N-glycans derived from striatum. α2-6-linked sialic acids were selectively dimethyl amidated (DMA) prior to permethylation. After derivatization, permethylated glycans carrying a 2-3 or 2-6-linked sialic acid are distinguished by a mass difference of 13 u. The entire sample preparation workflow yielded permethylated non-, mono- and di- multiply sulfated fractions and each was subjected to MALDI-MS screening before LC-MS/MS analysis. Only a partial mass range of the LC-MS profile of the di-/multiply sulfated N-glycan fraction averaged over 15 min is shown. All relevant disulfated glycan peaks were detected as doubly charged in negative ion mode. Most signals were further subjected to HCD MS^2^ under data dependent acquisition mode but not all yielded MS^2^ data allowing unambiguous structural assignment. The presence of α2-6-sialylated, 6-sulfo LacNAc glycotope, as inferred from the *mlz* values of the parent ions, was confirmed by the diagnostic MS^2^ ion at *mlz* 902 (Figure S14) for the two differentially disialylated biantennary N-glycans. (**b**) Transcript levels of *St6gal1* and *Chst2*, enzymes involved in the synthesis of α2-6-sialylated 6-sulfo LacNAc, were assessed in the striatum of HD mice at 13 weeks of age using qPCR analysis. Transcript levels of *Chst2* were significantly down-regulated in the striatum of HD mice at 13 weeks of age. Data are shown as mean ± SEM (n = 6 animals per group). (**c**) Flow cytometry analysis using the KN343 antibody, which specifically recognizes α2-6-sialylated 6-sulfo LacNAc, was performed to quantify the levels of α2-6-sialylated 6-sulfo LacNAc in various brain cell types from both wild-type and HD mice. Percentage of KN343^+^ cells were significantly reduced in HD brains compared to WT brains. (**d**) The analysis revealed that α2-6-sialylated 6-sulfo LacNAc (KN343^+^) was predominantly present on astrocytes, with a lesser presence on neurons. α2-6-sialylated 6-sulfo LacNAc was barely detectable on microglia, oligodendrocytes, and endothelial cells. Data are presented as mean ± SEM (n = 6 independent experiments). \**p* < 0.05 by unpaired *t*-test.

**Figure 9.**
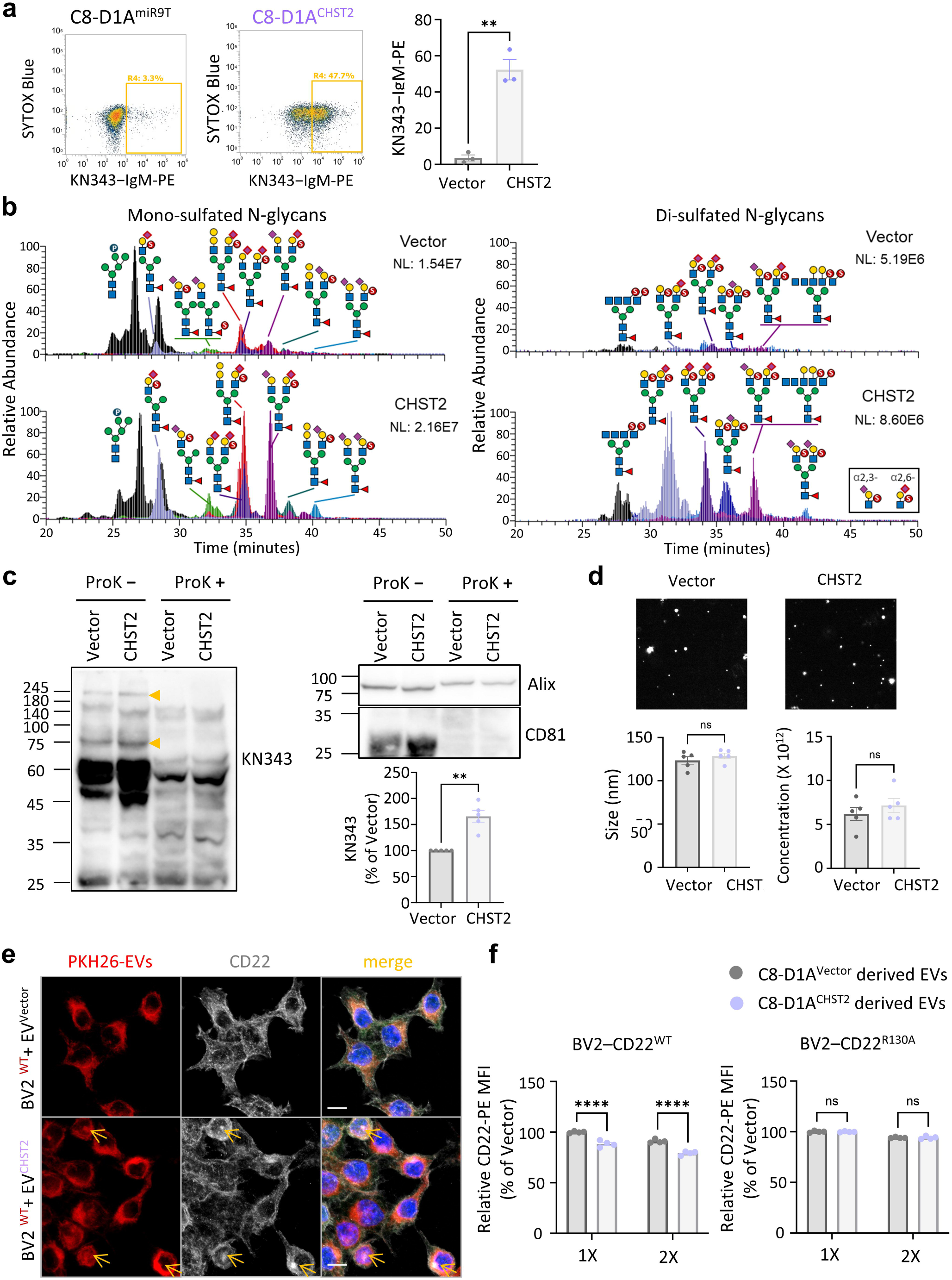
Ligand-enriched extracellular vesicles regulate surface CD22 expression. **(a)** C8-D1A astrocytes with stable CHST2 expression or vector control were generated using a lentiviral system. Flow cytometry analysis using the KN343 antibody confirms that CHST2 overexpression increased α2-6-sialylated 6-sulfo LacNAc expression. Data are presented as mean ± SEM (n = 3). \*\**p* < 0.01 by unpaired *t*-test. (**b**) LC-MS/MS analysis of DMA-derivatized, permethylated sulfated N-glycans derived from the astrocytic cell line expressing either C8-D1A–Vector or C8-D1A–CHST2. Extracted ion chromatograms of representative mono- and di-sulfated N-glycans containing one or two sialic acids in either α2-3 or α2-6 linkage demonstrated a significant increase in N-glycans carrying terminal α2,6-sialylated 6-sulfo-LacNAc in C8-D1A–CHST2 cells. This finding was verified by MS^2^ analysis and by quantifying the summed total intensity of the diagnostic MS^2^ ion at *mlz* 902 (Figure S14) across all data-dependent MS^2^ spectra. (**c**) EVs were isolated from the conditioned medium of C8-D1A–Vector and C8-D1A–CHST2 astrocytes and analyzed by Western blot. EV proteins were probed with the KN343 antibody (yellow arrowhead) and validated using EV markers (Alix and CD81). CHST2 overexpression significantly enhanced the amount of α2-6-sialylated 6-sulfo LacNAc-modified proteins, as indicated by the KN343 signal. Data are presented as mean ± SEM (n = 5). \*\**p* < 0.01 by paired *t*-test. (**d**) NTA analysis showing the size distribution and concentration of EVs. Data are presented as mean ± SEM (n = 5). (**e**) EVs were labeled with PKH26, a red fluorescent lipophilic dye, and incubated with BV2-CD22 cells. Confocal imaging demonstrated that PKH26-labeled EVs were efficiently taken up by BV2-CD22 cells. EVs from C8-D1A–CHST2 astrocytes induce internalization of CD22, as indicated by colocalization of PKH26-EV (red) and CD22 (grey) puncta (yellow arrows). Scale bar = 10 µm. (**f**) BV2-CD22 cells were incubated with EVs for 24 hours, and surface CD22 levels were examined by flow cytometry analysis. EVs from C8-D1A–CHST2 astrocytes reduced surface CD22 in BV2-CD22^WT^ but BV2-CD22^R130A^ cells, as reflected by MFI. Data are presented as mean ± SEM (n = 4). \*\*\*\**p* < 0.0001 by one -way ANOVA.

Since neutralizing CD22 with Cy34.1 enhanced microglial phagocytosis of myelin debris in microglial (Figure 5e-f), and given that myelin breakdown and thinning of the corpus callosum are prominent pathological features in patients and mice with HD [52–56], we evaluated the structural integrity in brains of HD mice using MRI [56, 57]. As expected, HD mice exhibited significant atrophy in multiple brain regions, including the cortex, striatum, corpus callosum, hippocampus, and cerebellum, compared with WT mice (Figure 7d-g, S11a-c), consistent with previous studies [55, 56]. Notably, the absence of CD22 partially restored the volume of the striatum, corpus callosum, and cerebellum in HD mice, suggesting that enhancing microglial phagocytic function may mitigate neurodegeneration [58, 59]. In WT brains, which exhibited no pathology, the absence of CD22 did not affect brain volume (Figure S11 d-f).

Since impaired microglial phagocytosis could prolong neuroinflammation and exacerbate secondary tissue damage during demyelination [58–60], we further evaluated the status of microglia isolated from adult brains of WT and HD mice, with or without CD22, using flow cytometry to analyze a panel of homeostatic and activation markers (Figure S12-13). Levels of homeostatic markers (including CX3CR1, P2RY12, and MerTK [37, 38]) were unchanged in HD mice and unaffected by CD22 deletion (Figure S13a-c). In contrast, microglial activation markers, such as AXL and Gal3 [10, 37, 38], were significantly elevated in HD mice (Figure 7h, S13d). Importantly, CD22 deletion prevented the expansion of Gal3-positive microglia, which are closely associated with pro-inflammatory responses in HD (Figure 7h) [10]. These findings suggest that CD22 deletion may restore microglial functionality, thereby limiting neuroinflammatory cascades and reducing secondary damage. This is consistent with previous studies demonstrating that effective microglial phagocytosis is essential not only for clearing cellular and myelin debris and limiting neuroinflammation but also for establishing an environment conducive to oligodendrocyte-mediated remyelination [58, 59, 61–63].

### Expression of CD22 ligands on astrocytes is impaired by mHTT

Although CD22 suppresses microglial phagocytosis independently of sialic acid binding, this does not rule out the possibility that interactions with ligands may still modulate its activity. To this end, we sought to determine whether natural ligands for CD22 are present in the brain. It is well documented that human CD22 binds both α2,6-linked Neu5Ac (N-acetyl neuraminic acid) and Neu5Gc (N-glycolyl neuraminic acid), whereas mouse CD22 preferentially binds α2,6-linked Neu5Gc [64]. Given that Neu5Gc is rarely found in the brain [65–67] and that sulfation plays an important role in modulating interactions between Siglecs and glycans [68, 69], we hypothesized that potential ligands for CD22 could be sulfated and sialylated glycans. Supporting this hypothesis, a recent study by Jung et al. demonstrated that sulfation of 6-O-GlcNAc by CHST2 (carbohydrate sulfotransferase 2) significantly enhances binding to both human and mouse CD22, suggesting α2-6-sialylated 6-sulfo LacNAc as a potential natural ligand for CD22 in the brain [69]. We next conducted mass spectrometry analysis of PNGase-F-released N-glycans from striatal tissues of WT and HD mice. Reduced N-glycans were selectively dimethylamidated (DMA) prior to permethylation, allowing distinction between permethylated glycans with 2-3 or 2-6-linked sialic acids by a mass difference of 13 u. Subsequent sulfoglycomic analysis of these permethylated sulfated N-glycans revealed that NeuAcα2-6Galβ1-4GlcNAc6S- was indeed present on a range of NeuAc-sialylated complex-type N-glycans (Figure 8a). The presence of the α2-6-sialylated 6-sulfo LacNAc glycotope was further confirmed by m/z values and the diagnostic MS^2^ ion at *mlz* 902 (Figure 8a, S14).

We next performed flow cytometry analysis to quantitatively compare the levels of α2-6-sialylated 6-sulfo LacNAc in various brain cell types from WT and HD mice, using an antibody (KN343) that specifically recognizes this glycan structure [68, 70]. Consistent with the notion that the suppression of *cis*-phagocytosis is independent of sialic acid binding, flow cytometry revealed that α2-6-sialylated 6-sulfo LacNAc is barely detectable on microglia (Figure 8d, S15f). No KN343 signal was detected on oligodendrocytes or endothelial cells either (Figure 8d, S15d-e). In contrast, KN343-positive signals were predominantly present on astrocytes, based on both percentage and mean fluorescence intensity (MFI), and to a lesser extent on neurons (Figure 8d, S15b-c). Importantly, the expression of this ligand was significantly reduced in astrocytes isolated from HD brains (Figure 8c). These findings are further supported by single-nucleus RNA sequencing (snRNA-seq), which showed that astrocytes express high levels of the enzymes ST6GAL1 (*St6gal1*) and CHST2 (*Chst2*), both of which are involved in the synthesis of α2-6-sialylated 6-sulfo LacNAc (Figure S16). In HD, *Chst2* expression is reduced, potentially leading to decreased production of CD22 ligands by astrocytes (Figure 8b), suggesting impaired sialoglycan–Siglec-mediated interactions between astrocytes and microglia.

We hypothesized that ligands expressed on astrocytes may interact in *trans* with CD22 on microglia, either through direct or indirect mechanisms. In the case of direct interaction, ligands on astrocytes may induce clustering of CD22, sequestering it away from ITAM-bearing phagocytic receptors in close molecular proximity and thereby relieving suppression. To test this hypothesis, we generated a stable astrocyte cell line (C8-D1A [71]) overexpressing CHST2, the enzyme responsible for 6-O-sulfation of GlcNAc. Flow cytometry using the KN343 antibody and glycomic analysis confirmed that CHST2 overexpression increased the surface expression of α2-6-sialylated 6-sulfo-LacNAc on C8-D1A cells (Figure 9a-b). However, when BV2-CD22 cells were co-cultured with C8-D1A–CHST2 cells, no CD22 clustering was observed (Figure S19), prompting us to explore the regulation of microglial CD22 by astrocytes through an indirect mechanism.

Earlier studies have shown that astrocytes can modulate the functions of other brain cells by secreting EVs [72–74]. Specifically, ligands on astrocytes may be secreted via EVs, which could promote CD22 internalization. For example, α2,6-sialylated ligand–decorated liposomes have been shown to enhance therapeutic uptake (e.g., doxorubicin) in CD22-expressing B cell lymphoma [46]. We thus hypothesized that the expression of CHST2 may enable astrocytes to release α2-6-sialylated 6-sulfo-LacNAc–decorated EVs to alleviate CD22-mediated suppression of microglial phagocytosis. To assess this possibility, EVs were collected from C8-D1A–CHST2 and C8-D1A–vector cells and analyzed by Western blot. The KN343-positive signals were significantly higher on EVs derived from C8-D1A–CHST2 cells compared with controls (Figure 9c). Nanoparticle tracing analysis (NTA) showed no significant differences in the size and concentration of EVs between the two groups (Figure 9d). EVs were then labeled with the red fluorescent dye PKH26 and incubated with BV2-CD22 cells. Confocal imaging confirmed efficient uptake of PKH26-labeled EVs by the BV2-CD22 cells. Importantly, EVs derived from C8-D1A–CHST2 induced internalization of CD22, as indicated by the colocalization of PKH26-labeled EVs (red) and CD22 (gray) puncta within cells (Figure 9e). We further evaluated CD22 levels on the cell surface following treatment with EVs. Flow cytometry analysis confirmed that treatment with C8-D1A–CHST2-derived EVs significantly reduced surface CD22 levels on BV2-CD22 cells (Figure 9f). These findings support the hypothesis that ligands on astrocytes are delivered via EVs to interact in *trans* with CD22 on microglia, promoting CD22 internalization and thereby modulating microglial phagocytic function.

### Chronic oxidative stress enhances the microglial CD22–astrocytic CHST2 axis in wild-type mice but impairs it in HD mice

To determine whether mHTT triggers the upregulation of microglial CD22, we generated a microglial cell line (BV2) stably expressing HTT_Ex1_-Q_89_ (a mutant HTT fragment harboring an abnormal polyQ expansion) or HTT_Ex1_-Q_19_ (a control HTT fragment with a normal polyQ length) (Figure S20a). Our results showed that expression of mHTT alone did not induce CD22 upregulation (Figure S20b). We then explored potential environmental factors in the brain by exposing cells to various HD-related stimuli, including those that induce inflammatory responses and oxidative stress [75]. Among these tested environmental factors, only oxidative stress specifically elevated CD22 expression (Figure 10a-e). This oxidative stress-mediated upregulation was selective for CD22, as Siglec-E expression remained unaffected (Figure S21). This finding was further confirmed in primary microglia (Figure 10f), supporting the conclusion that oxidative stress specifically drives CD22 upregulation.

**Figure 10.**
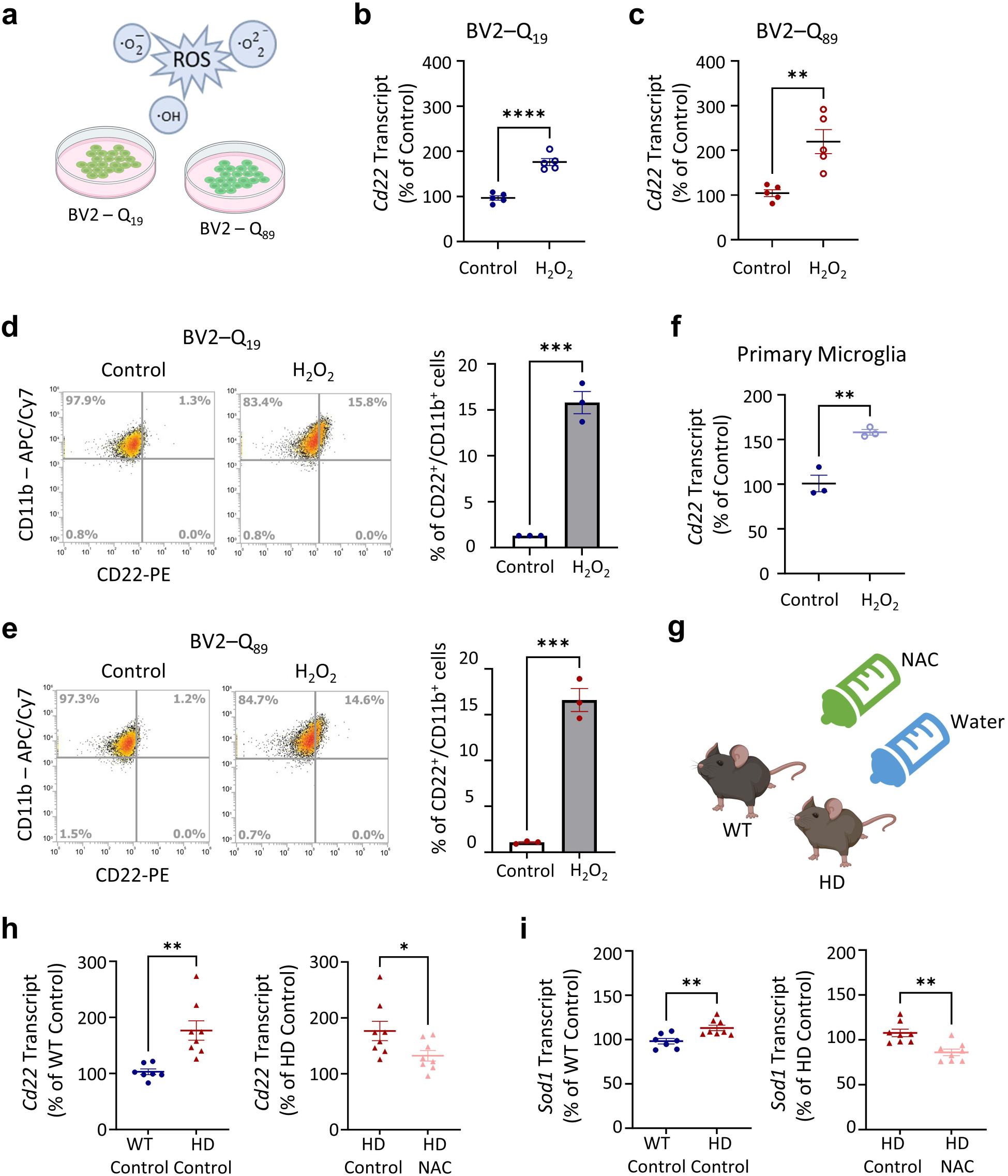
Chronic oxidative stress induces upregulation of CD22. (**a**) BV2 cell lines with stable HTT_Ex1_–Q_19_ (wild-type; designated as BV2–Q_19_) and HTT_Ex1_–Q_89_ (mutant; designated as BV2–Q_89_) expression were exposed to hydrogen peroxide for 24 hours to induce oxidative stress. Exposure to oxidative stress upregulates *Cd22* transcript level in both BV2–Q_19_ (**b**) and BV2–Q_89_ (**c**) cells, as analyzed by qPCR analysis. Data are presented as mean ± SEM (n = 5). Flow cytometry showed an increased population of CD22-immunoreactive microglia in both BV2–Q_19_ (**d**) and BV2–Q_89_ (**e**) cells. Data are presented as mean ± SEM (n = 3). (**f**) Exposure to oxidative stress also upregulates *Cd22* transcript level in neonatal primary microglia. Data are presented as mean ± SEM (n = 3 biological replicates). (**g**) Wild-type (WT) and HD mice were fed with NAC in the drinking water from 7 to 13 weeks of age to mitigate oxidative stress accumulation, and brain tissues were collected at 13 weeks of age. Transcript levels of *Cd22* (**h**) and *Sod1* (**i**) in the striatum were examined by qPCR analysis. Data are presented as mean ± SEM (n = 7-8 animals per group). \**p* < 0.05, \*\**p* < 0.01, \*\*\**p* < 0.001 and \*\*\*\**p* < 0.0001 by unpaired *t*-test.

This finding is particularly important, as elevated oxidative stress has been previously reported in HD patients and HD mice [76–79]. To further confirm that CD22 upregulation in the brain is mediated by oxidative stress, we treated HD mice with an antioxidant (N-acetyl cysteine, NAC; administered in drinking water) from 7 weeks of age (pre-symptomatic stage) to 13 weeks (late stage) to reduce systemic oxidative stress [76]. Importantly, NAC treatment effectively prevented CD22 upregulation (Figure 10h), supporting the role of oxidative stress in driving CD22 expression. In control brains, which exhibited normal levels of oxidative stress, NAC treatment did not affect the expression of *Cd22* or *Sod1*, confirming the selectivity of the NAC treatment. (Figure S22).

We next evaluated how the level of the ligand for CD22 is regulated under similar conditions. Since mutant HTT can also be detected in astrocytes [80], we assessed whether the expression of mHTT affects the levels of 6-sialylated 6-sulfo LacNAc. To this end, we generated stable C8-D1A cell lines expressing HTT_Ex1_-polyQ_89_ or HTT_Ex1_-Q_19_. Our results indicate that the presence of mHTT alone does not lead to the downregulation of ST6GAL1 (*St6gal1*) or CHST2 (*Chst2*) (Figure S23). However, upon exposure to oxidative stress, *Chst2* expression was significantly upregulated in C8-D1A cells expressing HTT_Ex1_-Q_19_ (C8-D1A–Q_19_), but not in those expressing HTT_Ex1_-Q_89_ (C8-D1A–Q_89_) (Figure 11a). Consequently, *Chst2* expression was significantly lower in the presence of mHTT. Flow cytometry analysis further confirmed that the percentage of KN343-positive cells was also significantly reduced in C8-D1A cells expressing mHTT (HTT_Ex1_-Q_89_) compared to those expressing HTT_Ex1_-Q_19_ under elevated oxidative stress (Figure 11b). Consistent results were observed in primary astrocytes isolated from wild-type and HD mice (Figure 11c). In line with this, we found that *Chst2* expression was effectively restored in HD mice treated with NAC (Figure 11d). No changes in the level of *St6gal1* were observed with the expression of mutant HTT or under elevated oxidative stress (Figure 11a, c). Collectively, these findings suggest that astrocytes may upregulate CD22 ligand production to modulate microglial phagocytosis in response to oxidative stress. However, this regulatory mechanism is impaired by the presence of mHTT, disrupting astrocyte-mediated support of microglial function under conditions of elevated oxidative stress.

**Figure 11.**
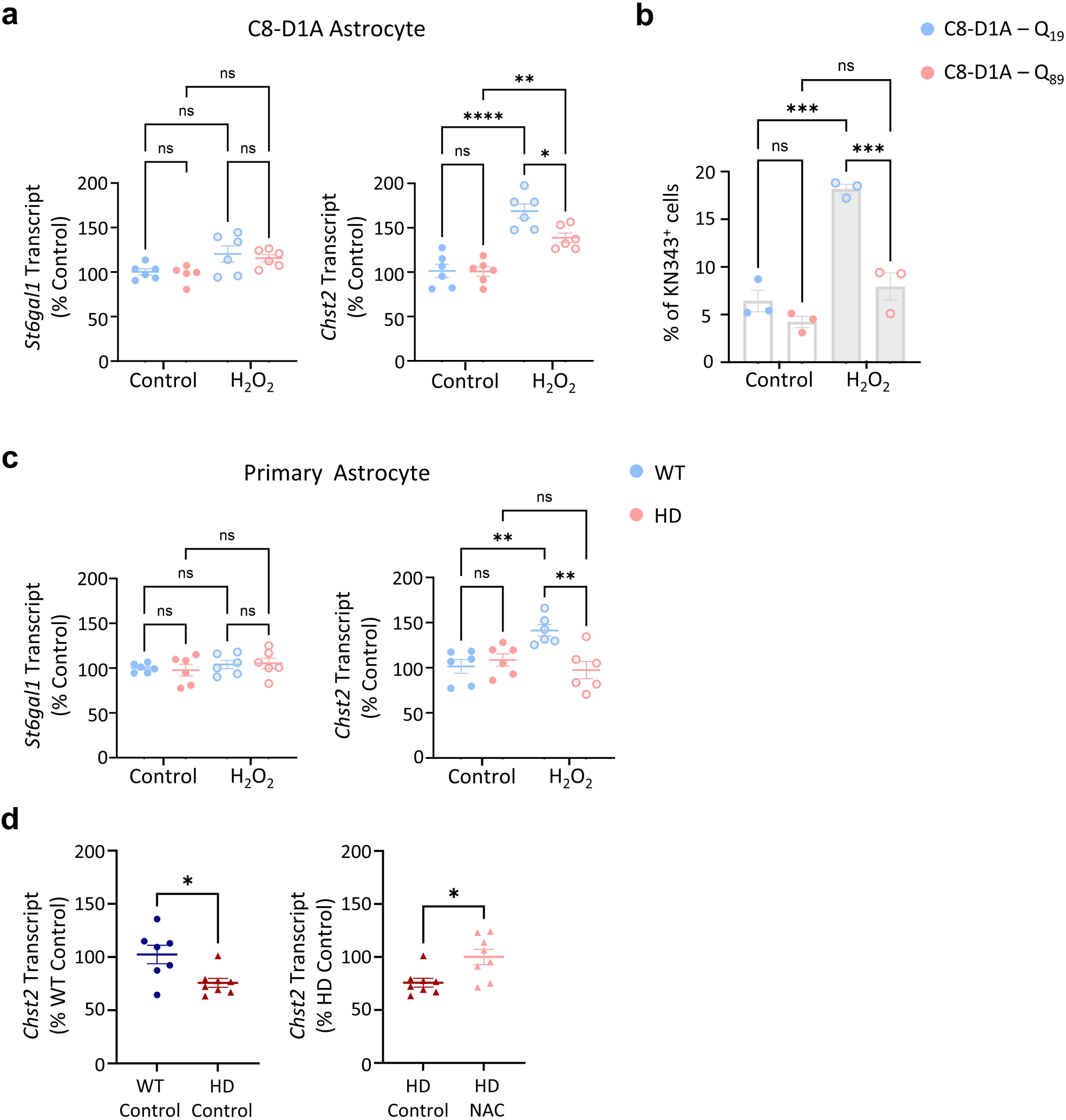
Presence of mHTT disrupts CHST2 expression in astrocytes under oxidative stress. C8-D1A cell lines with stable HTT_Ex1_–Q_19_ (wild-type; designated C8-D1A–Q_19_) and HTT_Ex1_–Q_89_ (mutant; designated C8-D1A–Q_89_) expression were exposed to hydrogen peroxide for 24 hours to induce oxidative stress. (**a**) Transcript levels of *St6gal1* and *Chst2*, enzymes involved in the synthesis of α2-6-sialylated 6-sulfo LacNAc, were examined by qPCR analysis. Exposure to oxidative stress upregulates *Chst2* in C8-D1A–Q_19_ but not C8-D1A–Q_89_ cells. Data are presented as mean ± SEM (n = 6). (**b**) Flow cytometry using the KN343 antibody demonstrated increased α2-6-sialylated 6-sulfo LacNAc (KN343⁺) signal following oxidative stress in C8-D1A–Q_19_, but not in C8-D1A–Q_89_ cells. Data are presented as mean ± SEM (n = 3). (**c**) Primary astrocytes derived from WT and HD mice were exposed to hydrogen peroxide for 24 hours to induce oxidative stress. Exposure to oxidative stress upregulates *Chst2* in WT but not HD-derived primary astrocytes. Data are presented as mean ± SEM (n = 6 biological replicates). (**d**) Transcript level of *Chst2* in the striatum of WT and HD mice fed with NAC revealed that *Chst2* was significantly downregulated in HD mice and restored by NAC treatment. Data are presented as mean ± SEM (n = 7-8 animals per group\**p* < 0.05, \*\**p* < 0.01, \*\*\**p* < 0.001 and \*\*\*\**p* < 0.0001 by unpaired *t*-test or two-way ANOVA.

## Discussion

The diversity of glycan structures on the mammalian cell surface plays a crucial role in regulating cell functions and interactions [81]. This complex glycan code is often interpreted by a range of glycan-binding proteins, such as Siglecs and galectins, which are expressed on immune cells, including microglia {the tissue resident macrophages in the brain) [82, 83]. Among these, sialic acids at the termini of glycans and their cognate receptors Siglecs are particularly noteworthy. However, the roles of sialoglycans and Siglecs in neurological disorders, especially HD, remain largely underexplored. In the present study, we identified CD22 upregulation in microglia as a driver of impaired phagocytosis in HD, mediated through ITIM–ITAM signaling and exacerbated by oxidative stress. Blocking CD22, either genetically or with a blocking antibody or ligand-coated EVs, restored microglial phagocytosis, improved motor function, reduced mHTT inclusions, and alleviated brain atrophy in HD mice. These findings uncover CD22 as an immune checkpoint that disrupts astrocyte–microglia communication, thereby presenting a potential therapeutic target in HD.

Our global profiling of Siglecs in the striatum of both human and mouse brains with HD provided insightful findings (Figure 1-2; Table S1-S2; Figure S1, S6). In the human caudate nucleus, aside from those presented in Figure 1a, we found no detectable expression of Siglec-1 and Siglec-7, consistent with previous reports of their absence in the cerebral cortex [84, 85]. Conversely, Siglec-10, which is highly expressed in the cerebral cortex, was undetectable in the caudate nucleus [84, 85]. In mice, Siglecs are mainly expressed in microglia, and at least eight different Siglecs could be detected. Mass cytometry analysis revealed differential expression of Siglecs across different microglial subpopulations (Figure 2). For instance, Siglec-H, a molecular signature for microglia, was broadly expressed across microglial subpopulations. In addition, CD33 and Siglec-E appeared in multiple subpopulations. In contrast, CD22, Siglec-F, and Siglec-G showed more restricted expression profiles. These observations underscore that individual Siglecs may regulate microglial functions in distinct subpopulations by interacting with region- or cell type-specific ligands. These findings support the growing recognition of microglial heterogeneity, whereby individual microglial subpopulations express distinct receptors and molecules that likely reflect specialized functions.

Among these, CD22 stood out due to its consistent and disease stage-dependent upregulation in the striatum of both HD patients and HD mice (Figure 1b), prompting a deeper investigation into its functional role. Our findings indicate that aberrant CD22 upregulation suppresses the phagocytic clearance function of microglia, aligning with an earlier finding of Pluvinage et al. [16], who first identified CD22 as a negative regulator of microglial phagocytosis in aging. Our study extends the findings of Pluvinage et al. by providing evidence that CD22 suppresses phagocytosis via sialic acid-independent ITIM-ITAM crosstalk (Figure 3-5). It also supports a model in which proximal ITIM-ITAM signaling, rather than direct ligand competition, underlies the ability of CD22 to antagonize ITAM-bearing phagocytic receptors (Figure 4, S24) [33, 55]. Our findings are consistent with the canonical role of ITIM-bearing receptors antagonizing ITAM-mediated signaling [43, 86, 87] and extend the established paradigm of CD22-mediated inhibition beyond BCR signaling.

Microglia are known to express a wide variety of phagocytic receptors that facilitate the clearance of diverse materials in the brain. Our study specifically focuses on the inhibitory interaction between CD22 and Dectin-1, providing mechanistic insight into how aberrant upregulation of CD22 may impair microglial phagocytic function during neurodegeneration (Figure 3-5). In addition to Dectin-1 (*Clec7a*), microglia also rely on other ITAM-bearing receptors, such as TREM2 and AXL, to mediate phagocytosis [60, 87, 88]. Notably, these phagocytic receptors are strongly upregulated in disease-associated microglia (DAM) or microglial-neurodegenerative phenotype (MGnD), specialized subpopulations that emerge in response to neurodegenerative stimuli [37, 38]. While such upregulation of phagocytic receptors in these subpopulations may reflect a compensatory mechanism to enhance clearance [36, 60, 88, 89], our findings suggest that the concurrent upregulation of CD22 in microglia may undermine this adaptive response. By engaging in ITIM-ITAM signaling crosstalk, CD22 may suppress the activity of these ITAM-bearing phagocytic receptors, thereby limiting the effectiveness of microglial clearance during neurodegeneration. This hypothesis aligns with recent studies showing that human CD33, another ITIM-bearing Siglec, inhibits TREM2-mediated phagocytosis of amyloid-beta through a similar ITIM-ITAM signaling mechanism [90, 91].

Mass cytometry analysis highlighted a detrimental role for CD22⁺ microglia, showing their association with upregulation of AXL (a marker of DAM) and concomitant downregulation of TMEM119 and P2RY12 (markers of homeostatic microglia) in cluster 8 (Figure 2f) [37, 38]. Multiple lines of evidence from HD mouse models further support a pathogenic role for CD22, as its genetic deletion ameliorates HD pathology and improves motor performance (Figure 6b). Consistent with these, recent reports demonstrated impaired microglial phagocytic function in late-stage HD patients and HD mice, to which the CD22 upregulation observed here may contribute to this defect [49, 92]. Importantly, we demonstrate that genetic removal of CD22 restores phagocytic function in microglia, as evidenced by in vitro phagocytosis assays (Figure 5f) and in vivo recovery of the phagocytic marker CD68 (Figure 7c). Most critically, restoration of microglial phagocytic capacity by genetically removing CD22 improved pathological outcomes in HD mice, including reduced mHTT inclusions, increased Darpp-32 expression, and attenuated brain degeneration (Figure 6c-f, 7d-g).

We further identified α2,6-sialylated-6-sulfo-LacNAc as a primary CD22 ligand in the brain, with astrocytes as its primary source (Figure 8), and demonstrated that it can be delivered to microglia via astrocyte-derived EVs (Figure 9). EVs are heterogeneous vesicles released by cells into the extracellular spaces to mediate paracrine or long-range communication. They are functionally diverse, carrying distinct surface antigens, as well as protein and nucleic acid cargos [93]. In the brain, glial cells are known to actively release EVs, and oxidative stress may further enhance their secretion [94, 95], suggesting that oxidative stress-induced EV release could represent a mechanism likely relevant in chronic neurodegenerative conditions such as HD. Previous studies have shown that astrocytic EVs may modulate microglial activity through miRNA cargos. For example, morphine-stimulated astrocytes were shown to release EVs enriched with miR-138, which activate microglia via the TLR7–NF-κB pathway [73], whereas astrocytic EVs containing miR-873a-5p promote microglial de-activation and attenuate neuroinflammation following traumatic brain injury [96]. These findings highlight the importance of EV-mediated astrocyte-microglia communication in acute injury models. In contrast, our study identifies a fundamentally distinct mechanism operating under oxidative stress or chronic neurodegeneration. Specifically, astrocytic EVs carry α2,6-sialylated-6-sulfo-LacNAc, a glycan ligand that induces CD22 endocytosis in microglia, releasing ITAM-bearing phagocytic receptors from CD22-mediated inhibition. This glycan-Siglec interaction functions as an immune checkpoint controlling microglial phagocytosis, and its disruption by mHTT and oxidative stress contributes to impaired clearance in HD. Our findings revealed a previously unrecognized layer of astrocyte-microglia communication during HD progression. This mechanism mirrors observations with CD33, where engagement with sialylated ligand-decorated liposomes or anti-CD33 blocking antibodies reduced CD33 surface expression and restored amyloid-beta uptake [97–99].

Oxidative stress emerged as a key upstream regulator of this aberrant regulation. We identified that oxidative stress specifically induces CD22 but not other Siglecs (such as Siglec-E), while simultaneously suppressing its ligand-synthesizing enzyme CHST2. Treatment with the antioxidant NAC prevented both changes (Figures 10h, 11d). These findings are important because chronic oxidative stress is a well-documented feature of neurodegenerative diseases, including HD and Alzheimer’s disease (AD). Similar to HD, the levels of CD22 transcripts were also significantly elevated in the hippocampus of two AD mouse models at late disease stages (APP/PS1 and THY-Tau22, 13 months; Figure S25)[100, 101], further supporting the pathological relevance of CD22 in neurodegenerative diseases.

## Conclusions

Ample evidence suggests that changes in the cell surface glycan code or in the expression of glycan-binding proteins can disrupt microglial communication with neighboring cells. These alterations may be driven by genetic factors linked to neurodegenerative diseases (such as AD and HD) and/or environmental stressors (including inflammation and oxidative stress). Such disruptions in microglial signaling can potentially exacerbate disease progression. Conversely, restoring proper microglial signaling in a timing-and context-dependent manner is critical for preserving brain homeostasis [10, 16, 31, 49, 51, 59, 91, 97]. Our findings demonstrate that astrocytes can modulate the phagocytic clearance function of microglia via Siglecs in response to oxidative stress. This regulatory mechanism is disrupted in HD. Our study uncovered a previously unappreciated orchestration of Siglecs involved in regulating microglial function and brain homeostasis. Further investigations on how diverse Siglecs and their glycan ligands contribute to immunomodulation may offer new therapeutic avenues for treating neurodegenerative diseases.

## Abbreviations

AD: Alzheimer’s disease
CHST2: Carbohydrate Sulfotransferase 2
HD: Huntington’s disease
EV: Extracellular vesicles
HTT: Huntingtin
ITIM: Immunoreceptor tyrosine-based inhibitory motif
ITAM: Immunoreceptor tyrosine-based activatory motif
K0: Knockout
MRI: Magnetic Resonance Imaging
NAC: N-acetyl cysteine
NTA: Nanoparticle Track Analysis
PLA: Proximity ligation assay
RT-qPCR: real-time quantitative polymerase chain reaction
SHP: Src homology 2 domain-containing protein tyrosine phosphatase
Siglee: Sialic acid-binding immunoglobulin-type lectin
snRNA-seq: Single nuclei RNA sequencing
ST6GALJ: Beta-Galactoside Alpha-2,6-Sialyltransferase 1
WT: Wild-type

## Ethics Approval

Anonymized human brain tissue specimens were obtained from the NIH NeuroBioBank (USA) in accordance with all applicable ethical guidelines and regulations. No additional informed consent was required as the samples were fully de-identified. Demographic information for the specimens used in RNA preparation is summarized in Table S1. All animal experimental procedures were performed in accordance with the guidelines established by the Institutional Animal Care and Use Committee (IACUC) of the Institute of Biomedical Sciences at Academia Sinica.

## Data Availability

RNA data SRA files will be deposited in the NCBI’s Sequence Read Archive and be released to the public upon publication. All supporting information and data are available in the article and supplementary files.

## Acknowledgements

We thank Dr. Yao-Ming Chang and Dr. Tai-Ming Ko in technical support and advice on RNA sequencing. We would like to thank the Flow Cytometry Core Faciltiy (Core Facility and Innovative Instrument Project AS-CFII-111-212) of the Institute of Biomedical Sciences, Academia Sinica for cell sorting and technical support. We thank the Light Microscopy Core Facility of the Institute of Biomedical Sciences, Academia Sinica for the technical support in image acquisition and analysis. We thank the Animal Imaging Facility, Academia Sinica and Taiwan Animal Consortium for the technical support in the *in vivo* MRI analysis of the mice brain. We thank the Academia Sinica Common Mass Spectrometry Facilities for Proteomics and Protein Modification Analysis (AS-CFII-111-209) for MS data acquisition. We thank the Core Facilities of Translational Medicine of BioTReC (National Biotechnology Research Park, Academic Sinica) for the technical support in NTA analysis. The figure abstract and icons were created with BioRender.com.

## Authors’ contributions

**YH Lee**: contributed to the conception and execution of the research project and wrote the first draft of the manuscript.

**JJ Siew**: contributed to cDNA preparation from human brain samples and analysis of the snRNA-seq data from mouse brain samples

**CW Lee**: contributed to sample preparation for mass cytometry and snRNA-seq analyses

**HM Chen**: contributed to animal breeding and assisted with behavioral analyses

**D Sridharan, YT Lu, PCJ Huang, YF Wang, SY Chen**: contributed to the design, optimization, validation and analysis of the mass cytometry panel

**SYGuu, PY Wang, HC Chang, KH Khoo**: contributed to the preparation and mass spectrometry-based analysis of N-glycans from mouse brain tissue

**SY Liang**: assisted in the analysis of scRNA-seq and snRNA-seq data from mouse brain samples and human public datasets

**T Angata, Y Chern**: contributed to the conception of the research project and provided critical guidance throughout the project’s execution and manuscript preparation

