## Supplementary material for "Oxidative Stress-Induced Microglial CD22 Upregulation Impairs Phagocytosis and Exacerbates Huntington’s Disease": Figure S

**Supplementary Table 1** **Brain tissue specimens from Human Brain and Spinal Fluid Resource Center for RNA preparation and qPCR analysis.**

| Specimen No. | Neuropathological Diagnosis | Clinical Diagnosis | Brain Region | Age | Gender | Post-Mortem Interval (hr) |
|---|---|---|---|---|---|---|
| #2691 | Huntington's Disease; not determined | Huntington's Disease; Depression | Caudate nucleus | 65 | Female | 13.0 |
| #2706 | Huntington's Disease; Vonsattel Grade 1-2 | Huntington's Disease; Dementia | Caudate nucleus | 43 | Male | 17.0 |
| #3091 | Huntington's Disease; Vonsattel Grade 4 | Huntington's Disease | Caudate nucleus | 44 | Female | 9.0 |
| #3358 | Huntington's Disease; Vonsattel Grade 1-2 | Huntington's Disease | Caudate nucleus | 63 | Male | 8.7 |
| #3881 | Huntington's Disease; Vonsattel Grade 2 | Huntington's Disease | Caudate nucleus | 65 | Male | 9.1 |
| #3805 | Normal brain | Acute renal failure; Diabetes Mellitus | Caudate nucleus | 70 | Male | 12.0 |
| #4332 | Normal brain | Colon cancer with liver metastasis | Caudate nucleus | 63 | Male | 12.0 |
| #4331 | Normal brain | Myocardial infarction | Caudate nucleus | 68 | Female | 23.7 |
| #4631 | Normal brain | Chronic obstructive pulmonary disease; Emphysema; Congestive heart failure; Hypertension | Caudate nucleus | 59 | Male | 20.2 |
| #4716 | Minimal changes consistent with aging | Dementia; Seizure disorder | Caudate nucleus | 55 | Female | 19.7 |

### Supplementary Figure 1

#### CD22 expression across brain cell types in mouse and human

a

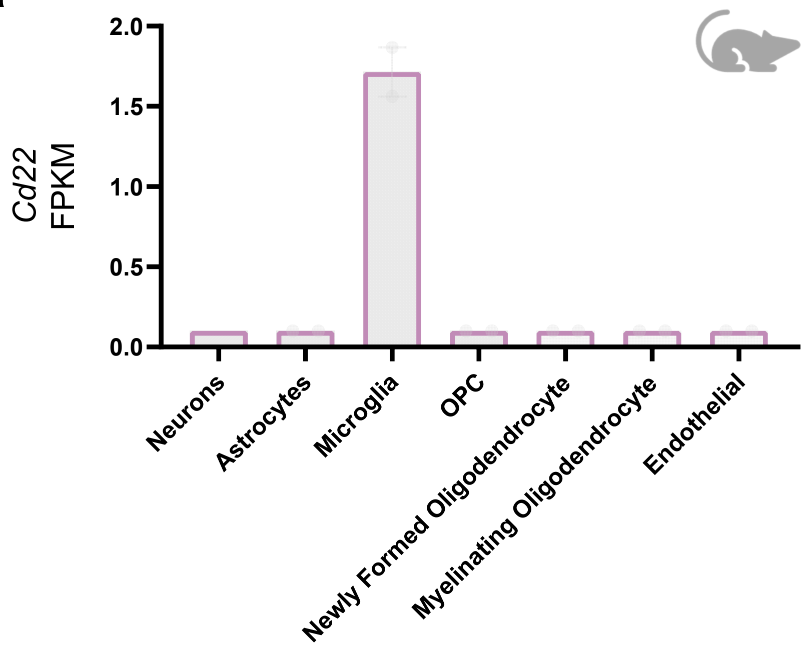


b

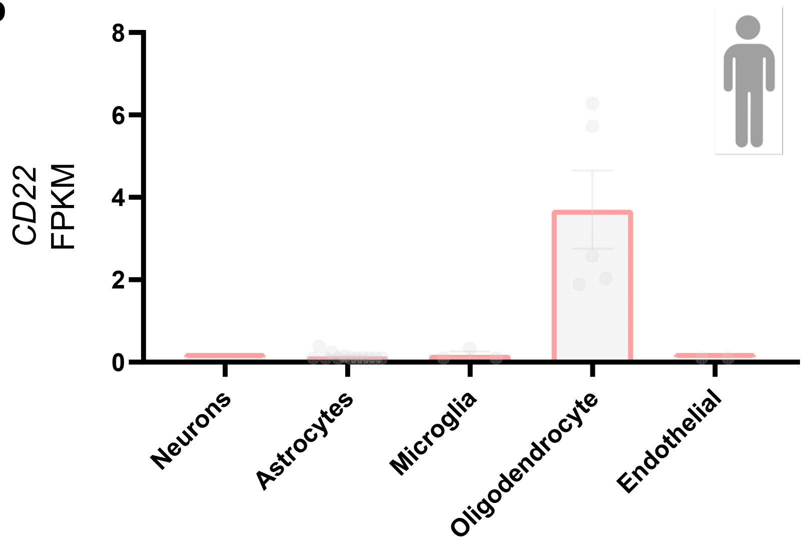

**Supplementary Table 2** **Expression of *CD22* in different cell type of the caudate nucleus of HD subjects.**

Data were retrieved from snRNA-seq dataset from Lee et al.

| Dataset ID | Region | Cell Type | Human Caudate Control $log_2avg$ | Human Caudate HD $log_2avg$ | $log_2FC$ | *p* value | *p* adjusted |
|---|---|---|---|---|---|---|---|
| GSE152058 | Caudate | Microglia | 1.74 | 5.72 | 3.8 | 4.15E-05 | 2.73E-03 |
| GSE152058 | Caudate | Oligodendrocytes | 11.18 | 11.69 | 0.351 | 6.75E-02 | 2.08E-01 |

### Supplementary Figure 2

#### Traditional flow cytometry confirms expansion of CD22-immunoreactive microglia in HD mice

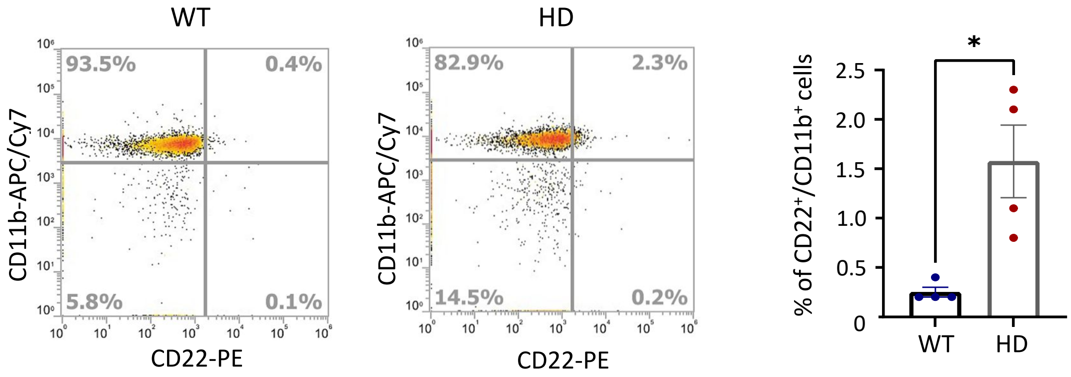

### Supplementary Figure 3

#### Gating hierarchy of FlowSOM

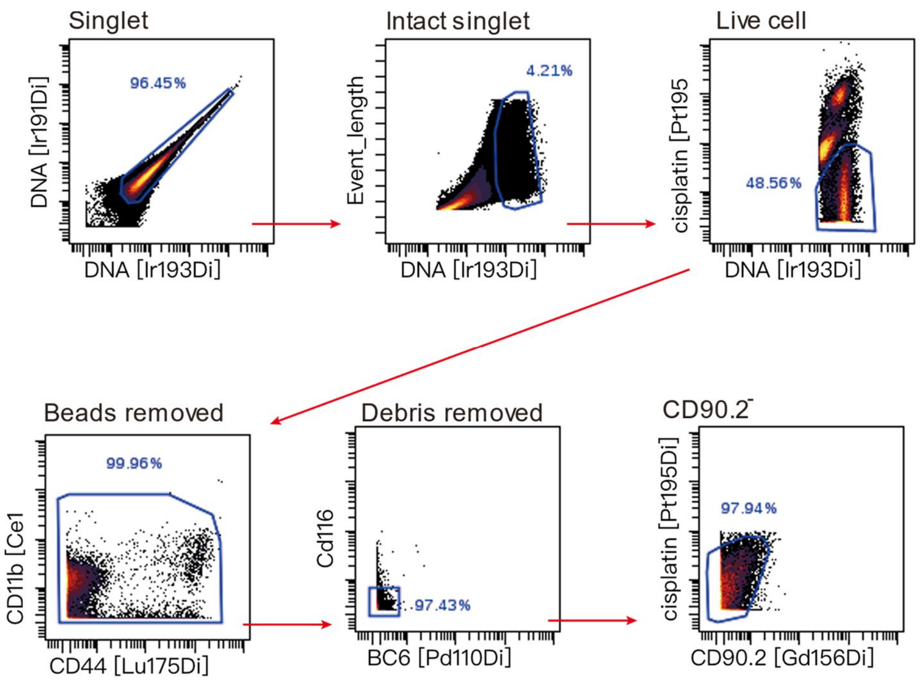

### Supplementary Figure 4 First UMAP

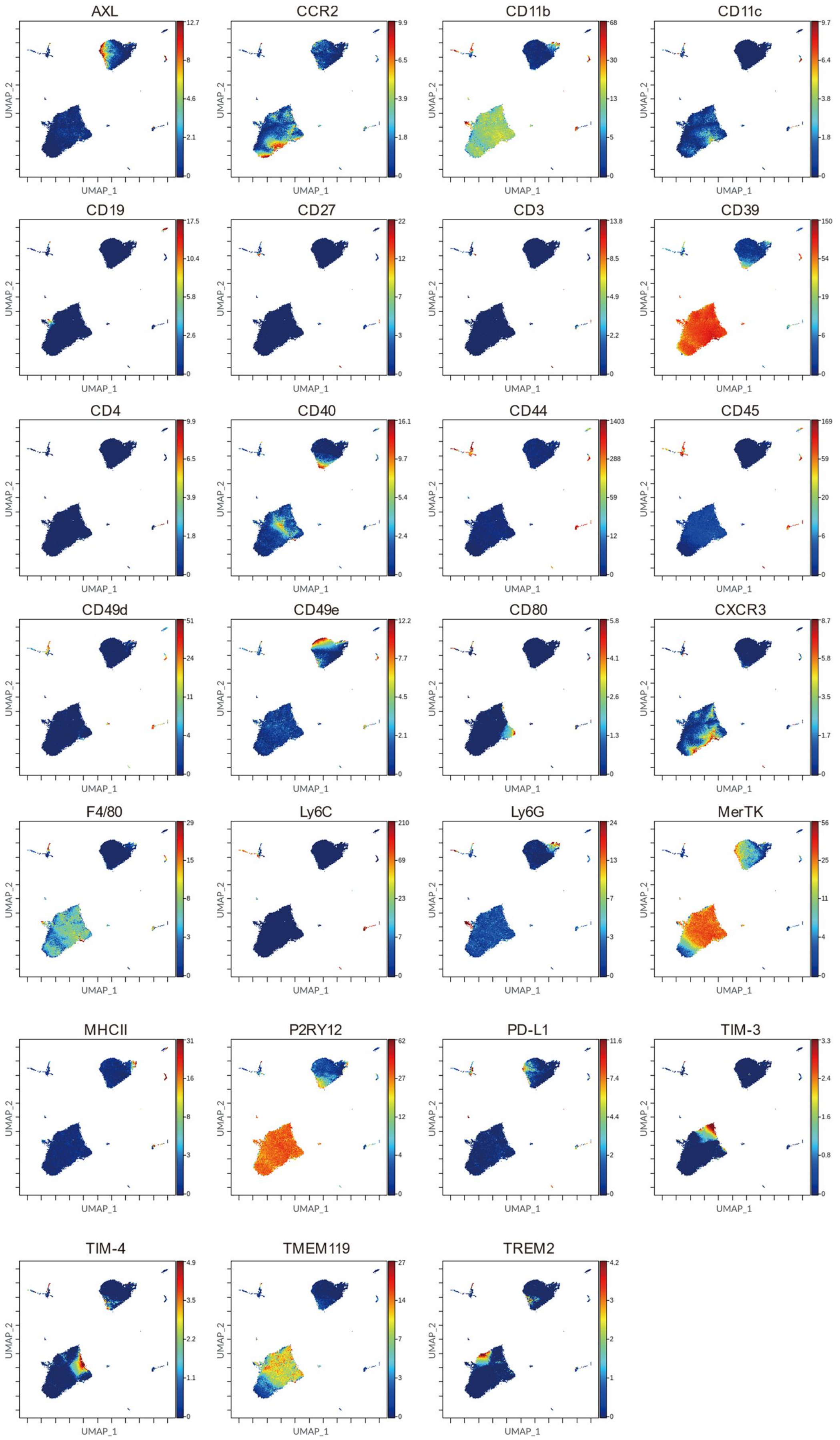

### Supplementary Figure 5

#### Population percentages of FlowSOM-identified metacluster

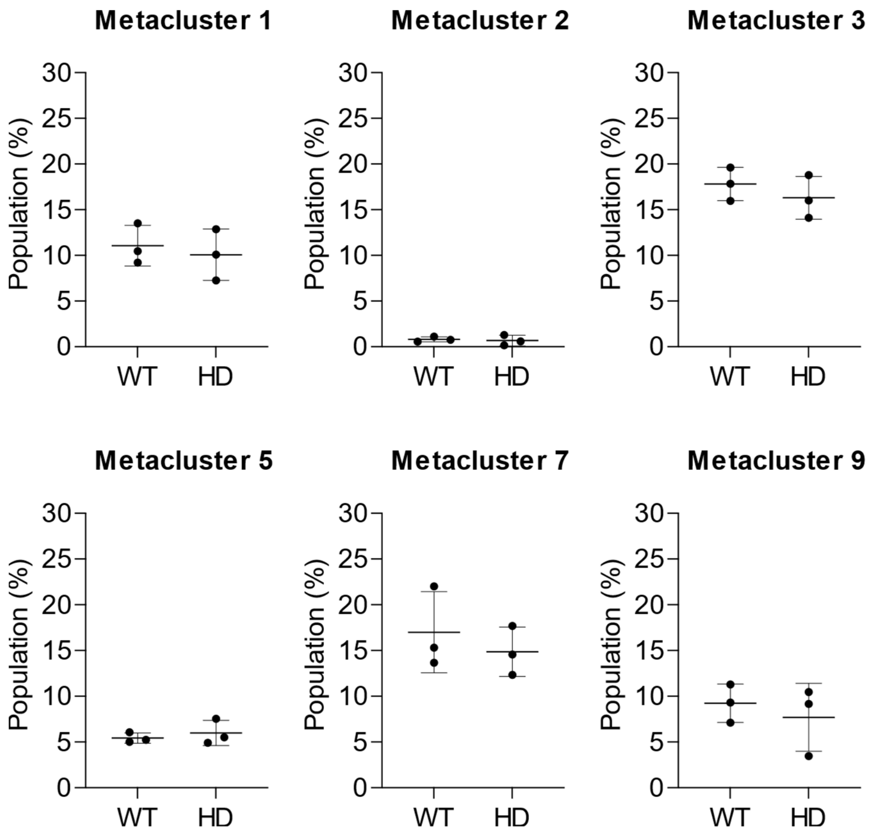

### Supplementary Figure 6

#### Second UMAP

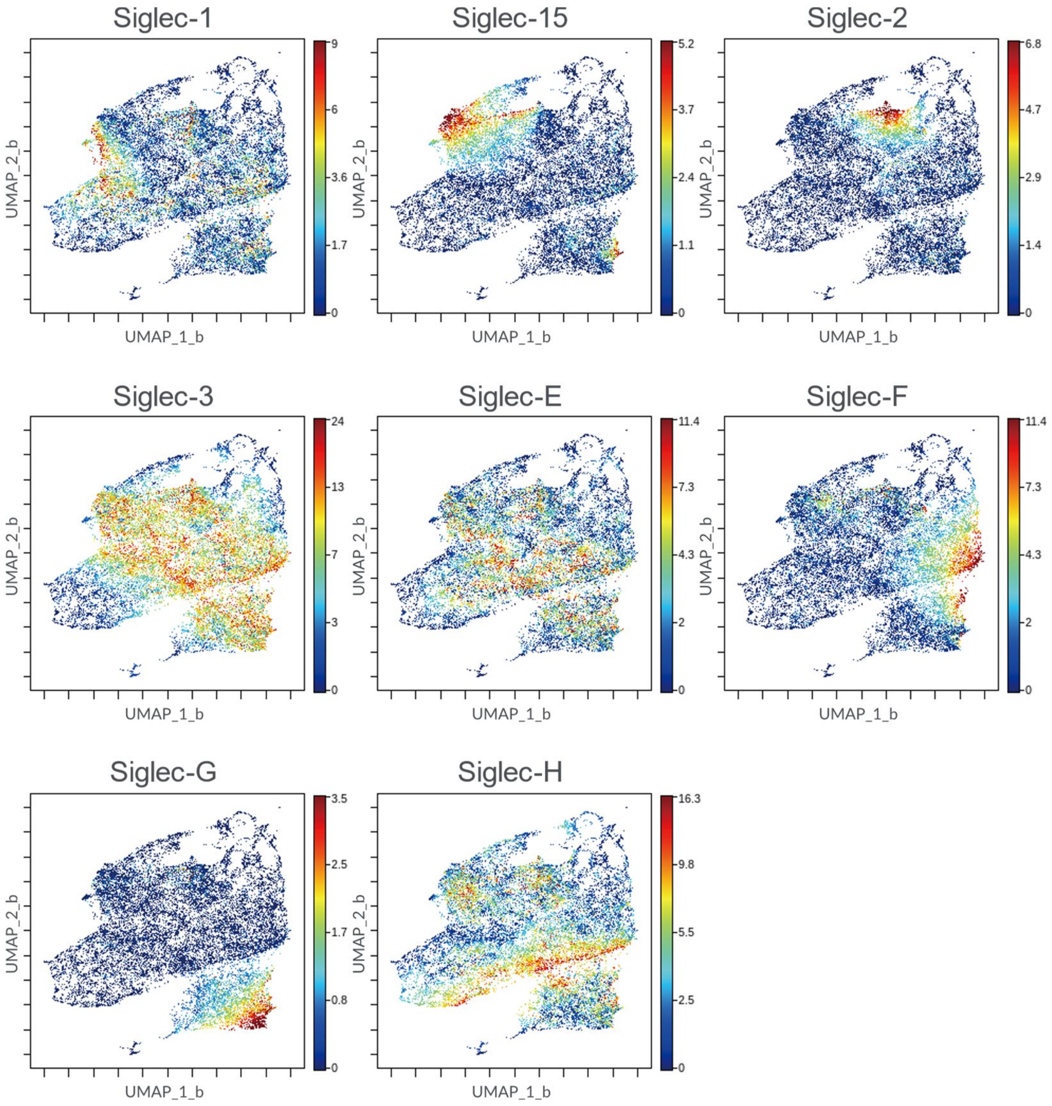

### Supplementary Figure 7

#### CD22 does not affect the inflammatory or oxidative stress response of microglia

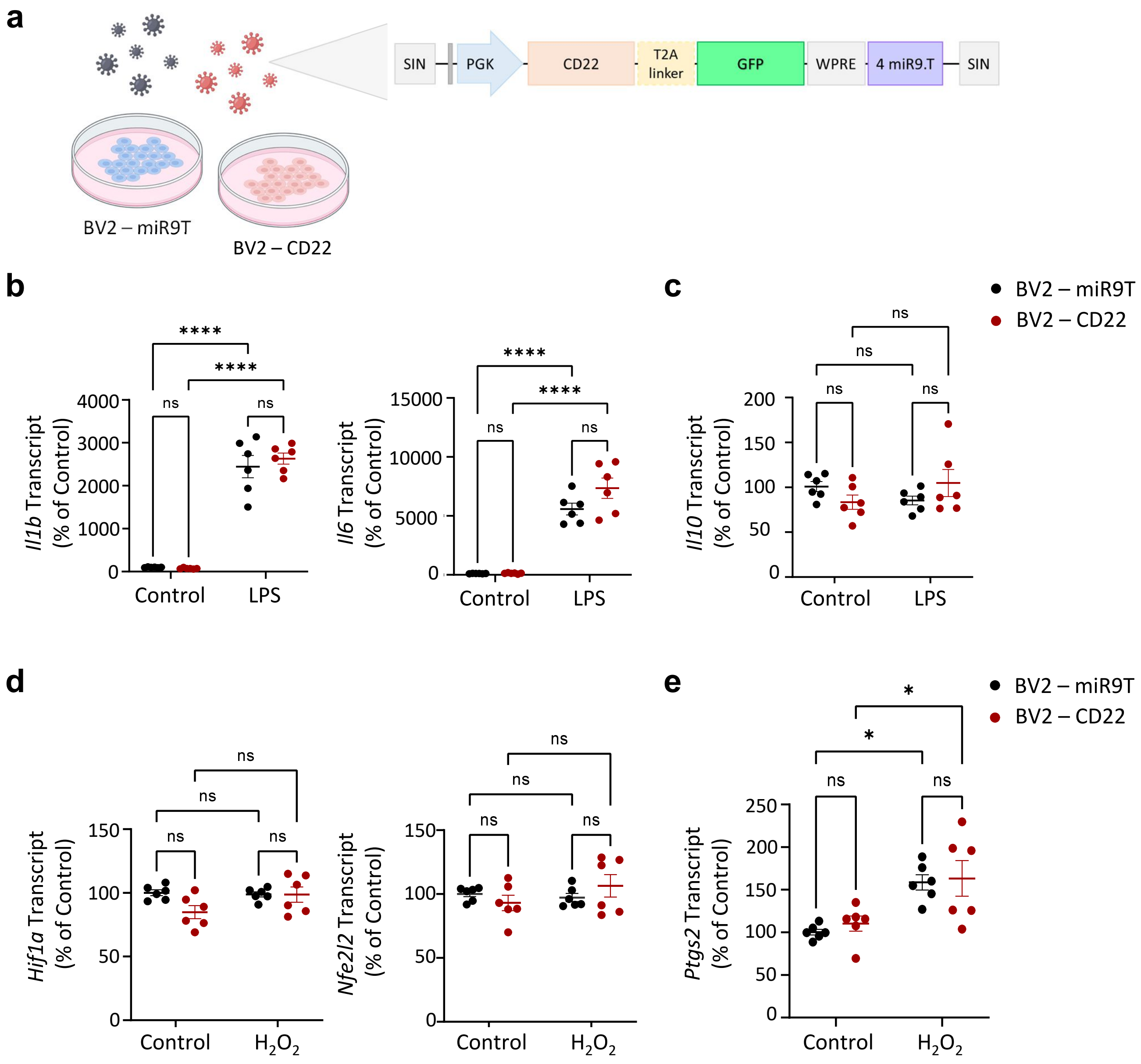

### Supplementary Figure 8

#### CD22 is co-localized with *Clec7a*-V5

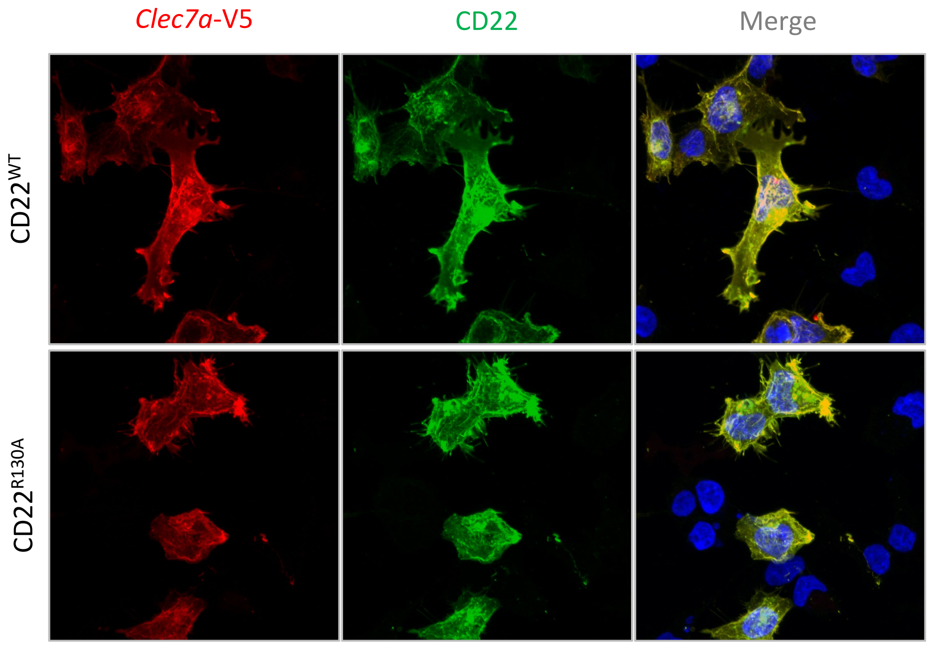

### Supplementary Figure 9

#### Knockout of CD22 did not affect body weight, limb clasping or life span

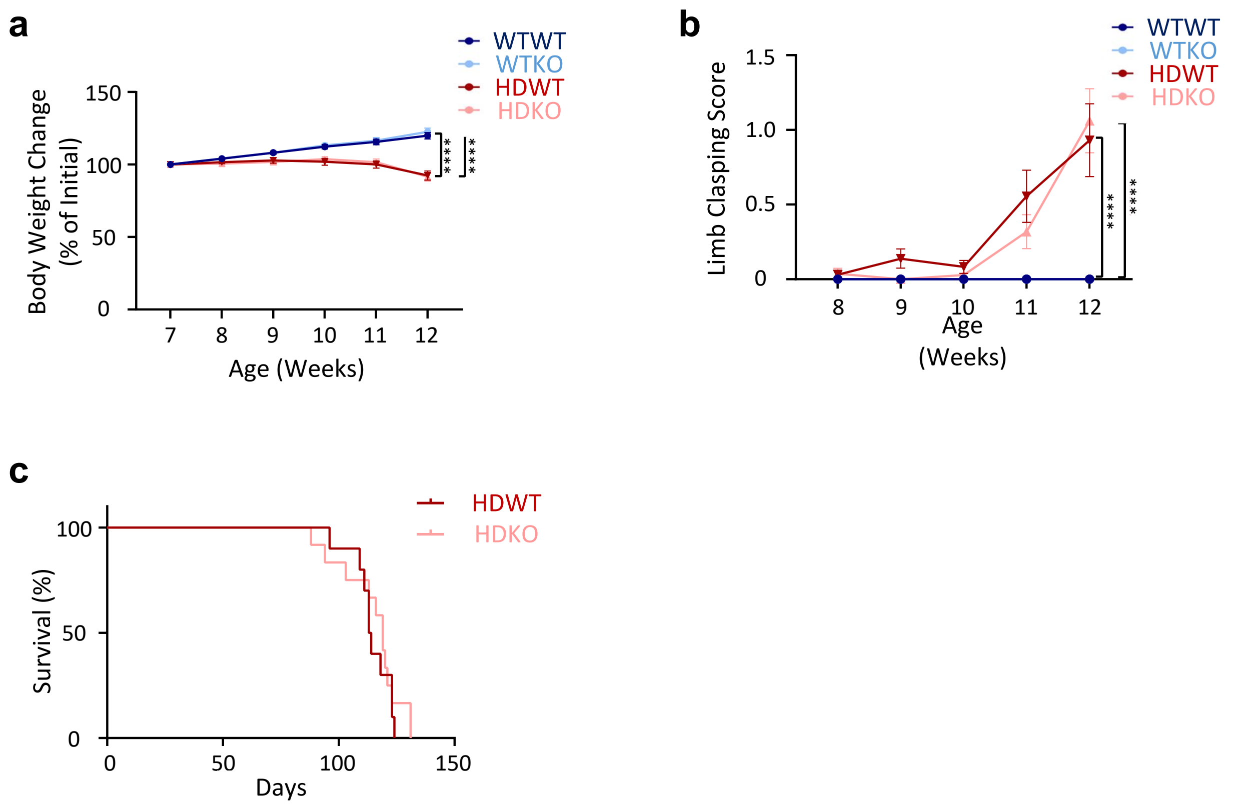

### Supplementary Figure 10

#### *Cd68* is significantly down-regulated in HD striatum

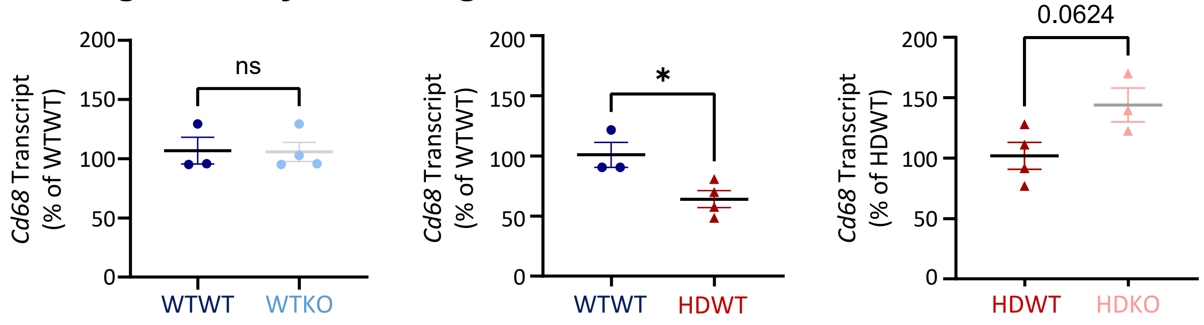

### Supplementary Figure 11

#### Knockout of CD22 does not affect the volume of the whole brain, cortex and hippocampus

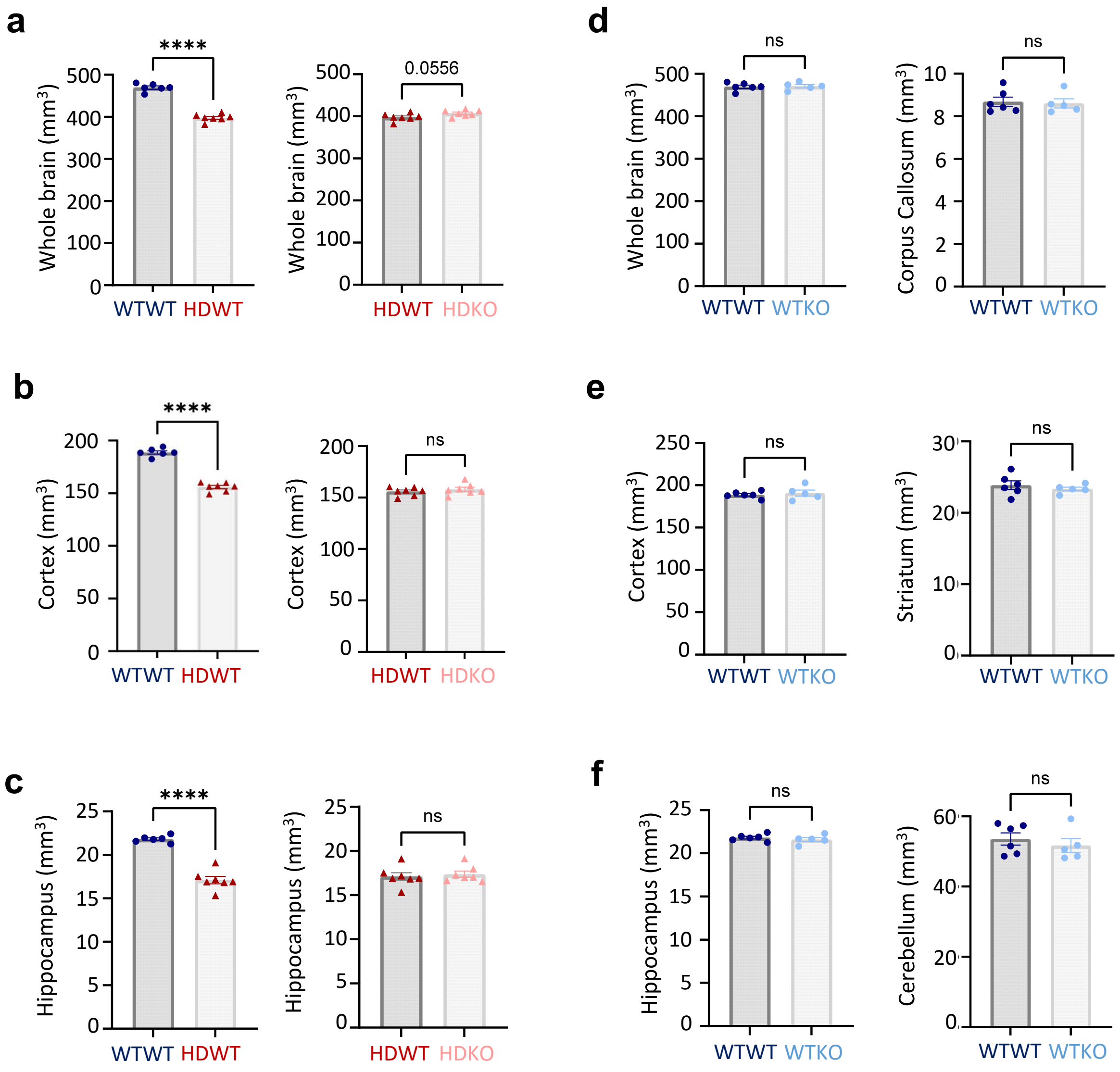

### Supplementary Figure 12

#### Gating strategy for CD11b+ microglia

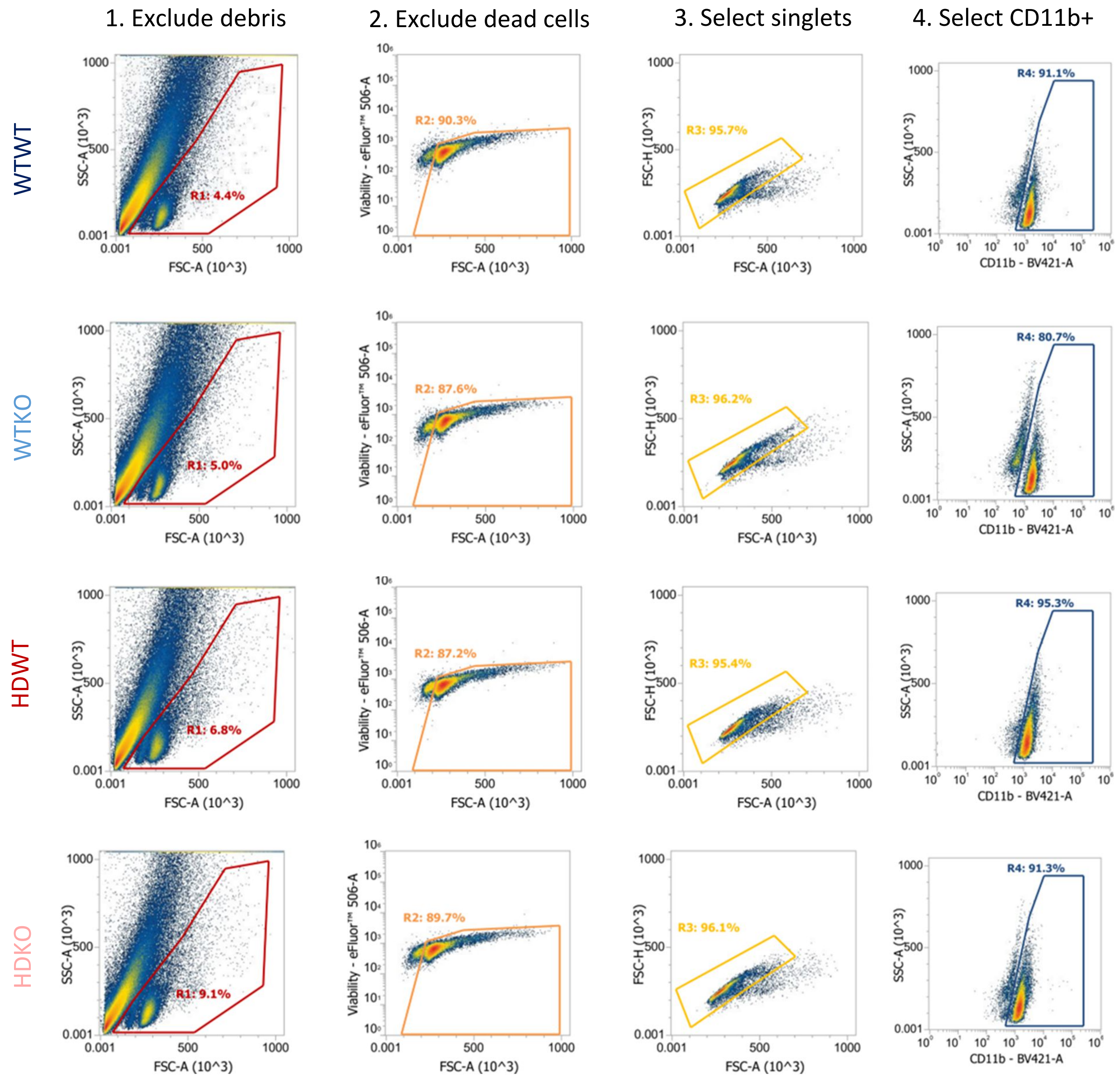

### Supplementary Figure 13

#### Analysis of homeostatic and activated microglial markers

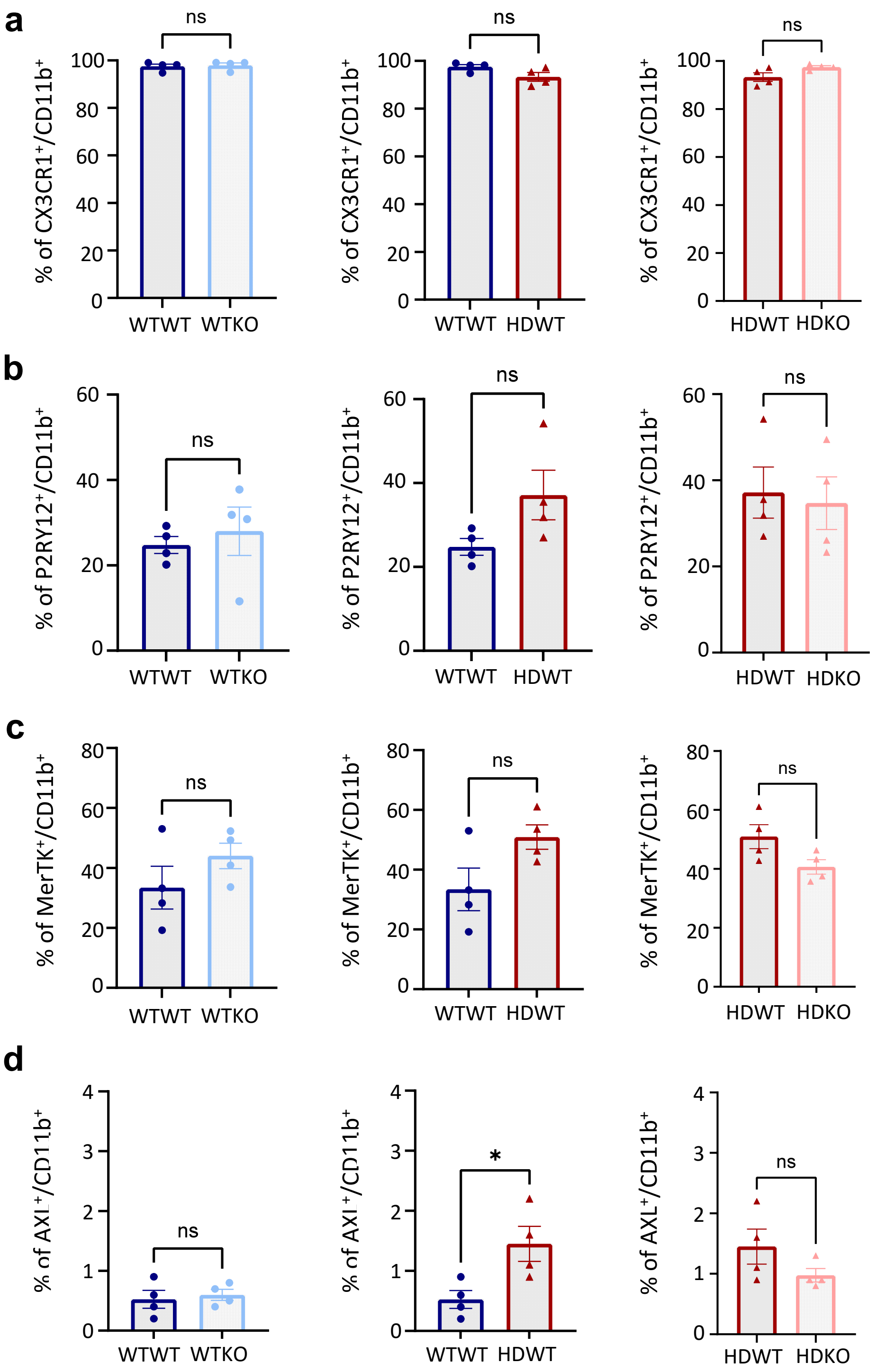

### Supplementary Figure 14

#### MS$^2$ of DMA-labeled disialylated N-glycans

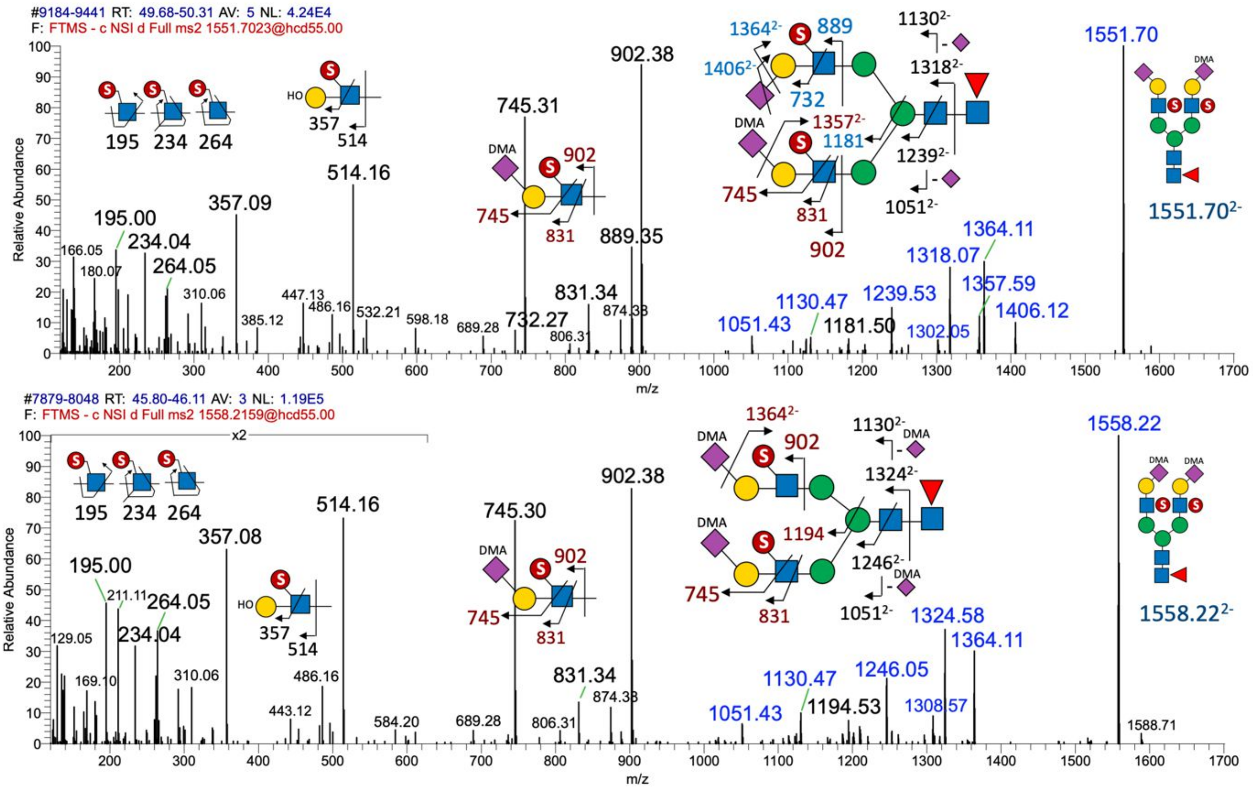

### Supplementary Figure 15

#### Gating strategy for cell type-specific α2-6-sialylated 6-sulfo LacNAc

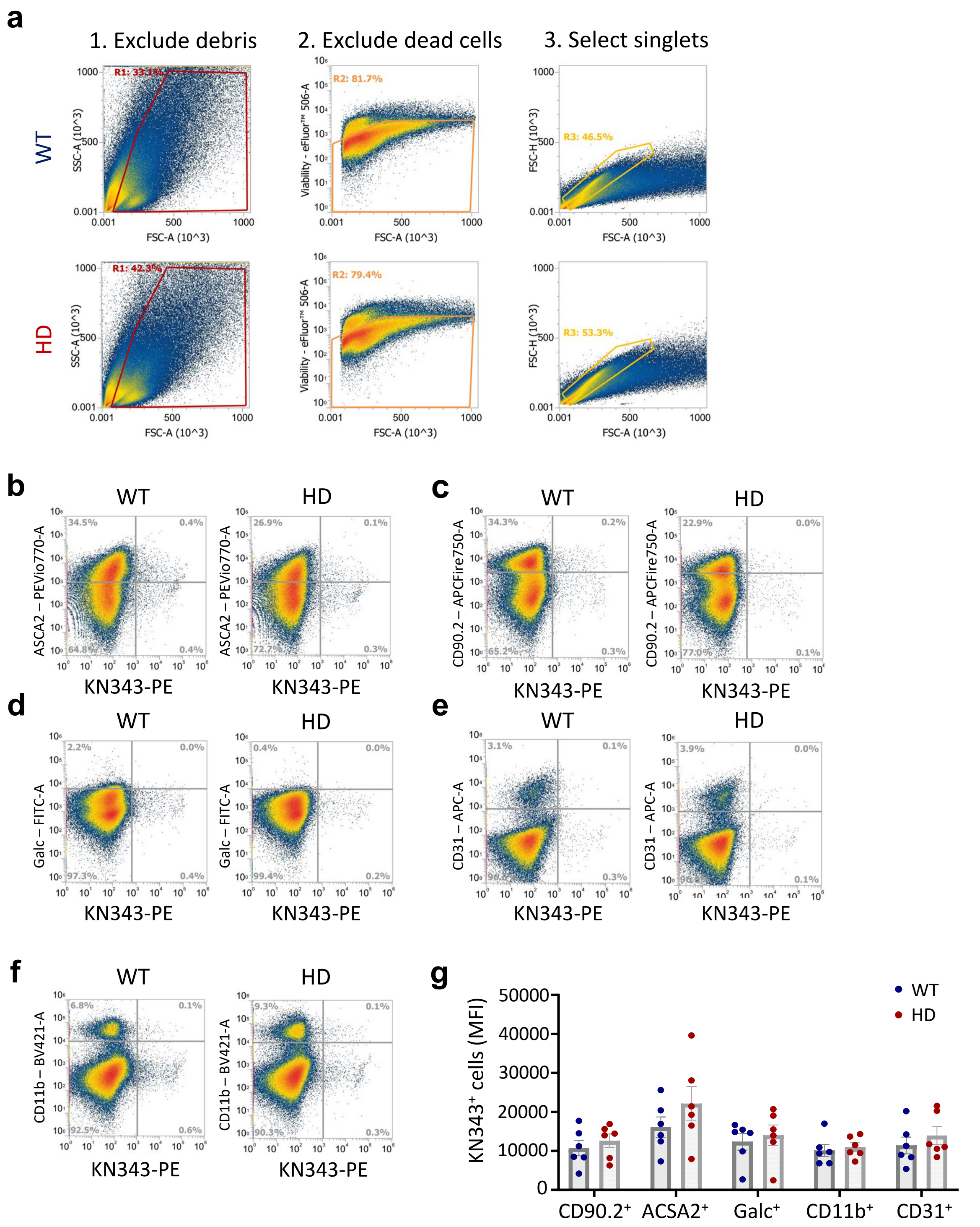

### Supplementary Figure 16

#### Expression of *St6gal1* and *Chst2* in single-nuclei RNA-seq data

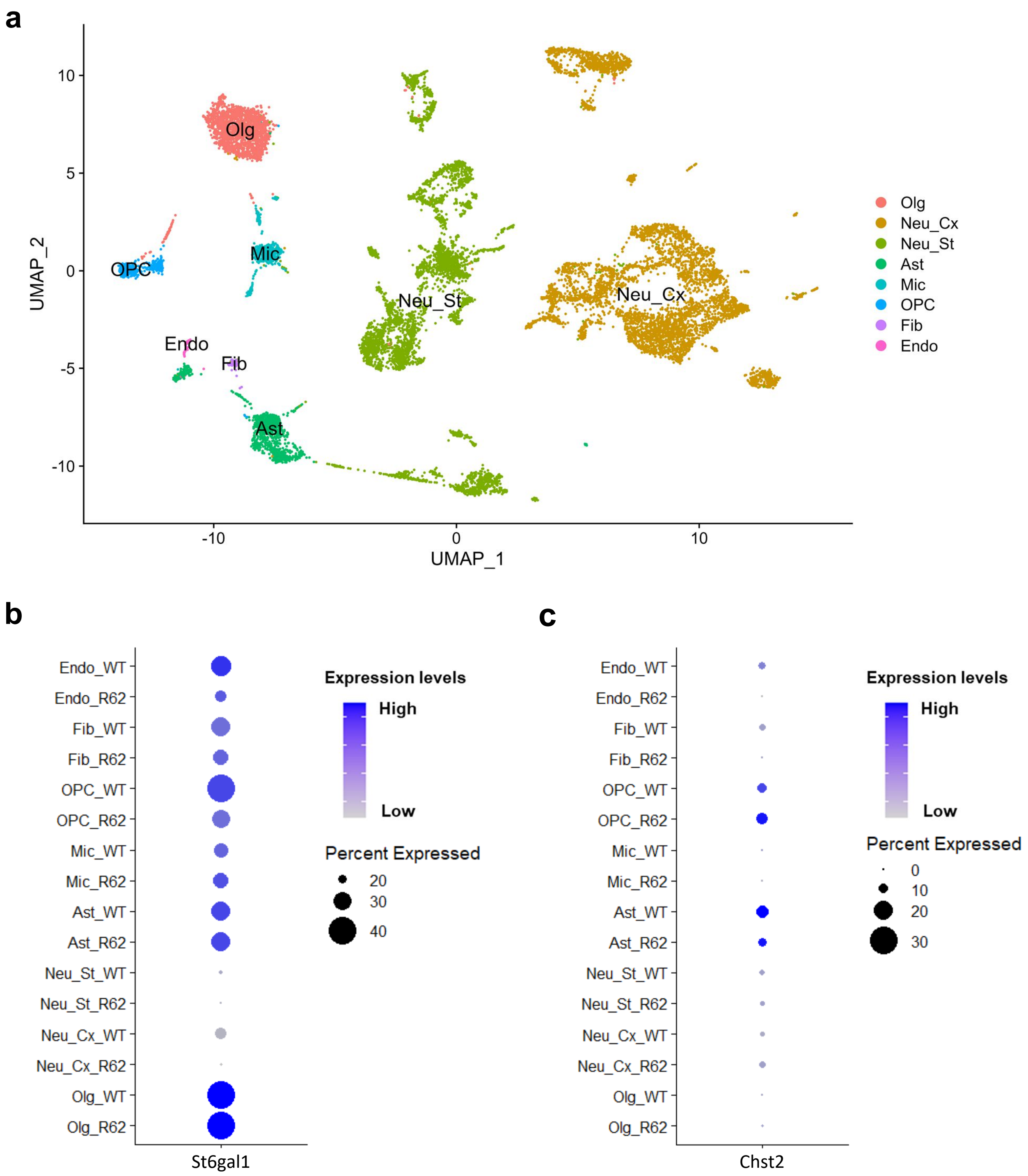

### Supplementary Figure 17

#### Quality control metric and cell annotation

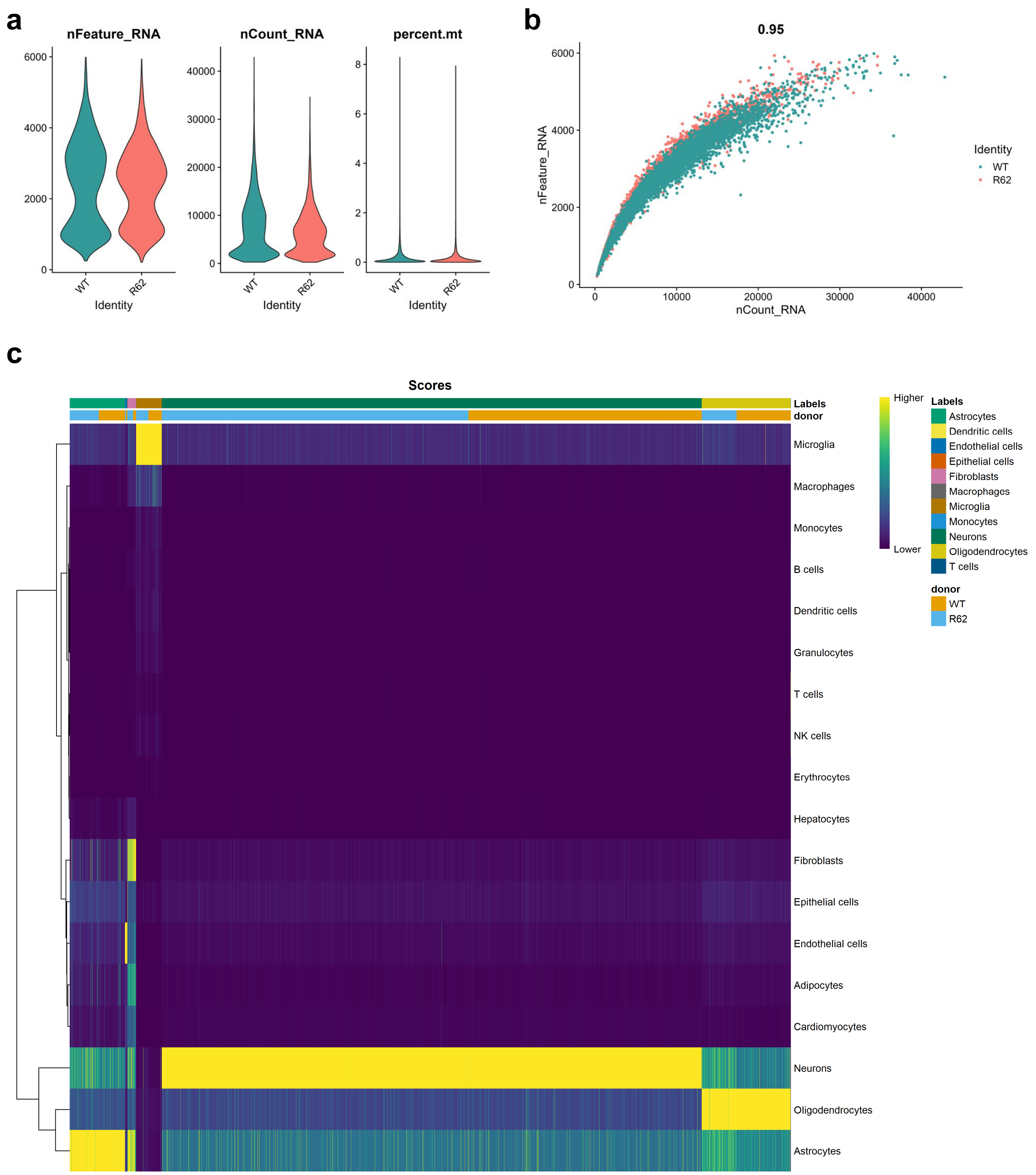

### Supplementary Figure 18

#### Cell type identification using marker genes

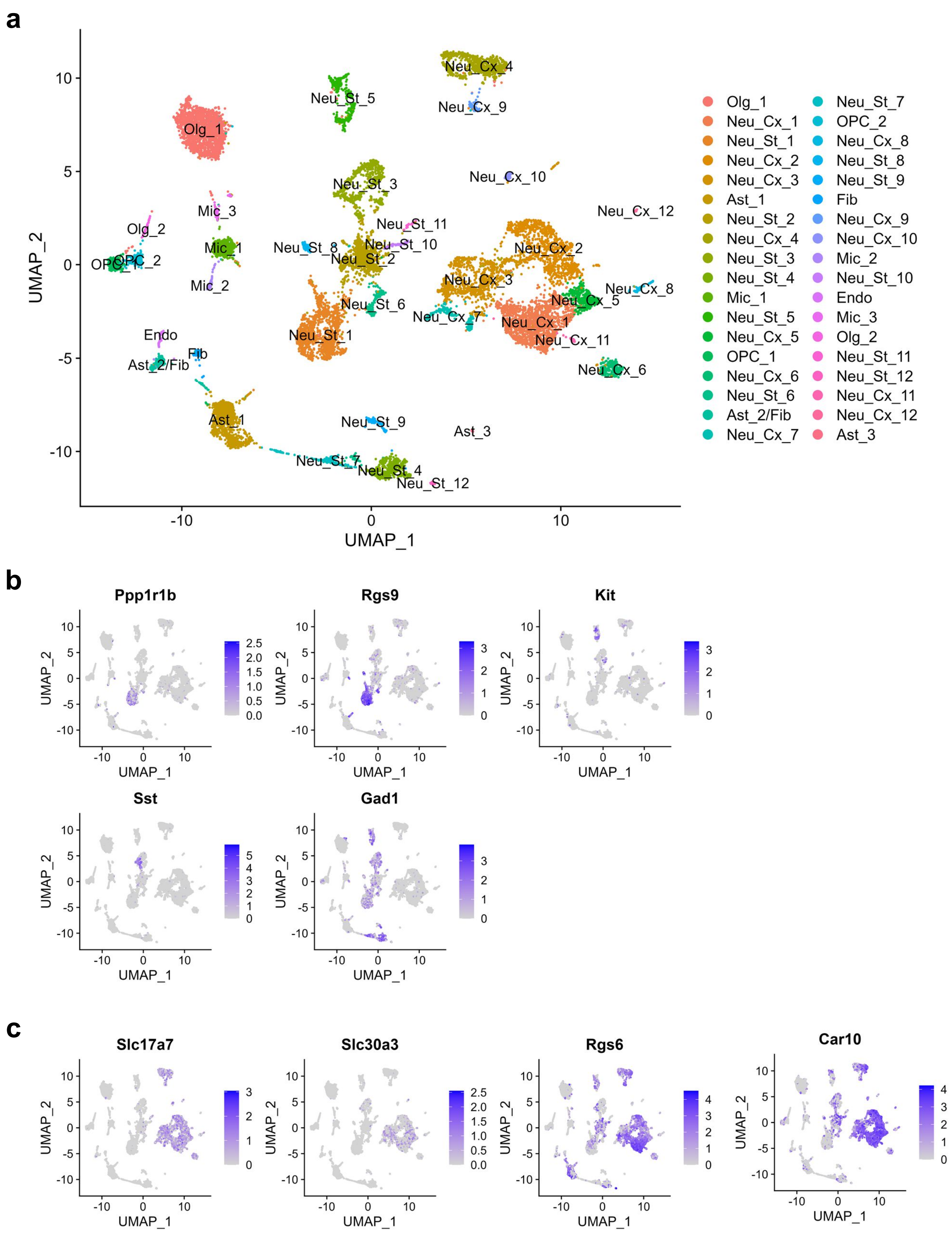

### Supplementary Figure 19

#### Co-culture of BV2-CD22 cells with C8-D1A–CHST2 cells

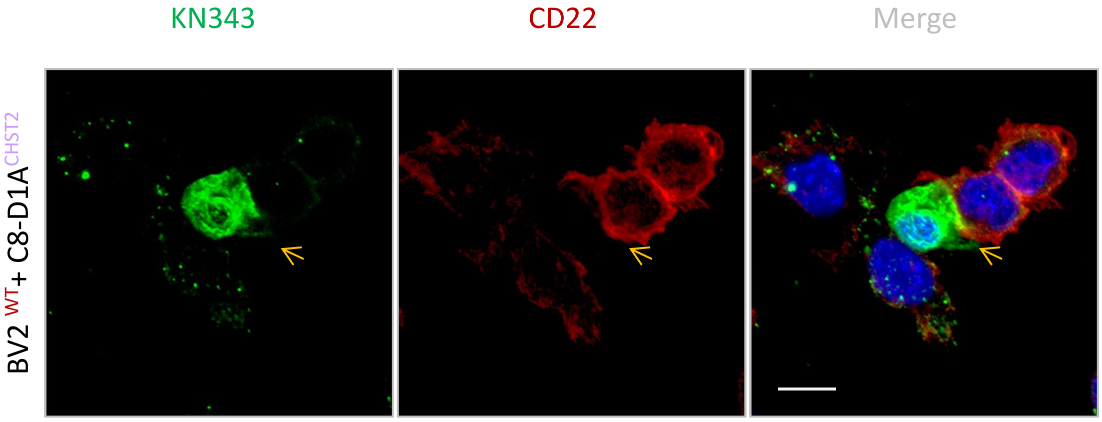

#### Supplementary Figure 20

##### Presence of mHTT does not alter CD22 transcript levels

**a**

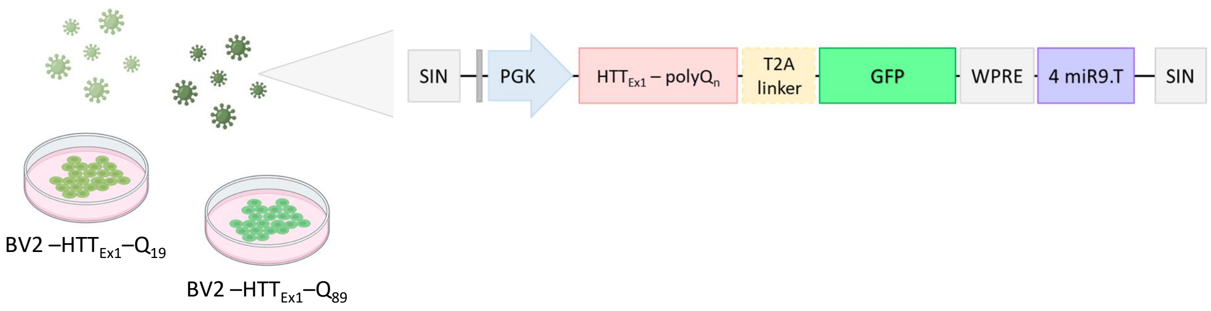


**b**

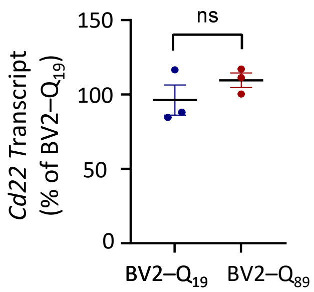

### Supplementary Figure 21

#### Exposure to oxidative stress does not alter Siglec-E transcript levels

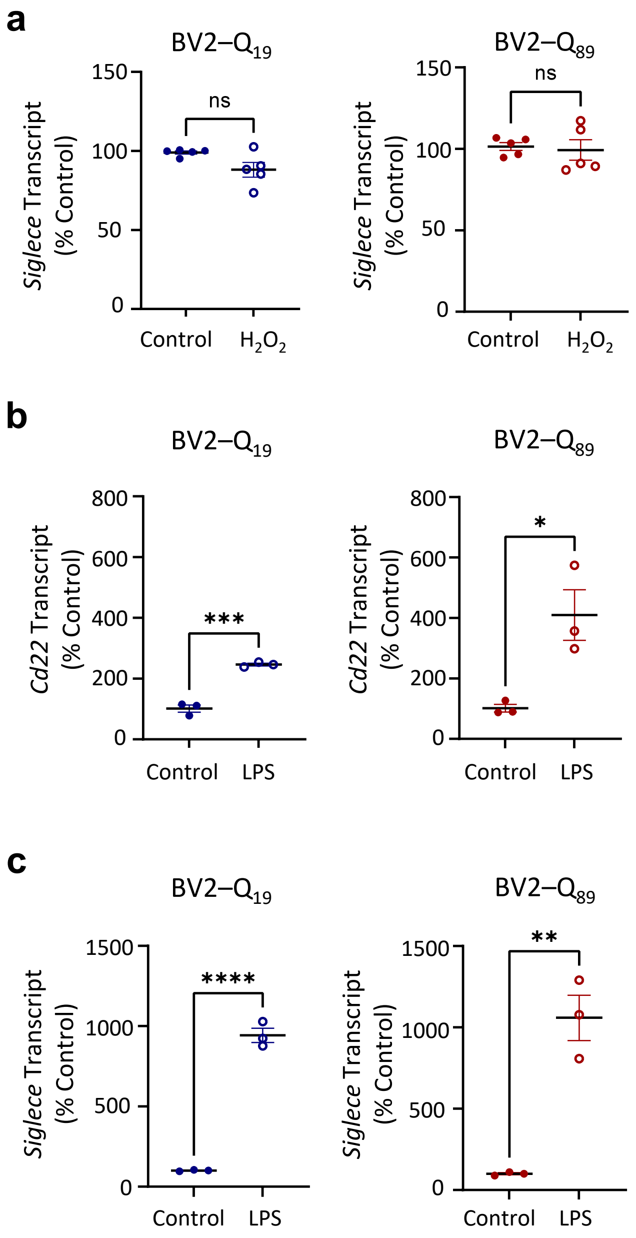

### Supplementary Figure 22

#### NAC treatment has no effect in WT mice

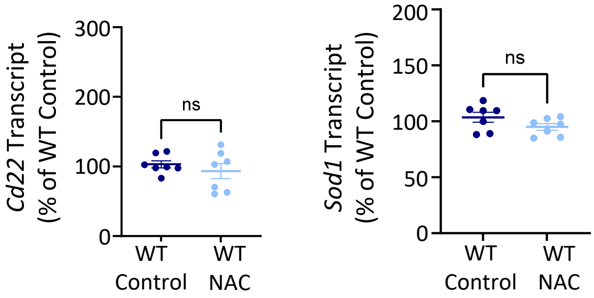

### Supplementary Figure 23

#### Presence of mHTT does not alter *St6gal1* and Chst2 expression in C8-D1A astrocyte

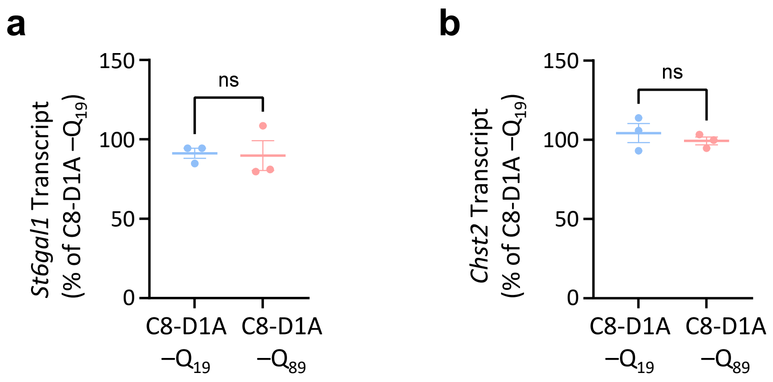

### Supplementary Figure 24

#### Co-immunoprecipitation analysis of CD22 and *Clec7a*-V5

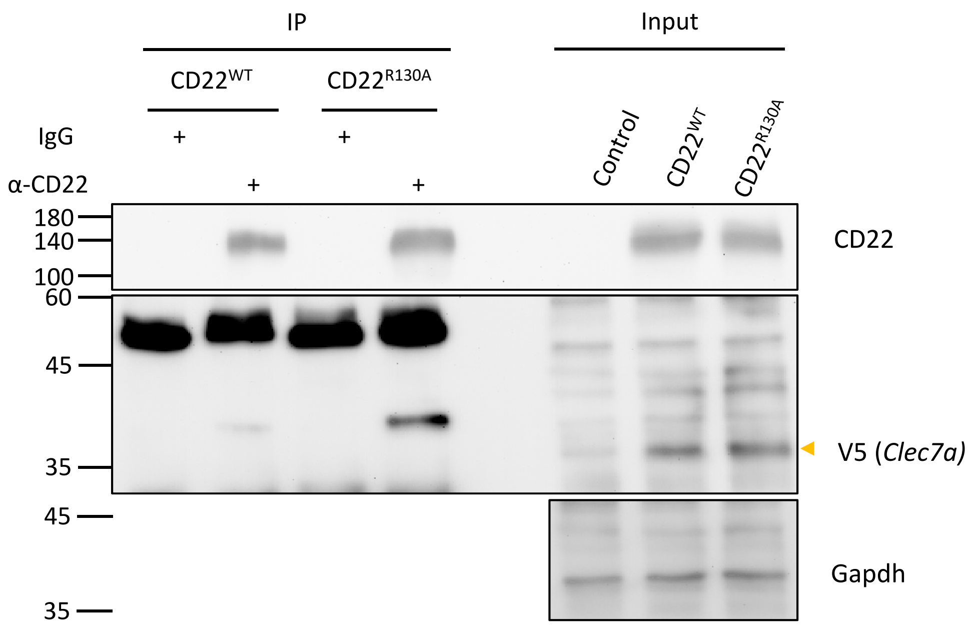

### Supplementary Figure 25

#### CD22 is significantly up-regulated in mouse model of AD

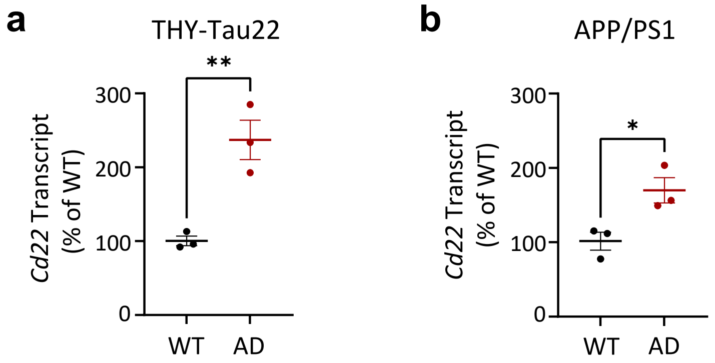

### Supplementary Table 3

| Gene | Primers |
|---|---|
| *SIGLEC1* | **F** 5'- CACTCTTCTTCCAGGTCCGAG -3' |
| | **R** 5'- ACCAACGATATGAGGTCCCCT -3' |
| *CD22* | **F** 5'- GCACCCTGAAACCCTCTACG -3' |
| | **R** 5'- ATCAAACTTCGAGGTGTTCTTGT -3' |
| *CD33* | **F** 5'- GGAGATGGCTCAGGGAAACAA -3' |
| | **R** 5'- GTAGGGTGGGTGTCATTCCTG -3' |
| *MAG* | **F** 5'- TACCCAGACACGCAGGAAAAA -3' |
| | **R** 5'- CCCAGGTCCGAGTGAGAATAG -3' |
| *SIGLEC5* | **F** 5'- CCTGTCAGATGAAACGCCAAG -3' |
| | **R** 5'- AGGACCGGAAGGTATGAGGTG -3' |
| *SIGLEC6* | **F** 5'- ACGACCCAGACGAAGAAGTG -3' |
| | **R** 5'- GCACAGAGAGCTTGGAAGATG -3' |
| *SIGLEC7* | **F** 5'- CTGGTCTTCCTCTCCTTCTGTG -3' |
| | **R** 5'- GCATCCTTCATGCCTATGTCTCC -3' |
| *SIGLEC8* | **F** 5'- TGTGTCCTACCCTCCTTGGAA -3' |
| | **R** 5’- GGCCCTCAAGGACTGAAAGAG -3’ |
| *SIGLEC9* | **F** 5’- AGACCGTCCATCTCAACGTGT -3’ |
| | **R** 5’- TGGGAGTGACAGAGATGAGCC -3’ |
| *SIGLEC10* | **F** 5’- TCATCTCAACGGCATTCTCCA -3’ |
| | **R** 5’- GGCTTTCTGATTCCGCTTCTG -3’ |
| *SIGLEC11* | **F** 5’- TGCTGCTTATGGCTACTGGTT -3’ |
| | **R** 5’- AACGCATTGCTCAGGAAACTA -3’ |
| *SIGLEC12* | **F** 5’- GGCTGTCCGACTCAACATATC -3’ |
| | **R** 5’- GTCGACAGCACAGACAAGGTG -3’ |
| *SIGLEC14* | **F** 5’- TCCAGCTCAATGTCTCCTATGC -3’ |
| | **R** 5’- ACCGACATGCCATTGCTCA -3’ |
| *SIGLEC15* | **F** 5’- CGCGGATCGTCAACATCTC -3’ |
| | **R** 5’- GTTCGGCGGTCACTAGGTG -3’ |
| *SIGLEC16* | **F** 5’- CGTGCACTGCAAATCAGG -3’ |
| | **R** 5’- CTCCTGCAAGACCTCACTCTG -3’ |
| *18s* | **F** 5’- GTAACCCGTTGAACCCCATT -3’ |
| | **R** 5’- CCATCCAATCGGTAGTAGCG -3’ |

### Supplementary Table 4

| Gene | Primers |
|---|---|
| *Cd22* | **F** 5'- CATCCCTAAAGTCTCCCCGTG -3' |
| | **R** 5'- CCAGGATTGGTGTGAAGCTCT -3' |
| *Cd22* (Exon11-specific) | **F** 5'- CTTGGGACTAGGCTTCTGCCTG -3' |
| | **R** 5'- TTTTCCTGAAGCCCCTGCTGGC -3' |
| *Siglece* | **F** 5'- CTCAGAATCCCCTGGTTGAA -3' |
| | **R** 5'- CAGGTTCATGGGCTTCATCT -3' |
| *St6gal1* | **F** 5'- ATGGGGCACCTACAGACAAC -3' |
| | **R** 5'- GCTTCTGATACCACTGCGGA -3' |
| *Chst2* | **F** 5'- CCGCTCGGGATGAAGGTATTT -3' |
| | **R** 5'- CCACTTGTAGTCCAAGAGGTTGA -3' |
| *Hif1a* | **F** 5'- AGAACTAAACACACAGCGGAGC -3' |
| | **R** 5'- ACGTCATGGGTGGTTTCTTGTA -3' |
| *Nfe2l2* | **F** 5'- CAGCATAGAGCAGGACATGGAG -3' |
| | **R** 5'- GAACAGCGGTAGTATCAGCCAG -3' |
| *Sod1* | **F** 5'- CGTCCATCAGTATGGGGACAA -3' |
| | **R** 5'- CAGGTCTCCAACATGCCTCTC -3' |
| *Ptgs2* | **F** 5'- ATCATAAGCGAGGACCTGGG -3' |
| | **R** 5'- TGAGTGTCTTTGACTGTGGGG -3' |
| *Il1b* | **F** 5'- TGAAGTTGACGGACCCCAAAA -3' |
| | **R** 5'- ATGAGTGATACTGCCTGCCTG -3' |
| *Il6* | **F** 5'- GAGGATACCACTCCCAACAGACC -3' |
| | **R** 5'- AAGTGCATCATCGTTGTTCATAC -3' |
| *Il10* | **F** 5'- ATTCCCTGGGTGAGAAGCTGA -3' |
| | **R** 5'- ACACCTTGGTCTTGGAGCTTA -3' |
| *Cd68* | **F** 5'- ACATGGCGGTGGAATACAAT -3' |
| | **R** 5'- TCAAGGTGAACAGCTGGAGA -3' |
| *Gapdh* | **F** 5'- TGACATCAAGAAGGTGGTGAAG -3' |
| | **R** 5'- AGAGTGGGAGTTGCTGTTGAAG -3' |

### Supplementary Table 5

| Target | Metal | Element | Vendor | Catalog |
|---|---|---|---|---|
| BC1 | 102 | Pd | | |
| BC2 | 104 | Pd | | |
| BC3 | 105 | Pd | | |
| BC4 | 106 | Pd | | |
| BC5 | 108 | Pd | | |
| BC6 | 110 | Pd | | |
| CD45 | 89 | Y | Biolegend | 103102 |
| GAPDH | 111 | Cd | Invitrogen | AM4300 |
| Ki67 | 112 | Cd | BD | 556003 |
| CCR2 | 113 | In | Biolegend | 160102 |
| P2RY12 | 114 | Cd | Biolegend | 848002 |
| H3K9ac | 115 | In | Cell signaling | 9649 |
| Ly6G | 139 | La | Biolegend | 127602 |
| CD11b | 140 | Ce | Biolegend | 101202 |
| F4/80 | 141 | Pr | Biolegend | 123102 |
| CD39 | 142 | Nd | Fluidigm | 3142005B |
| CD11c | 143 | Nd | BD | 553799 |
| p4EBP1 | 144 | Nd | Cell signaling | 2855 |
| CD4 | 145 | Nd | BD | 553043 |
| iNOS | 146 | Nd | eBioscience | 14-5920-82 |
| pH2AX | 147 | Sm | Sigma-Aldrich | 05-636 |
| CD49e | 148 | Nd | Biolegend | 103801 |
| CD49d | 149 | Sm | Biolegend | 103709 |
| CD27 | 150 | Nd | Biolegend | 124202 |
| Ly6C | 151 | Eu | Biolegend | 128002 |
| CD3 | 152 | Sm | Fluidigm | 3152004B |
| PD-L1 | 153 | Eu | Biolegend | 124302 |
| Siglec-H | 154 | Sm | Biolegend | 129602 |
| Siglec E | 155 | Gd | R&D | MAB5806 |
| CD90.2 | 156 | Gd | Fluidigm | 3156006B |
| CD40 | 157 | Gd | Biolegend | 102902 |
| CXCR3 | 158 | Gd | eBioscience | 16-1831-81 |
| Osteopontin (OPN) | 159 | Tb | R&D | AF808 |
| CD206 | 160 | Gd | Biolegend | 141702 |
| TMEM119 | 161 | Dy | Abcam | ab220249 |
| TIM-3 | 162 | Dy | Biolegend | 119702 |
| CD80 | 163 | Dy | Biolegend | 104735 |
| TIM4 | 164 | Dy | Merck Millipore | MABC958 |
| H3K27me3 | 165 | Ho | GeneTex | GTX50901 |
| MerTK | 166 | Er | R&D | AF591 |
| Siglec-3(CD33) | 167 | Er | R&D | AF2220 |
| Axl | 168 | Er | R&D | AF854 |
| TREM-2 | 169 | Tm | Bio-rad | MCA4772GA |
| Siglec G | 170 | Er | Biolegend | 163302 |
| Siglec-2(CD22) | 171 | Yb | R&D | MAB2296 |
| Siglec-15 | 172 | Yb | (in-house) | (in-house) |
| CD19 | 173 | Yb | Biolegend | 115502 |
| SiglecF | 174 | Yb | BD | 552125 |
| CD44 | 175 | Lu | BD | 553131 |
| Siglec-1(CD169) | 176 | Yb | Biolegend | 142402 |
| MHCII | 209 | Bi | Biolegend | 107602 |
| Cell-ID | 191 | Ir | Fluidigm | 201192B |
| Cell-ID | 193 | Ir | Fluidigm | 201192B |
| Cisplatin | 198 | Pt | Sigma-Aldrich | P4394 |

### Supplementary Figure Legends

**Supplementary Figure 1. CD22 expression across brain cell types in mouse and human**

CD22 expression in various cell types in the mouse (**a**) and human (**b**) brain. Data were retrieved from single-cell RNA-seq dataset from Zhang et al [4] and Zhang et al [5].

**Supplementary Figure 2. Traditional flow cytometry confirms expansion of CD22-immunoreactive microglia in HD mice**

Traditional flow cytometry analysis using adult primary microglia isolated from WT and HD brains at 13 weeks of age confirmed the expansion of CD22-immunoreactive microglia in HD mice. Data are presented as mean ± SEM (n = 4 animals per group). *$p < 0.05$ by unpaired *t*-test.

**Supplementary Figure 3. Gating hierarchy of FlowSOM**

Cells are first gated on live, singlet, and $CD90.2^{-}$ populations for data analysis.

**Supplementary Figure 4. First UMAP**

An initial UMAP of $C90.2^{-}$ population was conducted using 27 immune markers: AXL, CCR2, CD11b, CD11c, CD19, CD27, CD3, CD39, CD4, CD40, CD44, CD45, CD49d, CD49e, CD80, CXCR3, F4/80, Ly6C, Ly6G, MerTK, MHCII, P2RY12, PD-L1, TIM-3, TIM-4, TMEM119, and TREM2. The microglia population was subsequently isolated based on $CD39^{+}$ and $P2RY12^{+}$ expression, and a secondary UMAP analysis was performed to focus on Siglec expression.

**Supplementary Figure 5. Population percentages of FlowSOM-identified metacluster**

Population percentages of the identified metaclusters were determined using FlowSOM analysis. Statistical analysis revealed no significant differences between groups. Data are presented as mean ± SD (n = 3 independent experiments).

**Supplementary Figure 6. Second UMAP**

Relative expression levels of individual Siglec markers in the microglia population, as shown in the secondary UMAP.

**Supplementary Figure 7. CD22 does not affect the inflammatory or oxidative stress response of microglia**

(**a**) Stable BV2 cell lines with stable CD22 (BV2-CD22) expression and vector control (BV2-miR9T) were generated via lentivirus transduction. CD22 was fused to a GFP reporter through a T2A linker to enable independent expression of CD22 and GFP, thereby preventing the GFP reporter from interfering with the folding or function of the CD22. Over-expression of CD22 does not affect the inflammatory response of BV2 microglia upon LPS stimulation (100 ng/mL for 24 hours), as indicated by qPCR analysis of pro-inflammatory markers (*Il1b* and *Il6*; **b**) and an anti-inflammatory marker (*Il10*; **c**). Similarly, over-expression of CD22 does not affect the oxidative stress response of BV2 microglia upon oxidative stress stimulation (50µM for 24 hours), as indicated by qPCR analysis of oxidative stress markers (*Hif1a* and *Nfe2l2*; **d**) and an oxidative stress-induced inflammation marker (*Ptgs2*; **e**). Data are presented as mean ± SEM (n = 6). *$p < 0.05$ and ****$p < 0.0001$ by two-way ANOVA.

**Supplementary Figure 8. CD22 is co-localized with *Clec7a*-V5**

HEK293T cells were transfected with Dectin-1 (*Clec7a*-V5), along with $CD22^{WT}$ or $CD22^{R130A}$ for 48 hours. Cells were then plated on coverslips and allowed to attach overnight before fixation with 4% paraformaldehyde. Immunofluorescence staining was performed using an anti-V5 antibody (red) to detect Dectin-1 (*Clec7a*) and anti-CD22 antibody (green) to detect CD22. Immunofluorescence staining revealed complete co-localization of Dectin-1 (*Clec7a*) and CD22, as indicated by the overlapping red and green fluorescence.

**Supplementary Figure 9. Knockout of CD22 did not affect body weight, limb clasping or life span**

Knockout of CD22 did not alter the body weight (**a**), limb clasping response (**b**) or life span (**c**) of the HD mice. Data are presented as mean ± SEM (n = 9-12 animals per group). *Specific comparison between WT and HD mice in each condition; #Specific comparison between CD22-WT and CD22-KO in each condition. ****$p < 0.0001$ by two-way ANOVA.

**Supplementary Figure 10. *Cd68* is significantly down-regulated in the HD striatum**

Transcript levels of *Cd68,* a lysosomal marker associated with phagocytic activity, were assessed in the striatum of HD mice at 13 weeks of age using qPCR analysis. Transcript levels of *Cd68* were significantly down-regulated in the striatum of HD mice at 13 weeks of age. Data are presented as mean ± SEM (n = 4 animals per group). *$p < 0.05$ by unpaired *t*-test.

**Supplementary Figure 11. Knockout of CD22 does not affect the volume of the whole brain, cortex, or hippocampus**

Quantification of the whole brain (**a**), cortical (**b**), and hippocampal (**c**) volumes showed that HD mice had reduced whole brain, cortical and hippocampal volumes compared to WT mice, but were not affected by CD22 knockout. Nevertheless, CD22 knockout does not affect the whole brain (**d**), cortical (**e**) and hippocampal (**f**) volumes of the WT mice. Data are presented as the means ± SEM (n = 5-8 animals per group). ****$p < 0.0001$ by unpaired *t*-test.

**Supplementary Figure 12. Flow cytometry gating hierarchy used to identify CD11b$^+$ microglia**

Adult microglia were isolated from the cortex and striatum of 13 weeks of age mice by mechanical dissociation. CD11b$^+$ cells were gated, and homeostatic and activated microglia markers were analyzed across WT/CD22$^{WT}$, WT/CD22$^{KO}$, HD/CD22$^{WT}$, and HD/CD22$^{KO}$ mice.

**Supplementary Figure 13. Analysis of homeostatic and activated microglial markers**

Markers associated with homeostatic status, including CX3CR1 (**a**), P2RY12 (**b**), and MerTK (**c**), were not significantly altered in HD mice and were unaffected by CD22 knockout. The activation-associated marker AXL (**d**) was significantly upregulated in HD mice, and CD22 knockout did not alter the proportion of AXL$^+$ microglia. Data are presented as mean ± SEM (n = 4 independent replicates). *$p < 0.05$ by unpaired *t*-test.

**Supplementary Figure 14. MS$^2$ of DMA-labeled disialylated N-glycans**

MS$^2$ of DMA-derivatized and permethylated disialylated biantennary N-glycans. Terminal 2-6-linked sialic acid is marked with DMA on the diamond symbol representing NeuAc. The B ion at *m/z* 902, along with an E and $^{3,5}$A ion at *m/z* 831 and 745, respectively, identify the presence of a terminal 2-6-sialylated, sulfated Hex-HexNAc glycotope, whereas the diagnostic trio of ions at *m/z* 195, 234, and 264 unambiguously locate the sulfate on the 6-position of an internal HexNAc. The corresponding non-DMA-tagged and hence 2-3-sialylated sulfated Hex-HexNAc afforded instead the B ion and $^{3,5}$A ions at *m/z* 889 and 732, respectively, which were not produced if both sialylation were 2-6-linked. The doubly charged parent ions and fragment ions resulting from losses of non-sulfated moiety were annotated in blue, whereas singly charged fragment ions carrying a single sulfate were annotated in black.

**Supplementary Figure 15. Gating strategy for cell type-specific α2-6-sialylated 6-sulfo LacNAc**

Brain cells were isolated from the cortex and striatum of 13 weeks of age mice by enzymatic dissociation. (**a**) Analysis was gated on live and singlet cells. Gating strategies used identify ACSA2$^+$ astrocytes (**b**), CD90.2$^+$ neurons (**c**), Galc$^+$ oligodendrocytes (**d**), CD31$^+$ endothelial cells (**e**) and CD11b$^+$ microglia (**f**) from isolated brain cells of WT and HD mice, followed by detection of KN343$^+$signal for α2-6-sialylated 6-sulfo LacNAc. (**g**) MFI (mean fluorescence intensity) of KN343$^+$ cells across cell types from the cytometry analysis. Data are presented as mean ± SEM (n = 6 independent experiments).

**Supplementary Figure 16. Expression of *St6gal1* and *Chst2* in single-nucleus RNA-seq data**

Brain nuclei were isolated from the cortex and striatum of 13 weeks of age mice for nucleus RNA sequencing. (**a**) UMAP plot of single-nuclei transcriptomes showing major cell populations, including oligodendrocytes (Olg), cortical neurons (Neu_Cx), striatal neurons (Neu_St), astrocytes (Ast), microglia

(Mic), oligodendrocyte precursor cells (OPC), fibroblasts (Fib), and endothelial cells (Endo). Dot plot showing expression levels and the percentage of cells expressing *St6gal1* (**b**) and *Chst2* (**c**) across cell types and genotypes, with notable differences between WT and HD mice that suggest potential alterations in sialylation pathways in HD.

**Supplementary Figure 17. Quality control metric and cell annotation**.

(**a**) Violin plots displaying the distribution of detected genes (*nFeature_RNA*), total UMI counts (*nCount_RNA*), and mitochondrial transcript percentages (*percent.mt*) for each dataset. (**b**) Scatter plot illustrating the relationship between total UMI counts and the number of detected genes across all cells (Pearson correlation coefficient = 0.95). (**c**) Heatmap presenting cell type annotations based on SingleR classification.

**Supplementary Figure 18. Cell type identification using marker genes**

(**a**) UMAP of all single nuclei showing cluster distribution. Initial clustering analysis revealed a total of 36 cell clusters. (**b**) Feature plots showing the expression of marker genes (*Ppp1r1b*, *Rgs9*, *Kit*, *Sst*, *Gad1*) used to identify striatal neurons. (**c**) Feature plots showing the expression of marker genes (*Slc17a7*, *Slc30a3*, *Rgs6*, *Car10*) used to identify cortical neurons.

**Supplementary Figure 19. Co-culture of BV2-CD22 cells with C8-D1A–CHST2 cells**

CD22-expressing BV2 microglia were co-cultured with CHST2-expressing C8-D1A astrocytes on coverslips for 24 hours. Immunofluorescence staining was performed using KN343 (green) to detect α2-6-sialylated 6-sulfo LacNAc, and anti-CD22 antibody (red) to detect CD22. Nuclei are stained with Hoechst (blue). No clustering of CD22 was observed when a BV2-$CD22^{WT}$ cell was located adjacent to the C8-D1A–CHST2 cell (yellow arrow).

**Supplementary Figure 20. Presence of mHTT does not alter CD22 transcript levels**

(**a**) Stable BV2 cell lines with consistent $HTT_{Ex1}$-$Q_{19}$ (BV2-$Q_{19}$; wild-type) and $HTT_{Ex1}$-$Q_{89}$ (BV2-$Q_{89}$; mutant) expression were generated by lentivirus transduction. $HTT_{Ex1}$-$Q_{19}$ or $HTT_{Ex1}$-poly$Q_{89}$ were fused to a GFP-reporter through a T2A-linker. The T2A linker will induce ribosomal skipping during translation and allowing independent expression of $HTT_{Ex1}$-$Q_{19}$ or $HTT_{Ex1}$-$Q_{89}$ and GFP, thereby preventing the GFP reporter from interfering with the folding or properties of the $HTT_{Ex1}$-$Q_{19}$ or $HTT_{Ex1}$-$Q_{89.}$ (**b**) In BV2-$Q_{89}$ cells expressing mutant HTT, CD22 transcript levels remain unchanged compared to BV2-$Q_{19}$ cells expressing wild-type HTT, as assessed by qPCR analysis. Data are presented as mean ± SEM (n = 3).

**Supplementary Figure 21. Exposure to oxidative stress does not alter Siglec-E transcript levels**

Stable BV2 cell lines with consistent $HTT_{Ex1}$-$Q_{19}$ (BV2-$Q_{19}$; wild-type) and $HTT_{Ex1}$-$Q_{89}$ (BV2-$Q_{89}$; mutant) expression were exposed to HD-related stimuli, including those that induce inflammatory responses and oxidative stress. (**a**) Exposure to oxidative stress does not alter Siglec-E transcript levels in either BV2-$Q_{19}$ or BV2-$Q_{89}$ cells. Data are presented as mean ± SEM (n = 5). Exposure to pro-inflammatory stimulant (100ng/mL LPS for 24 hours) induced up-regulation of both CD22 (**b**) and Siglec-E (**c**) in both BV2-$Q_{19}$ and BV2-$Q_{89}$ cells. This suggests that neuroinflammation triggers a broad inflammatory response, whereas oxidative stress elicits a more selective pathway that primarily affects CD22. Data are presented as mean ± SEM (n = 3). *$p < 0.05$, **$p < 0.01$, ***$p < 0.001$ and ****$p < 0.0001$ by unpaired *t*-test.

**Supplementary Figure 22. NAC treatment has no effect in WT mice**

Wild-type (WT) and HD mice were fed with NAC in the drinking water from 7 to 13 weeks of age to mitigate oxidative stress accumulation, and brain tissues were collected at 13 weeks of age. NAC treatment does not affect the levels of *Cd22* and *Sod1* in WT mice. Data are presented as mean ± SEM (n = 7 animals per group).

**Supplementary Figure 23. Presence of mHTT does not alter *St6gal1* and *Chst2* expression in C8-D1A astrocyte**

C8-D1A cell line with stable $HTT_{Ex1}$–$Q_{19}$ (wild-type; designated C8-D1A–$Q_{19}$) and $HTT_{Ex1}$–$Q_{89}$ (mutant; designated C8-D1A–$Q_{89}$) were generated using a lentiviral system. In C8-D1A–$Q_{89}$ cells expressing mutant HTT, the transcript levels of *St6gal1* (**a**) and *Chst2* (**b**) remain unchanged compared to C8-D1A–$Q_{19}$ cells expressing wild-type HTT, as assessed by qPCR analysis. Data are presented as mean ± SEM (n = 3).

**Supplementary Figure 24. Co-immunoprecipitation analysis of *Clec7a*-V5 and CD22**

HEK293T cells were transfected with the indicated construct for 48 hours and subjected to co-immunoprecipitation assays. The indicated lysates (2 mg) were incubated with anti-CD22 antibody for 90 minutes at 4°C to allow complex formation. The immunoprecipitated proteins were released from the immunocomplexes and analyzed by Western blot analysis. Dectin-1 (*Clec7a*-V5) was detected using an anti-V5 antibody, as indicated by the yellow arrowhead. No direct interaction between $CD22^{WT}$ or $CD22^{R130A}$ and Dectin-1 was detected.

**Supplementary Figure 25. CD22 is significantly up-regulated in the mouse models of AD**

qPCR analysis using cDNA prepared from the hippocampus of AD mice (n=3 per genotype) at 13 months-of-age (late stage). CD22 is significantly up-regulated in the hippocampus of both APP/PS1 (**a**) and THY-Tau22 (**b**) mice. Data are presented as mean ± SEM (n = 3 animals per group). *$p < 0.05$ and *$p < 0.01$ by unpaired *t*-test.

#### Supplementary Table 6

Excel file containing a list of marker genes per annotated cell cluster.

#### Supplementary Table 7

Excel file containing differentially expressed genes (DEGs) between WT and HD mice across annotated cell types in snRNA-seq data ($log_2FC \geq 0.25$).
